# Leveraging the Genetic Correlation between Traits Improves the Detection of Epistasis in Genome-wide Association Studies

**DOI:** 10.1101/2022.11.30.518547

**Authors:** Julian Stamp, Alan DenAdel, Daniel Weinreich, Lorin Crawford

## Abstract

Epistasis, commonly defined as the interaction between genetic loci, is known to play an important role in the phenotypic variation of complex traits. As a result, many statistical methods have been developed to identify genetic variants that are involved in epistasis, and nearly all of these approaches carry out this task by focusing on analyzing one trait at a time. Previous studies have shown that jointly modeling multiple phenotypes can often dramatically increase statistical power for association mapping. In this study, we present the “multivariate MArginal ePIstasis Test” (mvMAPIT) — a multi-outcome generalization of a recently proposed epistatic detection method which seeks to detect *marginal epistasis* or the combined pairwise interaction effects between a given variant and all other variants. By searching for marginal epistatic effects, one can identify genetic variants that are involved in epistasis without the need to identify the exact partners with which the variants interact — thus, potentially alleviating much of the statistical and computational burden associated with conventional explicit search-based methods. Our proposed mvMAPIT builds upon this strategy by taking advantage of correlation structure between traits to improve the identification of variants involved in epistasis. We formulate mvMAPIT as a multivariate linear mixed model and develop a multi-trait variance component estimation algorithm for efficient parameter inference and *P*-value computation. Together with reasonable model approximations, our proposed approach is scalable to moderately sized GWA studies. With simulations, we illustrate the benefits of mvMAPIT over univariate (or single-trait) epistatic mapping strategies. We also apply mvMAPIT framework to protein sequence data from two broadly neutralizing anti-influenza antibodies and approximately 2,000 heterogenous stock of mice from the Wellcome Trust Centre for Human Genetics. The mvMAPIT R package can be downloaded at https://github.com/lcrawlab/mvMAPIT.

## Introduction

Genome-wide association (GWA) studies have contributed substantially in the discovery of genetic markers associated with the architecture of disease phenotypes^1–6^. Epistasis, commonly defined as the interaction between genetic loci, has long been thought to play a key role in defining the genetic architecture underlying many complex traits and common diseases^7–11^. Indeed, previous studies have detected pervasive epistasis in many model organisms^12–35^. Substantial contributions of epistasis to phenotypic variance have been revealed for many complex traits^36,37^ and have been suggested to constitute an important component of evolution^38^. Furthermore, modeling epistasis in addition to additive and dominant effects has been shown to increase phenotypic prediction accuracy in model organisms^39–41^ and facilitate genomic selection in breeding programs^42–44^. Despite a longstanding and currently ongoing debate about the contribution of non-additive effects on the architecture of human complex traits^22,45–52^, recent genetic mapping studies have also identified evidence of epistatic interactions that significantly contribute to quantitative traits and diseases^53–56^, and some have recently shown that gene-by-gene interactions can drive heterogeneity of causal variant effect sizes across diverse human populations^57^. Importantly, epistasis is often proposed as a key contributor to missing heritability — the proportion of heritability not explained by the top associated variants in GWA studies^7,58–61^.

Many statistical methods have been developed to facilitate the identification of epistasis in complex traits and diseases. Generally, these existing tools can be classified into two frameworks. In the first framework, explicit searches are performed to detect significant pairwise or higher-order interactions. More specifically, they use various strategies including exhaustive search^62–64^, probabilistic search^65^, or prioritization based on a predefined set of biological annotations of signaling pathways or genomic regulatory units^66,67^. Different statistical paradigms have been implemented for these explicit search-based approaches including various frequentist tests^62,68,69^, Bayesian inference^70–73^, and, most recently, detecting epistasis using machine learning^74,75^. Indeed, the explosion of large-scale genomic datasets has provided the unique opportunity to integrate many of these techniques as standard statistical tools within GWA analyses. Many modern GWA applications have datasets that can include hundreds of thousands of individuals genotyped at millions of markers and phenotyped for thousands of traits^76,77^. Due to the potentially large space of genetic interactions (e.g., *J* (*J* – 1)/2 possible pairwise combinations for *J* variants in a study), explicit search-based methods often suffer from heavy computational burden. Even with various efficient computational improvements^65,68,78–80^, exploring over a large combinatorial domain remains a daunting task for many epistatic mapping studies. More importantly, because of a lack of *a priori* knowledge about epistatic loci, exploring all possible combinations of genetic variants can result in low statistical power after correcting for multiple hypothesis tests.

As a departure from the explicit search strategy, the second category of epistatic mapping methods attempts to address the previously mentioned challenges by detecting *marginal* epistasis. Specifically, instead of directly identifying individual pairwise or higher-order interactions, these approaches focus on identifying variants that have a non-zero interaction effect with any other variant in the dataset. For example, the “MArginal ePIstasis Test” (MAPIT)^81^ assesses each variant (in turn) and identifies candidate markers that are involved in epistasis without the need to identify the exact partners with which the variants interact — thus, alleviating much of the statistical power concerns and heavy computational burdens associated with explicit search-based methods. As a framework, the marginal epistatic strategy has been implemented in both linear mixed models and machine learning and has been used for case-control studies^82^, pathway enrichment applications^83^, heritability estimation^12^, and even extended to explore different sources of non-additive genetic variation (e.g., gene-by-environment interactions)^84,85^. However, despite its wide adoption, this approach can still be underpowered for traits with low heritability or “polygenic” traits which are generated by many mutations of small effect^81^.

To date, both the explicit search and marginal epistasis detection methodologies have only focused on analyzing one phenotype at a time. However, many previous genetic association studies have extensively shown that jointly modeling multiple phenotypes can often dramatically increase power for variant detection^86^. In this work, we present the “multivariate MArginal ePIstasis Test” (mvMAPIT) — a multi-outcome generalization of the MAPIT model which aims to take advantage of the relationship between traits to improve the identification of variants involved in epistasis. We formulate mvMAPIT as a multivariate linear mixed model (mvLMM) and extend a previously developed variance component estimation algorithm for efficient parameter inference and *P*-value computation in the multi-trait setting^87^. Together with reasonable model approximations, our proposed approach is scalable to moderately sized GWA studies. With detailed simulations, we illustrate the benefits of mvMAPIT in terms of providing effective type I error control and compare its power to the univariate (or single-trait) mapping strategy used in the original MAPIT model. Here, part of our main contribution is the demonstration that the power in our proposed multivariate approach is driven by the correlations between the effects of pairwise interactions on multiple traits. To close, we also apply the mvMAPIT framework to protein sequence data from a nearly combinatorially complete library of two broadly neutralizing anti-influenza antibodies^88^ and to 15 quantitative hematology traits assayed in a heterogenous stock of mice from Wellcome Trust Centre for Human Genetics^89–91^.

## Results

### Overview of the multivariate marginal epistasis test

The “multivariate MArginal ePIstasis Test” (mvMAPIT) is a multi-outcome extension of the statistical framework MAPIT which aims to identify variants that are involved in epistatic interactions by leveraging the covariance structure of non-additive genetic variation that is shared between multiple traits. The key idea behind the concept of marginal epistasis is to identify variants that are involved in epistasis while avoiding the need to explicitly conduct an exhaustive search over all possible interactions between pairs of variants. As an overview of mvMAPIT and its corresponding software implementation, we will assume that we have access to a GWA study on *N* individuals denoted as 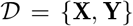 where **X** is an *N* × *J* matrix of genotypes with *J* denoting the number of SNPs (each of which is encoded as {0, 1, 2} copies of a reference allele at each locus *j*) and **Y** denoting a *D* × *N* matrix holding *D* different traits that are measured for each of the *N* individuals. We will let **y**_*d*_ represent the *N*-dimensional phenotypic vector for the *d*-th trait. For convenience, we will assume that the genotype matrix and the traits of interest have been mean-centered and standardized. Unlike standard exhaustive search tests for epistasis, mvMAPIT works by examining one variant at a time. For the *j*-th variant, we consider the following mvLMM formulation

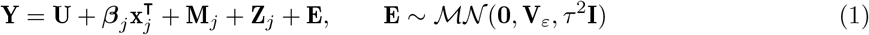

where **U** is a *D* × *N* dimensional matrix which contains population-level intercepts that are the same for all individuals within each trait; **x**_*j*_ is an *N*-dimensional vector for the *j*-th genotype that is the focus of the model; ***β***_*j*_ is a *D*-dimensional vector of additive effects for the *j*-th genotype; 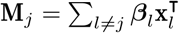 is the combined additive effects from all other *l* ≠ *j* variants across the *D* traits with effect sizes ***β***_*l*_ and effectively represents the polygenic background of all variants except for the *j*-th; and 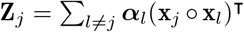 is the summation of all pairwise interaction effects **x**_*j*_ ∘ **x**_*l*_ (i.e., element-wise multiplication) between the *j*-th variant and all other *l* ≠ *j* variants with regression coefficients ***α***_*l*_ across the *D* traits; and **E** denotes an *D* × *N* matrix of residual errors that is assumed to follow a matrix-variate normal distribution with mean **0**, within column covariance **V**_*ε*_ among the *D* traits, and independent within row covariance (scaled by *τ*^2^) among the *N* individuals in the study. The term **Z**_*j*_ is the main focus of the model and represents the collection of marginal epistatic effects of the *j*-th variant — formally defined as the summation of its epistatic interaction effects with all other variants. In this study, we will demonstrate mvMAPIT while analyzing *D* = 2 traits at a time, but note that the framework can easily be applied to more phenotypes (see Materials and Methods). Similarly, while we focus on pairwise statistical epistasis in the above formulation, extension of the mvMAPIT framework to detect higher order interactions is straightforward^81^.

The model specified in Eq. (1) becomes an underdetermined linear system for many modern GWA applications (i.e., in biobanks where genotyped markers *J* > *N* individuals). As a result, we need to make additional modeling assumptions on the regression coefficients to make the generative model identifiable. Here, we follow standard linear modeling approaches^81,92–95^ by first letting **B** = [***β***_*l*_]_*l*≠*j*_ and **A** = [***α***_*l*_]_*l*≠*j*_ denote matrices of coefficients. Then we assume that these matrices follow matrixvariate normal distributions where 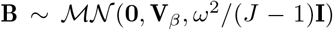 and 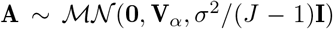, respectively. With the probabilistic assumption of normally distributed effect sizes, the model defined in Eq. (1) is equivalent to a multivariate variance component model where 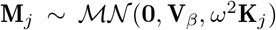 with 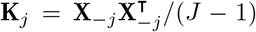 being an additive genetic relatedness matrix that is computed using all genotypes other than the *j*-th SNP; and 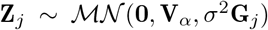 with **G**_*j*_ = **D**_*j*_**K**_*j*_**D**_*j*_ being a non-additive relatedness matrix computed based on all pairwise interaction terms involving the *j*-th SNP. Here, we let **D**_*j*_ = diag(**x**_*j*_) denote an *N* × *N* diagonal matrix with the *j*-th genotype as its only nonzero elements. It is also important to note that both **K**_*j*_ and **G**_*j*_ change with every new *j*-th marker that is tested. The key takeaway from this variance component model formulation is that *σ*^2^ represents a measure on the marginal epistatic effect for each variant in the data.

The goal of mvMAPIT is to identify variants that have non-zero interaction effects with any other variant in the data across multiple traits. To accomplish this, we examine each SNP in turn and assess the null hypothesis *H*_0_: *σ*^2^ = 0. In practice, we use a computationally efficient method of moments algorithm called MQS^87^ to estimate model parameters and to carry out calibrated statistical tests within mvMAPIT. More specifically, to estimate variance components, we first (right) multiply Eq. (1) by a variant-specific projection (or hat) matrix 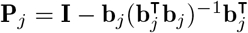 with **b** = [**1**; **x**_*j*_] and **1** being an *N*-dimensional vector of ones. This procedure projects the model onto a column space that is orthogonal to the intercept and the genotypic vector of interest **x**_*j*_ which allows us to rewrite Eq. (1) as the following

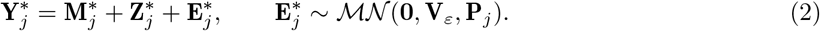

Here, in addition to previous notation, 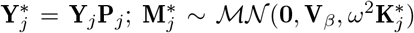 with 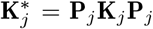; and 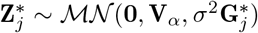 with 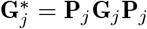. The joint analysis of multiple traits requires a generalization of the MQS algorithm to also include method of moments estimators for covariance components between outcomes. Without loss of generality, let 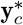 and 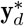 be the *c*-th and *d*-th rows of the measured phenotypic matrix 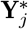. Our multivariate extension of MQS implements an approach which first fits univariate models (i.e., the setting where *c* = *d*) and then combines the resulting *P*-values with those stemming from a “covariance statistic” which looks for shared marginal epistatic effects between all pairwise combinations of the *D* traits. The MQS estimate for the marginal epistatic component takes on the quadratic form

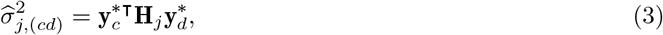

where 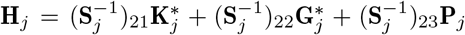 with elements (**S**_*j*_)_*rs*_ = tr(**Σ**_*jr*_**Σ**_*js*_) for matrices subscripted as 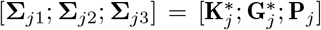, and tr(•) is used to denote the matrix trace function. The corresponding standard error for the test statistic in Eq. (3) can be approximated as the following^87,96^

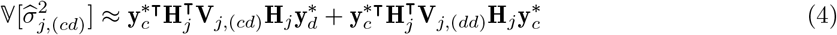

with 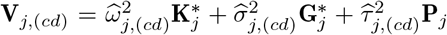, being the covariance between any two traits of interest. Note the indices *c* and *d* range over all *D* traits and that a different 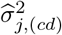 is computed for all pairwise combinations of the *c*-th and *d*-th traits in the data.

We implement a combinatorial strategy to carry out hypothesis testing and derive *P*-values using the test statistics computed in Eq. (3). This is done in three key steps. In the first step, we fit the univariate models for all *D* traits of interests. This case mirrors the original MAPIT model. In mvMAPIT, this means that the variance component point estimate is computed using only one trait row in **Y** (i.e, *c* = *d*). Here, we use a hybrid approach where we first implement a normal test for each variant by default, and then we apply an exact method for the variants that have *P*-values from the normal test that fall below the nominal significance threshold of 0.05 to correct for possible inflation^81^. To implement the normal test, we simply compute a z-score by dividing the test statistic in Eq. (3) by its standard deviation in Eq. (4) with **V**_*j*,(*cd*)_ = **V**_*j*,(*dd*)_. For the SNPs needing the exact test, we utilize the fact that the MQS variance component estimate follows a mixture of chi-square distributions under the null hypothesis. This is derived from both the standard normality assumption on each trait **y**\* and the quadratic form of the statistic in Eq. (3). More specifically, we say that 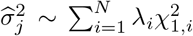 where 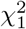 are chi-square random variables with one degree of freedom and (λ_1_,…, λ_*N*_) are the eigenvalues of

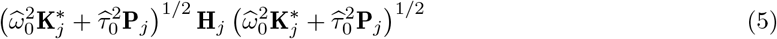

with 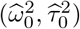 being the MQS estimates of 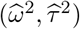 under the null hypothesis. Several approaches have been suggested to obtain *P*-values under a mixture of chi-square distributions. In this work, we use the Davies^97^ method (see Data and Software Availability).

In the second step of the hypothesis testing procedure, we derive *P*-values for the hypothesis that a given variant is interacting with others in determining traits *c* and *d* (where *c* ≠ *d*). This amounts to deriving covariance components for all pairwise combinations of traits where Eq. (3) takes on a bilinear form. In this setting, we again use a normal test this time by dividing each covariance test statistic with its standard deviation in Eq. (4). As we will show below, the *P*-values derived for the covariance components with the asymptotic normal approximation tend to have generally conservative behavior with respect to type I error control under the null hypothesis. Indeed, deriving an exact test to guard against deflation and potentially exhibit better power under the alternative could be done; however, we do not explore this line of work here.

In the third and final step of the hypothesis testing, we combine all *P*-values from the first two steps into an overall marginal epistatic *P*-value. Each individual *P*-value corresponds to the effect one variant has on the variance of one trait or covariance between a pair of traits. The combined *P*-value corresponds to the marginal epistatic effect that one variant has on a set of traits. Without loss of generality, assume that we are studying *D* = 2 traits. In this case, we would have *T* = 3 sets of *P*-values (two marginal sets from **y**_1_ and **y**_2_ individually and one covariance set from analyzing {**y**_1_, **y**_2_} together). We combine *P*-values using two different strategies. The first assumes that each of the *t* = 1,…, *T* tests are (effectively) independent and implements Fisher’s method^98^ which combines *P*-values into a single chi-square test statistic using the formula 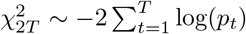 where *p_t_* denotes the *P*-value from the *t*-th test. The second approach assumes an unknown dependency structure between each of the *T* tests and computes a harmonic mean^99^ *P*-value where 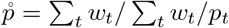. The term 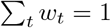 represents a sum weights which we uniformly set to be *w_t_* = 1/*T* for all *P*-values. There are many complex traits for which epistatic effects are assumed to make small contributions to their overall broad-sense heritability^50–52^. Intuitively, this combinatorial approach is meant to aggregate over the signal identified in one trait and leverage the genetic correlation between traits to improve power. A full theoretical derivation of mvMAPIT and details about its corresponding software implementation can be found in Materials and Methods.

#### Note on settings where mvMAPIT is designed to be most powered

The formulation of the general estimates in Eq. (3) and (4) highlight an important takeaway in that the mvMAPIT covariance statistic models epistatic pairs that together affect the architecture of multiple traits. It is not meant to identify individual SNPs that are involved in epistasis for multiple traits while being a member of different interacting pairs. To clarify this, consider two simple scenarios in Figure 1 where we have two phenotypes (**y**_1_ and **y**_2_) that are generated by a combination of four SNPs (**x**_1_, **x**_2_, **x**_3_, **x**_4_). In the first scenario, we say that (in expectation) 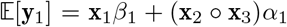 and 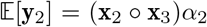 (Figure 1A); while, in the second scenario, 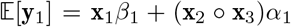 and 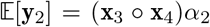 (Figure 1B). The key to power in the mvMAPIT framework is that, in the first scenario, the interaction between **x**_2_ and **x**_3_ appears in both traits with nonzero correlation between the effect sizes *α*_1_ and *α*_2_. This is in contrast to the second scenario where there is a common variant involved in epistasis but it is a member of two different sets of interactions that affect each trait. The mvMAPIT covariance statistic captures the situation illustrated in the first scenario (Figure 1A) but not in the second (Figure 1B).

**Figure 1.**
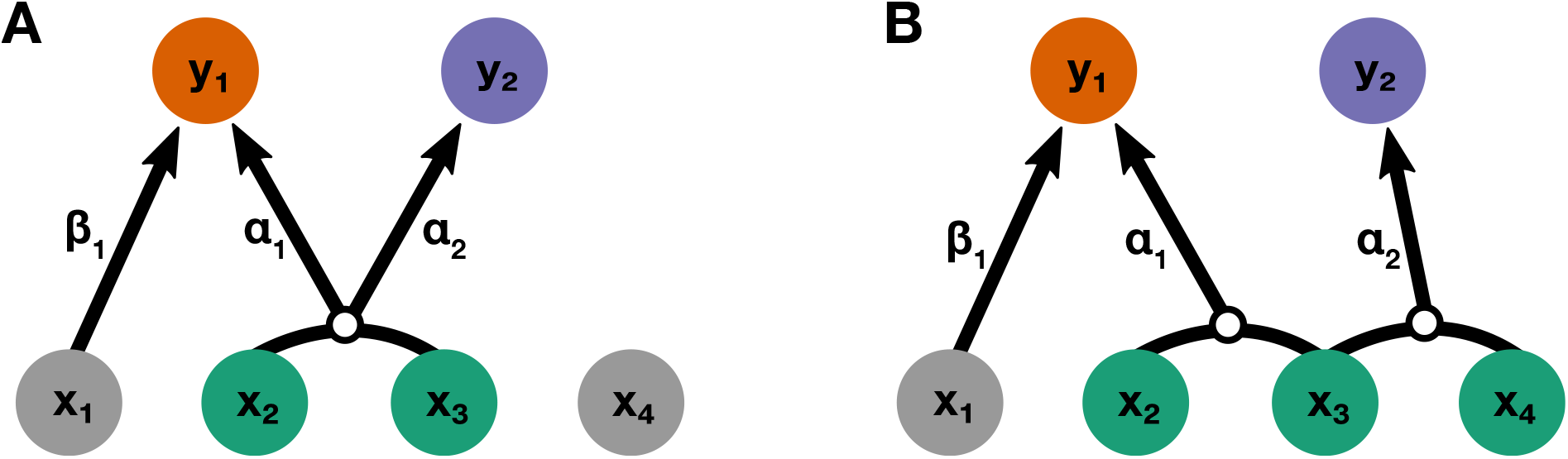
Schematic of the types of shared interactions modeled by the multivariate marginal epistasis test. Consider two simple, proof-of-concept simulation scenarios where two traits (**y**_1_, **y**_2_) are generated by a combination of four SNPs (**x**_1_, **x**_2_, **x**_3_, **x**_4_). Panel **(A)** shows the first scenario where (in expectation) 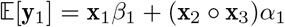 and 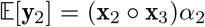. Panel **(B)** shows the second scenario where 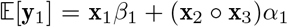 and 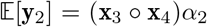. In both panels, variant **x**_1_ only has an additive effect *β*_1_ on trait **y**_1_. The mvMAPIT approach models correlations between the effects of a given interaction on multiple traits. Therefore, mvMAPIT is designed to identify SNPs involved in the first scenario where the interaction between variants **x**_2_ and **x**_3_ is shared between traits with nonzero correlated effect sizes *α*_1_ and *α*_2_. This is in contrast to the second case, where variant **x**_3_ is important to both traits but through distinct interactions with variants **x**_2_ and **x**_4_, respectively.

### mvMAPIT produces calibrated *P*-values and conservative type I error rates

In this section, we make use of a previously described simulation scheme^12,81^ in order to investigate whether mvMAPIT and its combinatorial inference approach preserves the desired type I error rate and produces well-calibrated *P*-values under the null hypothesis. Here, we generate synthetic phenotypes using real genotypes from the 22nd chromosome of the control samples in the Wellcome Trust Case Control Consortium (WTCCC) 1 study^100^. Altogether, these data consist of *N* = 2,938 individuals and *J* = 5,747 SNPs. Since the goal of mvMAPIT is to search for variants involved in epistatic interactions, we consider the null model to be satisfied when the phenotypic variation of the synthetic traits are solely driven by additive effects. Here, we first subsample the genotypes for *N* = 1,000, 1,750, and 2,500 observations. Next, we randomly select 1,000 causal SNPs and simulate continuous phenotypes by using the linear model **Y** = **BX**^┬^ + **E**. The additive effect sizes for each causal SNP are drawn as 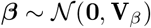 across traits, and then we scale all terms to ensure a narrow-sense heritability of 60%. In these simulations, we vary the correlation of the additive genetic effects such that we have traits with independent additive effects (*v*_*β*,12_ = 0), traits with highly correlated additive effects (*v*_*β*,12_ = 0.8), and traits with perfectly correlated additive effects (*v*_*β*,12_ = 1). We assess the calibration of the *P*-values that are produced by mvMAPIT during each of the three key steps in its combinatorial hypothesis testing procedure. That is, we evaluate (1) the *P*-values resulting from the univariate test on each trait, (2) the *P*-values derived from the covariance test, and (3) the final overall *P*-value that is computed by combining the first two sets of *P*-values via Fisher’s method or the harmonic mean. Note that we expect the *P*-values from the first univariate test to be well-calibrated since it is equivalent to the MAPIT model. Figures 2 and S1–S2 show the quantile-quantile (QQ) plots based on *P*-values combined using Fisher’s method while Figures S3–S5 depicts results while using the harmonic mean. Similarly, Tables 1 and S1–S5 show the empirical type I error rates estimated for mvMAPIT at significance levels *P* = 0.05, 0.01, and 0.001, respectively.

**Figure 2.**
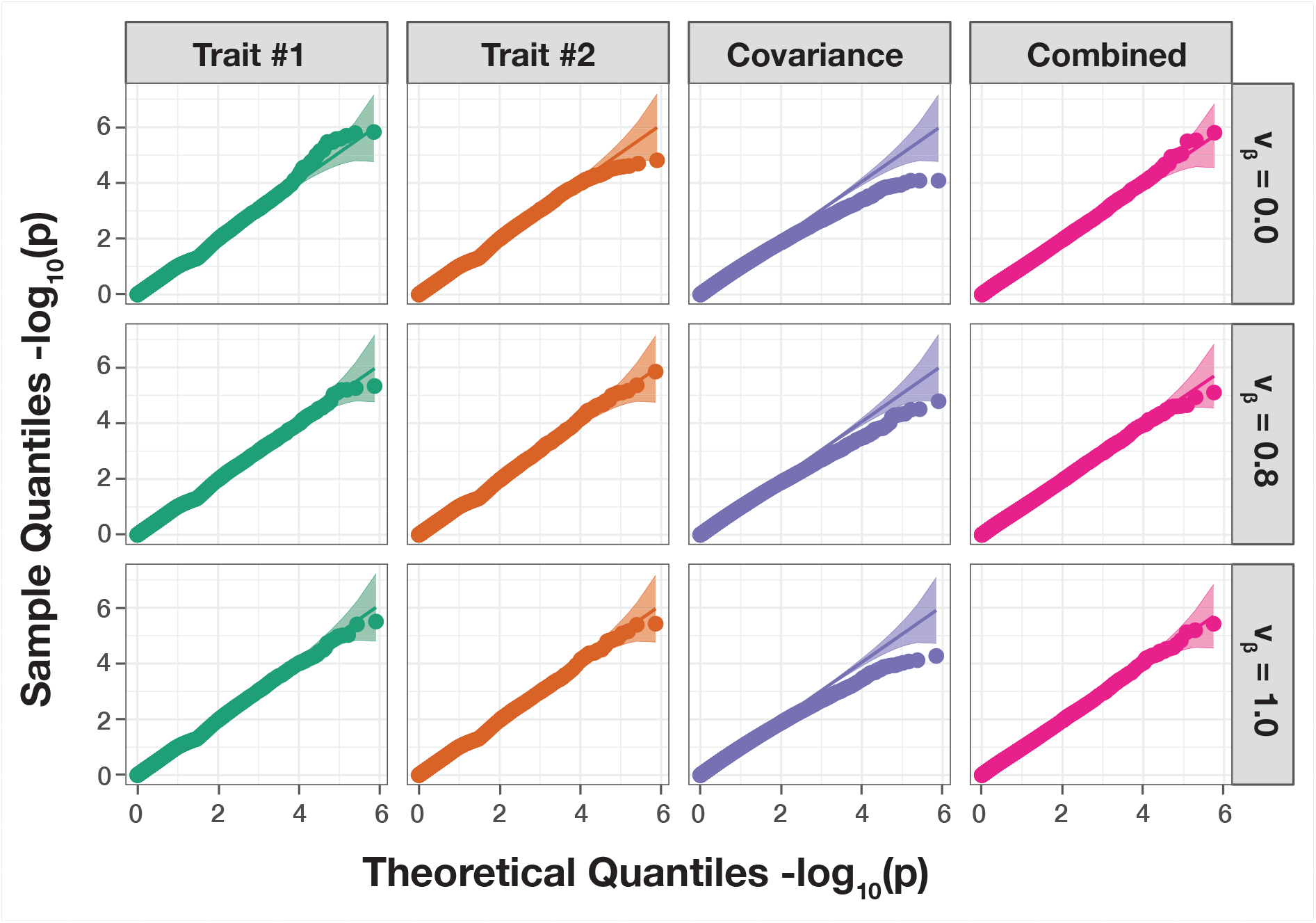
The mvMAPIT framework using Fisher’s method produces well-calibrated *P*-values when traits are generated by only additive effects (sample size *N* = 2,500 individuals). In these simulations, quantitative traits are simulated to have narrow-sense heritability *h*^2^ = 0.6 with an architecture made up of only additive genetic variation. Each row of quantile-quantile (QQ) plots corresponds to a setting where the additive genetic effects for a causal SNP have different correlation structures across traits. In these simulations, we consider scenarios where we have independent traits (*v_β_* = 0), highly correlated traits (*v_β_* = 0.8), and perfectly correlated traits (*v_β_* = 1). The first two columns show *P*-values resulting from the univariate MAPIT test on “trait #1” and “trait #2”, respectively. The third column depicts the “covariance” *P*-values which corresponds to assessing the pairwise interactions affecting both traits is. Lastly, the fourth column shows the final “combined” *P*-value which combines the *P*-values from the first three columns using Fisher’s method. The 95% confidence interval for the null hypothesis of no marginal epistatic effects is shown in grey. Each plot combines results from 100 simulated replicates.

**Table 1.** The mvMAPIT framework using Fisher’s method preserves type I error rates under the null model when traits are generated by only additive effects (sample size *N* = 2,500 individuals). In these simulations, quantitative traits are simulated to have narrow-sense heritability *h*^2^ = 0.6 with an architecture made up of only additive genetic variation. Each row corresponds to a setting where the additive genetic effects for a causal SNP have different correlation structures across traits. In these simulations, we consider scenarios where we have traits with independent additive effects (*v_β_* = 0), traits with highly correlated additive effects (*v_β_* = 0.8), and traits with perfectly correlated additive effects (*v_β_* = 1). We assess the calibration of the *P*-values that are produced by mvMAPIT during each of the three key steps in its combinatorial hypothesis testing procedure (see Materials and Methods). We show type I error rates resulting from *P*-values taken from the “univariate” test on each trait independently, the “covariance” *P*-values which corresponds to assessing the pairwise interactions affecting both traits, and the final “combined” *P*-value. Results are summarized over 100 simulated replicates. Values in the parentheses are the standard deviations across replicates.

| | Add. Effect Corr. | $P = 0.05$ | $P = 0.01$ | $P = 0.001$ |
| --- | --- | --- | --- | --- |
| Univariate | $v_\beta = 0.0$ | 0.030 ($1 \times 10^{-2}$) | 0.009 ($2 \times 10^{-3}$) | 0.0010 ($9 \times 10^{-4}$) |
| | $v_\beta = 0.8$ | 0.030 ($1 \times 10^{-2}$) | 0.009 ($2 \times 10^{-3}$) | 0.0010 ($7 \times 10^{-4}$) |
| | $v_\beta = 1.0$ | 0.030 ($1 \times 10^{-2}$) | 0.009 ($3 \times 10^{-3}$) | 0.0009 ($7 \times 10^{-4}$) |
| Covariance | $v_\beta = 0.0$ | 0.040 ($1 \times 10^{-2}$) | 0.006 ($2 \times 10^{-3}$) | 0.0003 ($3 \times 10^{-4}$) |
| | $v_\beta = 0.8$ | 0.040 ($1 \times 10^{-2}$) | 0.007 ($2 \times 10^{-3}$) | 0.0004 ($5 \times 10^{-4}$) |
| | $v_\beta = 1.0$ | 0.040 ($1 \times 10^{-2}$) | 0.006 ($2 \times 10^{-3}$) | 0.0003 ($4 \times 10^{-4}$) |
| Combined | $v_\beta = 0.0$ | 0.040 ($1 \times 10^{-2}$) | 0.006 ($2 \times 10^{-3}$) | 0.0003 ($3 \times 10^{-4}$) |
| | $v_\beta = 0.8$ | 0.040 ($1 \times 10^{-2}$) | 0.007 ($2 \times 10^{-3}$) | 0.0004 ($5 \times 10^{-4}$) |
| | $v_\beta = 1.0$ | 0.040 ($1 \times 10^{-2}$) | 0.006 ($2 \times 10^{-3}$) | 0.0003 ($4 \times 10^{-4}$) |

Overall, mvMAPIT conservatively controls type 1 error rate, both in the presence of nonzero correlation between additive effects on the two traits and even with small sample sizes in the data. This result holds regardless of how *P*-values are combined in the model. The QQ-plots of the *P*-values for all three components in mvMAPIT follow the expected uniform distribution for the univariate and combined analysis. Notably, because of the approximations used to compute the standard error of the test statistic in Eq. (18), the multivariate extension of the MQS-based testing procedure in mvMAPIT can result in conservative *P*-values for the covariance components under the null.

### Improved detection of epistatic variants using mvMAPIT in simulations

We test the power of mvMAPIT across different genetic trait architectures via an extensive simulation study (Materials and Methods). Once again, we generate synthetic phenotypes using real genotypes from the 22nd chromosome of the control samples in the WTCCC 1 study^100^. As a reminder, these data consist of *N* = 2,938 individuals and *J* = 5,747 SNPs. In these simulations, we randomly choose 1,000 causal variants to directly affect the genetic architecture of *D* = 2 phenotypes. All causal SNPs are assumed to have a non-zero additive effect on both traits. Next, we randomly select a set of epistatic variants from the 1,000 causal SNPs and divide them into two interacting groups (again see Materials and Methods). We will denote these groups #1 and #2 as 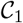 and 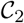, respectively, with 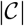 denoting the cardinality of the group. One may interpret the epistatic SNPs in 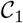 as being the “hub nodes” in an interaction network where each of these variants interact with all of the SNPs assigned to 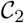. We generate synthetic traits by using the multivariate linear model **Y** = **BX^┬^** + **AW^┬^** + **E** where, in addition to previous notation, **W** is matrix of interactions between the SNPs assigned to the groups 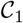 and 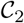. The additive and interaction coefficients for causal SNP effects across traits are drawn as 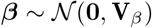 and 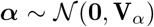, respectively. As a final step, we scale all terms to ensure that all genetic effects explain a fixed proportion of the total phenotypic variation. We assume a wide-range of simulation scenarios by varying the following parameters:

1. broad-sense heritability: *H*^2^ = 0.3 and 0.6;
2. proportion of phenotypic variation that is explained by additive effects: *ρ* = 0.5 and 0.8;
3. number of causal SNPs assigned to the interaction groups: 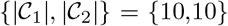 and {10, 20};
4. correlation between epistatic effects: *v*_*α*,12_ = 0 and 0.8.

All results presented in this section are based on 100 different simulated phenotypes for each parameter combination.

The main point of these simulations is to highlight the potential power gained from taking a multivariate approach to epistatic detection. To that end, in each of the simulation scenarios, we assess (*i*) the power of running the univariate MAPIT model on each trait individually, (*ii*) the marginal epistatic effects detected by the covariance test, and (*iii*) the power from the overall association identified by mvMAPIT. Figures 3 and S6–S8 show the empirical power of the univariate MAPIT model, the covariance test, and mvMAPIT while using Fisher’s method at various multiple hypothesis testing correction thresholds. Figures S9–S12 depict the same information but with mvMAPIT using the harmonic mean to combine *P*-values. We also compare each method’s ability to rank true positives over false positives via receiver operating characteristic (ROC) and precision-recall curves (Figures 4 and S13–S15). There are several key takeaways from these simulation results. Overall, the ability of the univariate MAPIT framework to detect group #1 and #2 causal variants depends on the proportion of non-additive phenotypic variation that they explain. This has been shown in previous demonstrations of the method^81^. For example, when there are 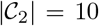 causal SNPs in group #2, each variant in the set is expected to explain (1 – *ρ*)*H*^2^/10% of the genetic variance. As we increase that number of causal SNPs in group #2 to 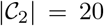, this proportion of variance explained by SNPs in group #2 will decrease which will make it more difficult to prioritize markers involved in interactions. Importantly, it is worth noting that the single-phenotypic test in MAPIT depends on the total interaction effects, rather than individual pairwise effects or the number of interacting pairs. An example of this can be seen by comparing Figure 3A to Figure S2A where the ability to group #1 variants is independent of the number of variants in group #2.

**Figure 3.**
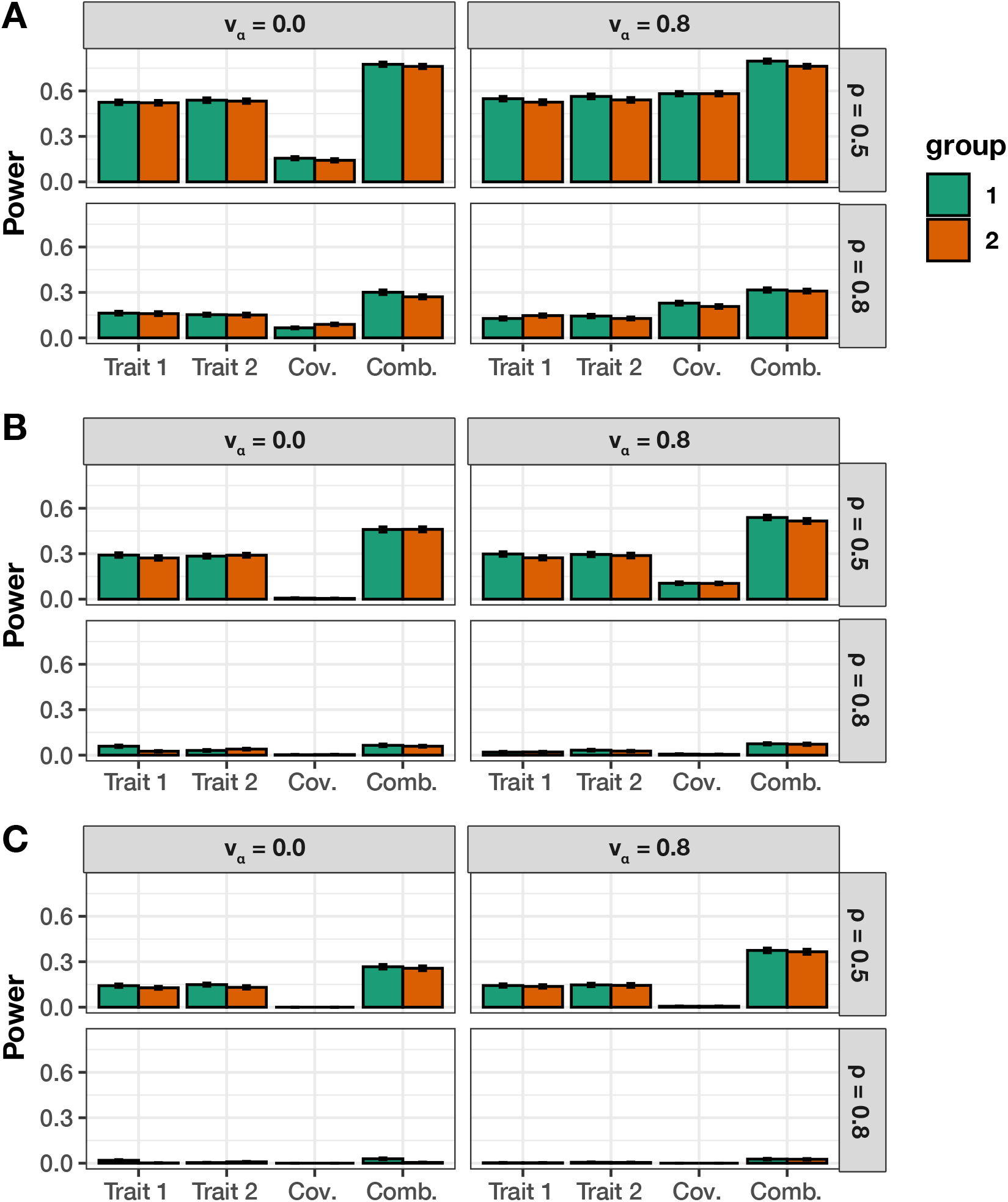
Empirical power of mvMAPIT with Fisher’s method to detect group #1 (10) and group #2 (10) epistatic variants across complex traits with moderate broad-sense heritability. In these simulations, both quantitative traits are simulated to have broad-sense heritability *H*^2^ = 0.6 with architectures made up of both additive and epistatic effects. The parameter *ρ* = {0.5, 0.8} is used to determine the portion of broad-sense heritability contributed by additive effects. Each column corresponds to a setting where the epistatic effects for interactive pairs have different correlation structures across traits. In these simulations, we consider scenarios where we have traits with independent epistatic effects (*v_α_* = 0) and traits with highly correlated epistatic effects (*v_α_* = 0.8). This plot shows the empirical power of mvMAPIT at significance levels **(A)** *P* = 5 × 10^−2^, **(B)** *P* = 5 × 10^−4^, and **(C)** *P* = 1 × 10^−5^, respectively. Group #1 and #2 causal markers are colored in green and orange, respectively. For comparison, the “trait #1” and “trait #2” bars correspond to the univariate MAPIT model, the “cov” bars corresponds to power contributed by the covariance test, and “comb” details power from the overall association identified by mvMAPIT in the combination approach. Results are based on 100 simulations per parameter combination and the horizontal bars represent standard errors.

**Figure 4.**
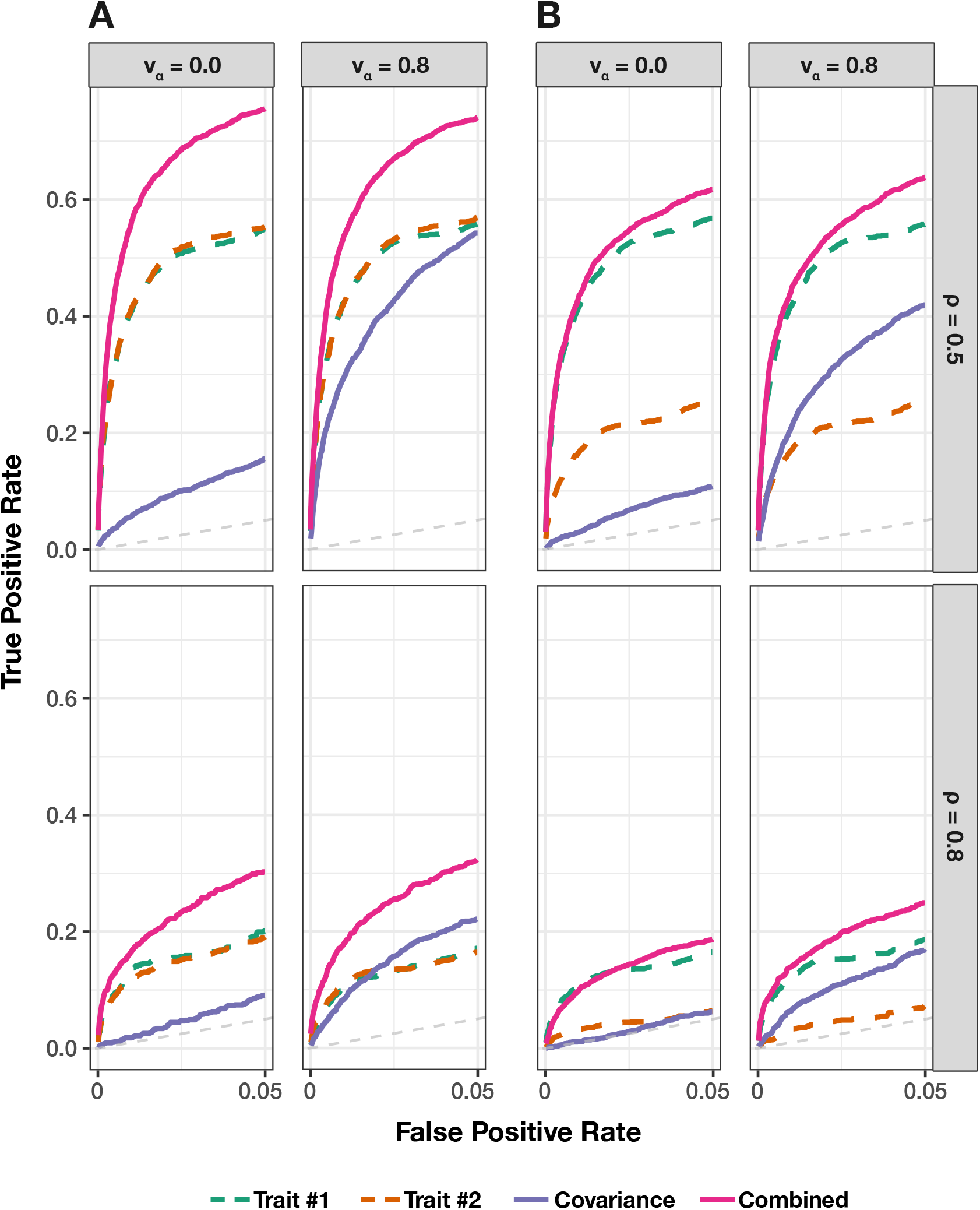
Receiver operating characteristic (ROC) curves comparing the ability of mvMAPIT with Fisher’s method to the univariate MAPIT model in detecting group #1 (10) and group #2 (10) epistatic variants across complex traits. In panel **(A)** both traits have broad-sense heritability *H*^2^ = 0.6; while in panel **(B)** one of traits has broad-sense heritability *H*^2^ = 0.6 and the other has heritability *H*^2^ = 0.3. Across the rows, the parameter *ρ* = {0.5,0.8} is used to determine the portion of broad-sense heritability contributed by additive effects. Each column corresponds to settings where the epistatic effects across traits are independent (*v_α_* = 0) or highly correlated (*v_α_* = 0.8). For comparison, the “trait #1” and “trait #2” dotted lines correspond to the univariate MAPIT model, the “covariance” solid purple line corresponds to power contributed by the covariance test, and the “combined” pink line shows power from the overall association identified by mvMAPIT in the multivariate approach. Note that the upper limit of the x-axis (i.e., false positive rate) has been truncated at 0.05. All results are based on 100 simulated replicates.

There are two situations where mvMAPIT shows significant gains over the univariate MAPIT modeling approach. Intuitively, the first case is when there is nonzero correlation between the effects of the epistatic interactions shared between traits (e.g., when *v*_*α*,12_ = 0.8). The sensitivity of the covariance hypothesis test depends on the strength of this correlation which can help increase power when combining over *P*-values in the final step of mvMAPIT. This becomes increasingly relevant in the low heritability cases. Figures 4 and S13–S15 demonstrate that the sensitivity of the covariance statistic is comparable to the univariate statistic for highly correlated epistatic effects (*v_α_* = 0.8) despite genetic variance being predominantly explained by additivity (*ρ* = 0.8). Secondly, using mvMAPIT to jointly analyze traits with shared genetic architecture but different levels of heritability provides a viable approach for studying non-additive variation in traits with low heritability. In Figures 4, S7, S8, and S11–S15, we simulated synthetic traits such that one has a moderate broad-sense heritability *H*^2^ = 0.6 and the other has heritability *H*^2^ = 0.3. In these scenarios, detecting variants involved in interactions increased for the trait with low heritability. In particular, the covariance component analysis is shown to play an important role in this improved detection (e.g., see Figure 4B).

### Synergistic epistasis in binding affinity landscapes for neutralizing antibodies

We apply the mvMAPIT framework to protein sequence data from Phillips et al. ^88^ who generated a nearly combinatorially complete library for two broadly neutralizing anti-influenza antibodies (bnAbs), CR6261 and CR9114. This dataset includes almost all combinations of one-off mutations that distinguish between germline and somatic sequences which total to *J* = 11 heavy-chain mutations for CR6261 and *J* =16 heavy-chain mutations for CR9114. Theoretically, a combinatorially complete dataset for 11 and 16 mutations will have 2,048 and 65,536 samples, respectively. In this particular study, we have have access to *N* = 1,812 complete observations for CR6261 and *N* = 65,091 complete measurements for CR9114. For our analysis with mvMAPIT, residue sequence information was encoded as a binary matrix with the germline sequence residues marked by zeros and the somatic mutations represented as ones. As quantitative traits, Phillips et al. ^88^ measure the binding affinity of the two antibodies to different influenza strains. Here, we assess the contribution of epistatic effects when binding to *H*_1_ and *H*_9_ for CR6261, and *H*_1_ and *H*_3_ for CR9114.

Once again, we report results after running mvMAPIT with Fisher’s method and the harmonic mean (Table S6). Figures 5A and S16A show Manhattan plots for *P*-values corresponding to the trait-specific marginal epistatic tests (i.e., the univariate MAPIT model), the covariance test, and the mvMAPIT approach. Here, green colored dots are positions that have significant marginal epistatic effects beyond a Bonferroni corrected threshold for multiple testing (*P* = 0.05/11 = 4.55 × 10^−3^ for CR6261 and *P* = 0.05/16 = 3.13 × 10^−3^ for CR9114, respectively). Interestingly, while the univariate MAPIT approach was able to identify significant marginal epistatic effects for CR6261, it lacked the power to identify significant positions driving non-additive variation in binding affinity for CR9114. Overall, the combined trait approach in mvMAPIT revealed marginal epistatic effects for positions 29, 35, 82, 83, and 84 in CR6261, and positions 30, 36, 57, 64, 65, 66, 82, and 83 for CR9114. Most notably, these same positions were also identified as contributing to pairwise epistasis by Phillips et al. ^88^. In the original study, the authors first ran an exhaustive-search to statistically detect significant interactions and then conducted downstream analyses to find that these positions are likely responsible for the antibodies binding to the influenza surface protein hemagglutinin. The regression coefficients from the exhaustive search, as reported by Phillips et al.^88^, are illustrated in panels B and C of Figures 5 and S16. Panel B illustrates interaction coefficients when assessing binding of CR6261with *H*_1_ (upper right triangle) and *H*_9_ (lower left triangle). Panel C shows the same information when assessing binding of CR9114 with *H*_1_ (upper right triangle) and *H*_9_ (lower left triangle). Our results show that mvMAPIT identifies all required mutations in these systems as well as most positions involved in at least one epistatic pair.

**Figure 5.**
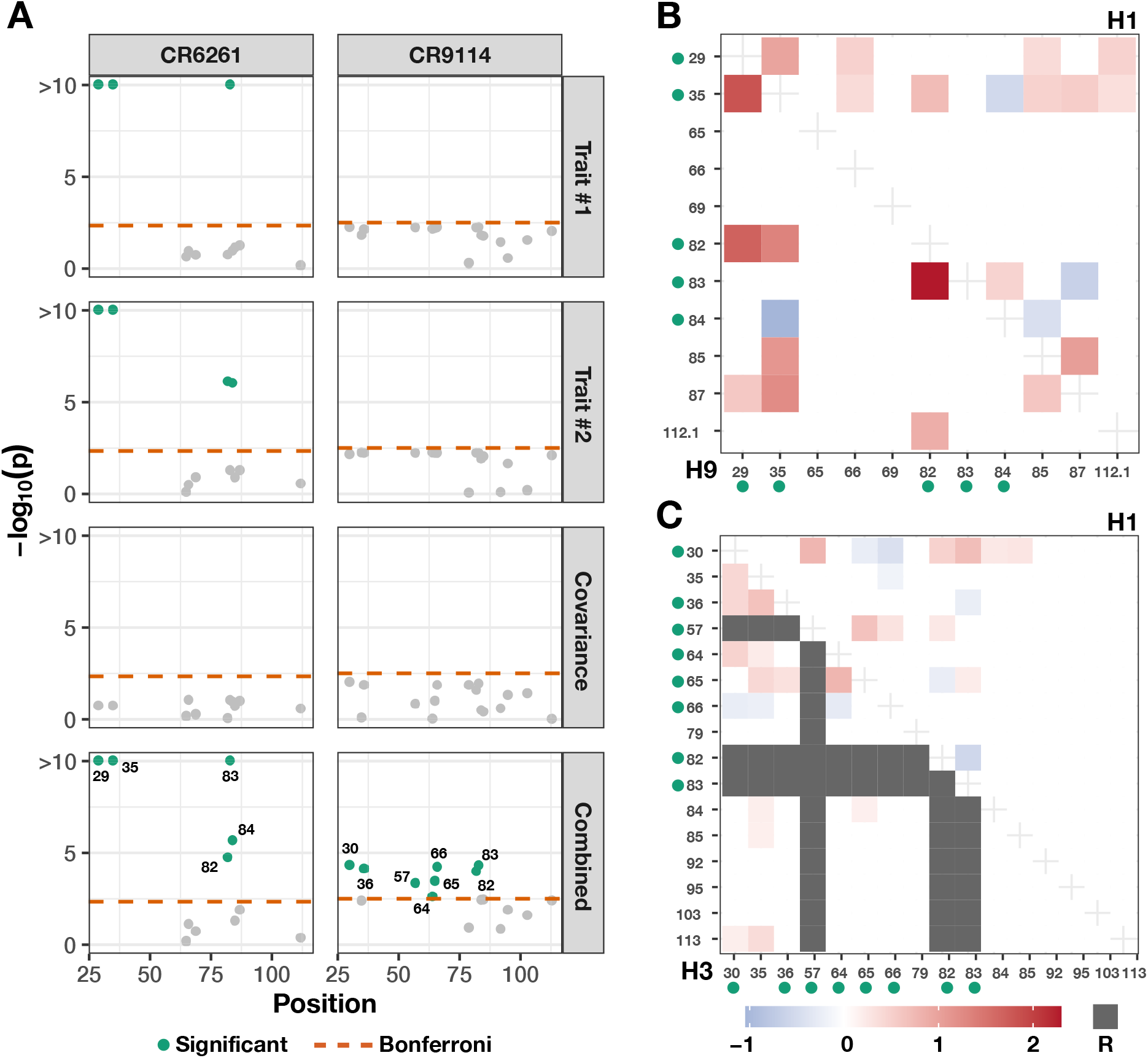
Applying mvMAPIT with Fisher’s method to broadly neutralizing antibodies recovers heavy-chain mutations known to be involved in epistatic interactions that affect binding against two influenza strains. These results are based on protein sequence data from Phillips et al. ^88^ who generated a nearly combinatorially complete library for two broadly neutralizing anti-influenza antibodies (bnAbs), CR6261 and CR9114. For each antibody, we assess binding affinity to different influenza strains. For CR6261, traits #1 and #2 are binding measurements to the antigens *H*_1_ and *H*_9_; while, for CR9114, we assess the same measurement for *H*_1_ and *H*_3_. Panel **(A)** shows Manhattan plots for the different sets of *P*-values computed during the mvMAPIT analysis. The red horizontal lines indicate a chain-wide Bonferroni corrected significance threshold (*P* = 4.55 × 10^−3^ for CR6261 and *P* = 3.13 × 10^−3^ for CR9114, respectively). The green colored dots are positions that have significant marginal epistatic effects after multiple correction. Panels **(B)** and **(C)** reproduce exhaustive search results originally reported by Phillips et al.^88^. The green dots next to the mutation labels on the axes are the residues that are significant in the multivariate MAPIT analysis and correspond to panel **(A)**. The shaded regions in panel **(B)** are the regression coefficients for pairwise interactions between positions when assessing binding of CR6261with *H*_1_ (upper right triangle) and *H*_9_ (lower left triangle). Similarly, panel **(C)** shows the same information when assessing binding of CR9114 with *H*_1_ (upper right triangle) and *H*_3_ (lower left triangle). Required mutations (indicated by R) are plotted in gray and left out of the analysis^88^.

### Joint modeling of hematology traits yields epistatic signal in stock of mice

In this section, we apply mvMAPIT to individual-level genotypes and 15 hematology traits in a heterogeneous stock of mice dataset from the Wellcome Trust Centre for Human Genetics^89–91^. This collection of data contains approximately *N* = 2,000 individuals depending on the phenotype (see Materials and Methods), and each mouse has been genotyped at *J* = 10,346 SNPs. As noted by previous studies, these data represent a realistic mixture of the simulation scenarios we detailed in the previous sections (i.e., varying different values of the parameter *ρ*). Specifically, this stock of mice is known to be genetically related with population structure and the genetic architectures of these particular traits have been shown to have different levels of broad-sense heritability with varying contributions from non-additive genetic effects.

For each pairwise trait analysis, we provide a summary table which lists the combined *P*-values after running mvMAPIT with Fisher’s method and the harmonic mean (Table S7). We also include results corresponding to the univariate MAPIT model and the covariance test for comparison. Overall, the single-trait marginal epistatic test only identifies significant variants for the large immature cells (LIC) after Bonferroni correction (*P* = 4.83 × 10^−6^). A complete picture of this can be seen in Figures S17 and S18 which depict Manhattan plots of our genome-wide interaction study for all combinations of trait pairs. Here, we can see that most of the signal in the combined *P*-values from mvMAPIT likely stems from the covariance component portion of the model. This hypothesis holds true for the joint pairwise analysis of (*i*) hematocrit (HCT) and hemoglobin (HGB) and (*ii*) mean corpuscular hemoglobin (MCH) and mean corpuscular volume (MCV) (e.g., see the third and fourth rows of Figures 6 and S19). One explanation for observing more signal in the covariance components over the univariate test could be derived from the traits having low heritability but high correlation between epistatic interaction effects. Recall that our simulation studies showed that the sensitivity of the covariance statistic increased for these cases. Notably, the non-additive signal identified by the covariance test is not totally dependent on the empirical correlation between traits (see Figure S20). Instead, as previously shown in our simulation study, the power of mvMAPIT over the univariate approach occurs when there is correlation between the effects of epistatic interactions shared between two traits. Importantly, many of the candidate SNPs selected by the mvMAPIT framework have been previously discovered by past publications as having some functional nonlinear relationship with the traits of interest. For example, the multivariate analysis with traits MCH and MCV show a significant SNP rs4173870 (*P* = 4.89 × 10^−10^) in the gene hematopoietic cell-specific Lyn substrate 1 (*Hcls1*) on chromosome 16 which has been shown to play a role in differentiation of erythrocytes^101^. Similarly, the joint analysis of HGB and HCT shows hits in multiple coding regions. One example here are the SNPs rs3692165 (P = 1.82 × 10^−6^) and rs13482117 (*P* = 8.94 × 10^−7^) in the gene calcium voltage-gated channel auxiliary subunit alpha2delta 3 (*Cacna2d3*) on chromosome 14, which has been associated with decreased circulating glucose levels^102^, and SNP rs3724260 (*P* = 4.58 × 10^−6^) in the gene *Dicer1* on chromosome 12 which has been annotated for anemia both in humans and mice^103^.

**Figure 6.**
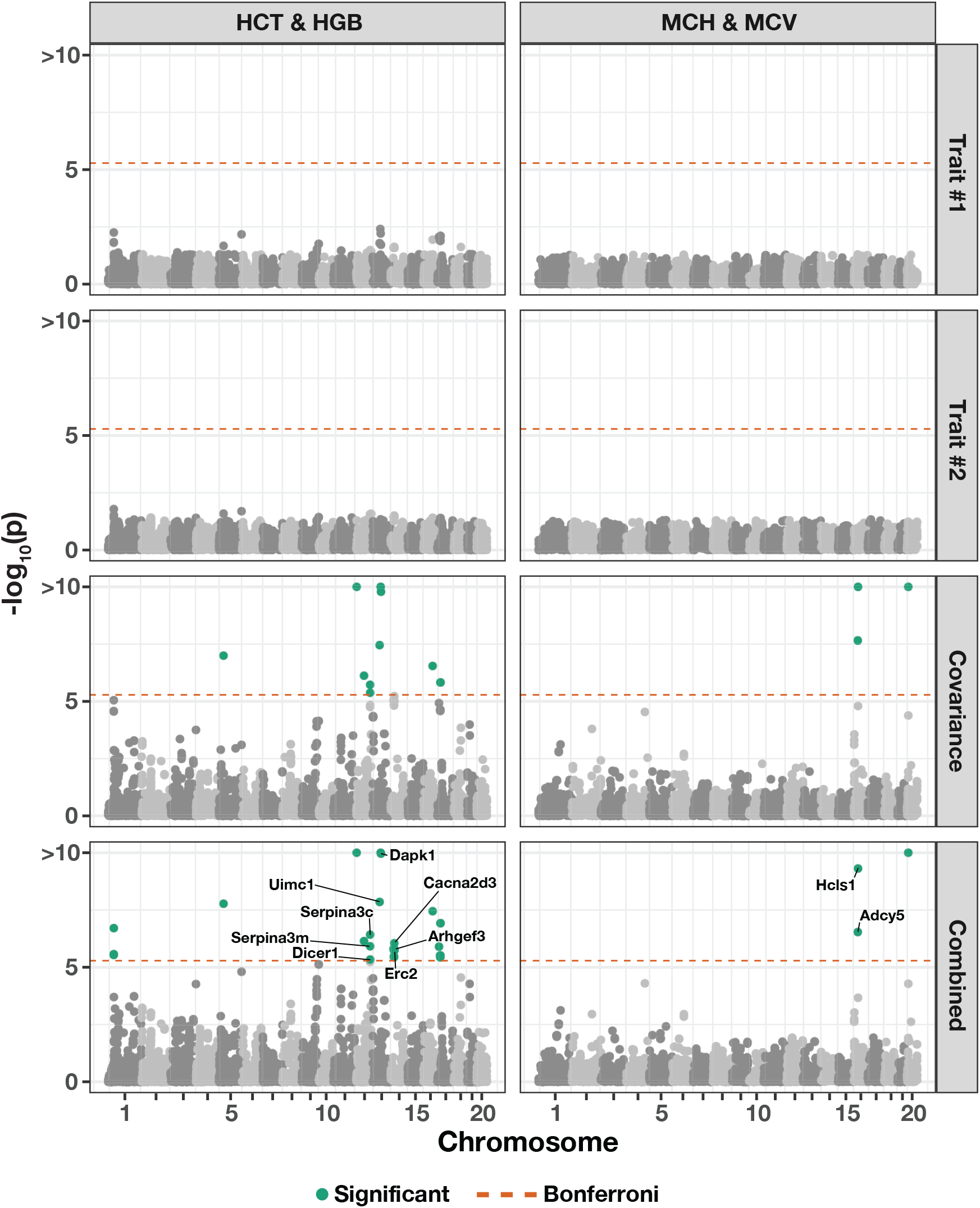
Manhattan plot of genome-wide interaction study for two pairs of hematology traits in the heterogenous stock of mice dataset from the Wellcome Trust Centre for Human^89–91^ using mvMAPIT with Fisher’s method. The trait pairs in this figure include hematocrit (HCT) and hemoglobin (HGB) in the left column and mean corpuscular hemoglobin (MCH) and mean corpuscular volume (MCV) in the right column. Here, we depict the *P*-values computed during each step of the mvMAPIT modeling pipeline. The red horizontal lines indicate a genome-wide Bonferroni corrected significance threshold (*P* = 4.83 × 10^−6^). The green colored dots are SNPs that have significant marginal epistatic effects after multiple test correction. Significant SNPs were mapped to the closest neighboring genes using the Mouse Genome Informatics database (http://www.informatics.jax.org)^106,107^.

Table 2 lists a select subset of SNPs in coding regions of genes that have been associated with phenotypes related to the hematopoietic system, immune system, or homeostasis and metabolism. Each of these are significant (after correction for multiple hypothesis testing) in the mvMAPIT analysis of related hematology traits. Some of these phenotypes have been reported as having large broad-sense heritability, which improves the ability of mvMAPIT to detect the signal. For example, the genes *Arf2* and *Cacna2d3* are associated with phenotypes related to glucose homeostasis, which has been reported to have a large heritable component (estimated *H*^2^ = 0.3 for insulin sensitivity^104^). Similarly, the genes *App* and *Pex1* are associated with thrombosis where (an estimated) more than half of phenotypic variation has been attributed to genetic effects (estimated *H*^2^ ≥ 0.6 for susceptibility to common thrombosis^105^).

**Table 2.** Notable SNPs with marginal epistatic effects after applying the mvMAPIT framework to 15 hematology traits in the heterogenous stock of mice dataset from the Wellcome Trust Centre for Human Genetics^89–91^. In the first two columns, we list SNPs and their genetic location according to the mouse assembly NCBI build 34 (accessed from Shifman et al. ^135^) in the format `Chromosome:Basepair`. Next, we give the results stemming from univariate analyses on traits #1 and #2, respectively, the covariance (cov) test, and the overall *P*-value derived by mvMAPIT using Fisher’s method. The last columns detail the closest neighboring genes found using the Mouse Genome Informatics database http://www.informatics.jax.org) ^106,107^, a short summary of the suggested annotated function for those genes, and the reference to the source of the annotation. See Table S7 for the complete list of SNP and SNP-set level results.

| SNP | Location | Trait #1 | Trait #2 | Trait #1 *P*-value | Trait #2 *P*-value | Cov. *P*-value | Comb. *P*-value | Gene | Genomic Annotation | Reference |
| --- | --- | --- | --- | --- | --- | --- | --- | --- | --- | --- |
| rs3699393 | 2:5887012 | MCV | PLT | 0.21 | 0.23 | $5.75 \times 10^{-7}$ | $4.9 \times 10^{-6}$ | *Upf2* | anemia and abnormal bone marrow cell development | 125 |
| rs13478092 | 5:3601413 | LIC | PLT | 0.034 | 0.58 | $1.67 \times 10^{-10}$ | $1.26 \times 10^{-9}$ | *Pex1* | abnormal venous thrombosis | 126 |
| rs3694887 | 5:102770070 | ALY | LIC | $1.26 \times 10^{-4}$ | 0.013 | $2.54 \times 10^{-6}$ | $1.55 \times 10^{-9}$ | *Aff1* | abnormal B and T cell number and morphology | 127 |
| rs3694887 | 5:102770070 | LIC | PLT | 0.013 | 0.28 | $5.47 \times 10^{-27}$ | $4.49 \times 10^{-26}$ | *Aff1* | abnormal B and T cell number and morphology | 127 |
| rs13478923 | 6:99475169 | ALY | LIC | $2.8 \times 10^{-4}$ | 0.12 | $1.79 \times 10^{-6}$ | $1.81 \times 10^{-8}$ | *Foxp1* | abnormal B cell differentiation, physiology, count | 128,129 |
| rs13478924 | 6:99571626 | ALY | LIC | $3.11 \times 10^{-4}$ | 0.12 | $2.70 \times 10^{-6}$ | $2.86 \times 10^{-8}$ | *Foxp1* | abnormal B cell differentiation, physiology, count | 128,129 |
| rs13478985 | 6:115245823 | MCV | WBC | 0.16 | 0.40 | $1.14 \times 10^{-81}$ | $1.34 \times 10^{-78}$ | *Atg7* | decreased bone marrow cell count | 130,131 |
| rs3723163 | 11:103800737 | HCT | LYM | 0.072 | 0.30 | $3.99 \times 10^{-107}$ | $2.66 \times 10^{-104}$ | *Arf2* | decreased fasting circulating glucose level | 102 |
| rs3723163 | 11:103800737 | HGB | WBC | 0.069 | 0.25 | $1.85 \times 10^{-7}$ | $6.76 \times 10^{-7}$ | *Arf2* | decreased fasting circulating glucose level | 102 |
| rs3724260 | 12:100163212 | HGB | HCT | 0.030 | 0.062 | $1.44 \times 10^{-5}$ | $4.58 \times 10^{-6}$ | *Dicer1* | anemia | 103 |
| rs3692165 | 14:27756640 | HCT | HGB | 0.026 | 0.037 | $9.9 \times 10^{-6}$ | $1.8 \times 10^{-6}$ | *Cacna2d3* | decreased circulating glucose level | 102 |
| rs13482117 | 14:27614362 | HCT | HGB | 0.023 | 0.03 | $5.9 \times 10^{-6}$ | $9.0 \times 10^{-7}$ | *Cacna2d3* | decreased circulating glucose level | 102 |
| rs13482288 | 14:81840412 | ALY | BAS | 0.036 | 0.65 | $1.78 \times 10^{-8}$ | $1.1 \times 10^{-7}$ | *Tdrd3* | abnormal B cell differentiation and physiology | 132 |
| rs4173870 | 16:35764290 | MCH | MCV | 0.14 | 0.71 | $1.20 \times 10^{-11}$ | $4.89 \times 10^{-10}$ | *Hcls1* | differentiation of erythrocytes | 101 |
| rs4212102 | 16:84204704 | PLT | WBC | 0.17 | 0.35 | $1.16 \times 10^{-10}$ | $2.44 \times 10^{-9}$ | *App* | increased susceptibility to induced thrombosis | 105,133 |
| rs4212186 | 16:84273330 | PLT | WBC | 0.17 | 0.36 | $5.88 \times 10^{-11}$ | $1.31 \times 10^{-9}$ | *App* | increased susceptibility to induced thrombosis | 105,133 |
| rs3711994 | 19:45078018 | ALY | LYM | $3.71 \times 10^{-4}$ | 0.10 | $1.04 \times 10^{-12}$ | $2.80 \times 10^{-14}$ | *Btrc* | abnormal lymphocyte morphology | 134 |

## Discussion

The marginal epistatic testing strategy offers an alternative to traditional epistatic mapping methods by seeking to identify variants that exhibit non-zero interaction effects with any other variant in the data^81–83^. This framework has been shown to drastically reduce the number of statistical tests needed to uncover evidence of significant non-additive variation in complex traits and, as a result, alleviates much of the empirical power concerns and heavy computational burden associated with explicit search-based methods. Still, models testing for marginal epistasis can be underpowered when applied to traits with low heritability or to “polygenic” traits where the interactions between mutations have small effect sizes^81^. In this work, we present the “multivariate MArginal ePIstasis Test” (mvMAPIT), a multi-outcome extension of the univariate marginal epistatic framework. Theoretically, we formulate mvMAPIT as a multivariate linear mixed model (mvLMM) where its ability to jointly analyze any number of traits relies on a generalized “variance-covariance” component estimation algorithm^87^. Through extensive simulations, we show that mvMAPIT preserves type I error rates and produces well-calibrated *P*-values under the null model when traits are generated only by additive effects (Figures 2 and S1–S5, and Tables 1 and S1–S5). In these simulation studies, we also show that mvMAPIT improves upon the identification of epistatic variants over the univariate test when there is correlation between the effects of genetic interactions shared between multiple traits (Figures 1, 3, and 4, and S6–S15). By analyzing two real datasets, we demonstrated the ability of mvMAPIT to recover heavy-chain mutations known to be involved in epistatic interactions that affect binding against two influenza strains^88^ (Figures 5 and S16, and Table S6) as well as to identify hematology trait relevant epistatic SNPs in heterogenous stock of mice^89–91^ that have also been detected in previous publications and functional validation studies (Figures 6 and S17–S20, and Tables 2 and S7). Lastly, we have made mvMAPIT an open-source R software package with documentation to facilitate its use by the greater scientific community.

The current implementation of the mvMAPIT framework offers many directions for future development and applications. First, like other marginal epistatic mapping methods, mvMAPIT is unable to directly identify detailed interaction pairs despite being able to identify SNPs that are involved in epistasis. As shown through our simulations and real data analyses, being able to identify SNPs involved in epistasis allows us to come up with an initial (likely) set of variants that are worth further exploration, and thus represents an important first step towards identifying and understanding detailed epistatic associations. In previous studies^66,81,108,109^, two-step *ad hoc* procedures have been suggested where, in our case, we would first run mvMAPIT and then focus on significant SNPs from the first step to further test all of the pairwise interactions among them in order to identify specific epistatic interaction pairs. While this approach has been shown to be effective in univariate (single-trait) analyses, this two-step procedure is still *ad hoc* in nature and could miss important epistatic associations. Exploring robust ways unify these two steps in a joint fashion would be an interesting area for future research. Second, in its current implementation, mvMAPIT can be computationally expensive for datasets with large sample sizes (e.g., hundreds of thousands of individuals in a biobank scale study). In this study, we develop a “variance-component component” extension to the MQS algorithm to estimate parameters in the mvMAPIT model. Theoretically, MQS is based on the method of moments and produces estimates that are mathematically identical to the Haseman-Elston (HE) cross-product regression^87,110,111^. In practice, MQS is not only computationally more efficient than HE regression, but also provides a simple, analytic estimation form that allows for exact *P*-value computation — thus alleviating the need for jackknife re-sampling procedures^112^ that both are computationally expensive and rely on assumptions of independence across individuals in the data^113^. Exploring different ways to reliably fit large-scale mvLMMs with multiple random effects is a consideration for future work. For example, as an alternative, recent studies have proposed randomized multi-component versions of HE regression for heritability estimation which scale up to datasets with millions of individuals and variants, respectively^114–116^. It would be interesting to develop a well-calibrated hypothesis test within the randomized HE regression setting so that it may be implemented within the mvMAPIT software for association mapping.

In the future, we plan to expand the mvMAPIT framework to also identify individual variants contributing other sources of non-additive genetic variation such as gene-by-environment (G×E) or gene-by-sex (G×Sex) interactions. We can do this by manipulating the marginal epistatic covariance matrix in Eq. (1) to encode how loci interact with one or more environmental instruments^84,85,116,117^. Lastly, we have focused here on applying mvMAPIT to simple quantitative traits. However, there are many important traits with significant non-additive genetic components in plants, animals, and humans that cannot be easily reduced to simple scalar values. Examples include longitudinal traits such as growth curves^118^, metabolic traits such as the relative concentrations of different families of metabolites^119^, and morphological traits such as shape or color^120^. Indeed, each of these traits can be decomposed into vectors of interrelated components, but treating these components as independent phenotypes within existing univariate epistatic mapping tools would be inefficient because of their statistical dependence. As an alternative, the mvMAPIT framework can be used to make joint inferences about epistasis across any number of correlated phenotypic components—which, in the case of longitudinal studies for example^121–124^, could be used to interrogate how non-additive variation of trait architecture changes or evolves over time.

## URLs

Multivariate marginal epistasis test (mvMAPIT) software, https://github.com/lcrawlab/mvMAPIT; univariate marginal epistasis test (MAPIT) software, https://github.com/lcrawlab/mvMAPIT; Wellcome Trust Centre for Human Genetics, http://mtweb.cs.ucl.ac.uk/mus/www/mouse/index.shtml; Mouse Genome Informatics database, http://www.informatics.jax.org.

## Acknowledgments

We thank members of the Weinreich and Crawford Labs for insightful comments on earlier versions of this manuscript as well as Sohini Ramachandran for helpful discussions. This research was conducted in part using computational resources and services at the Center for Computation and Visualization at Brown University. This research was supported in part by a David & Lucile Packard Fellowship for Science and Engineering awarded to L. Crawford. Any opinions, findings, and conclusions or recommendations expressed in this material are those of the author(s) and do not necessarily reflect the views of any of the funders.

## Author Contributions

LC conceived the study. DW and LC supervised the project and provided resources. JS, AD, and LC developed the methods. JS developed the software and performed the analyses. All authors wrote and revised the manuscript.

## Materials and Methods

### The marginal epistasis test for single traits

The original motivation behind the original “MArginal ePIstasis Test” (MAPIT) was to identify variants that are involved in epistasis while avoiding the need to explicitly conduct an exhaustive search over all possible pairwise interactions^81^. In this section, we give a statistical overview of the univariate version of MAPIT where the objective is to search for marginal epistatic effects (i.e., the combined pairwise interaction effects between a given variant and all other variants) that drive the genetic architecture of single traits. To begin, consider a genome-wide association (GWA) study with *N* individuals who have been genotyped for *J* single nucleotide polymorphisms (SNPs) encoded as {0, 1, 2} copies of a reference allele at each locus. In the MAPIT framework, we examine one SNP at a time (indexed by *j*) and consider the following linear model

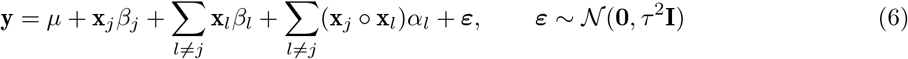

where **y** is an *N*-dimensional vector of phenotypic states for a quantitative trait of interest measured in the *N* individuals; *μ* is an intercept term; **X** denotes an *N* × *J* matrix of genotypes with **x**_*j*_ and **x**_*l*_ representing *N*-dimensional vectors for the *j*-th and *l*-th SNPs; *β_j_* and *β_l_* are the respective additive effects; **x**_*j*_ ∘ **x**_*l*_ denotes the Hadamard (element-wise) product between two genotypic vectors with corresponding interaction effect size *α_l_*; ***ε*** is a normally distributed error term with mean zero and scale variance term *τ*^2^; and **I** denotes an *N* × *N* identity matrix. For convenience, we will assume that both the genotype matrix (column-wise) and trait of interest have been mean-centered and standardized. It is also worth noting that, while we limit the above to the task of identifying second order (i.e., pairwise) interactions between genetic variants, extensions of MAPIT to higher-order epistatic and gene-by-environmental effects have been shown to be straightforward to implement^84,85,116,117^.

#### Variance component model formulation

Since many modern GWA applications present scenarios that would make Eq. (6) an underdetermined linear system (i.e., in biobanks where genotyped markers *J* > *N* individuals), the MAPIT framework follows other standard approaches^81,92–95^ to ensure model identifiability by assuming that the additive and interaction effect sizes follow univariate normal distributions where 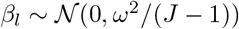 and 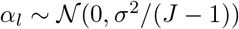 for *l* ≠ *j*, respectively. This key normal assumption on the regression coefficients allows for Eq. (6) to be equivalently represented as the following variance component model

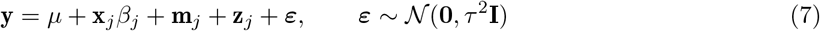

where, in addition to previous notation, 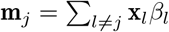 is the combined additive effects from all variants other than the *j*-th; and 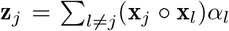 denote the summation of all pairwise interaction effects between the *j*-th variant and all other variants. Under the variance component formulation in Eq. (7), the two random effects can also be expressed probabilistically as 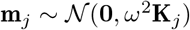 where 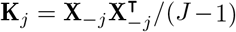 is an additive genetic relatedness matrix that is computed using all genotypes other than the *j*-th SNP, and 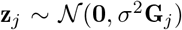 where **G**_*j*_ = **D**_*j*_**K**_*j*_**D**_*j*_ is a non-additive relatedness matrix computed based on all pairwise interaction terms involving the *j*-th SNP. Here, we let **D**_*j*_ = diag(**x**_*j*_) denote an *N* × *N* diagonal matrix with the *j*-th genotype as its only nonzero elements. It is also important to note that both **K**_*j*_ and **G**_*j*_ change with every new *j*-th marker that is tested.

#### Univariate point estimates

Intuitively, the key takeaway from the variance component model formulation is that *σ*^2^ represents a measure on the marginal epistatic effect for each variant in the data. Therefore, in order to identify variants that have significant non-zero interaction effects, we must assess the null hypothesis *H*_0_: *σ*^2^ =0 for each variant in the dataset. The original MAPIT framework uses a computationally efficient method of moments algorithm called MQS^87^ to estimate model parameters and to carry out calibrated statistical tests. Briefly, MQS produces point estimates that are mathematically identical to the Haseman-Elston (HE) cross-product regression^87,110,111^. To implement this algorithm, we first specify a two-dimensional matrix **b**_*j*_ = [**1**, **x**_*j*_] with **1** being an *N*-dimensional vector of ones, and then we multiply both sides of Eq. (7) by a variant-specific projection 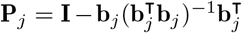 which maps the model onto a column space that is orthogonal to the intercept and the genotypic vector **x**_*j*_. This process simplifies the model specification of MAPIT to the following

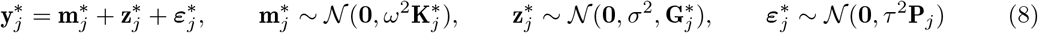

where we denote 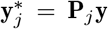; 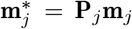; 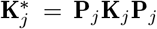; 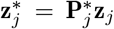; 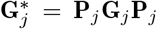; and 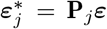, respectively. The method of moments estimator for the variance components in Eq. (8) is naturally based on the second moment matching equations where, in expectation, we have

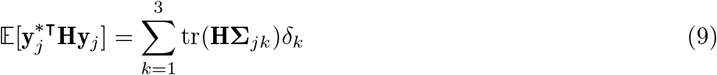

where **H** is a symmetric and non-negative definite matrix used to create weighted second moments, tr(•) denotes the trace of a matrix, and we use shorthand to represent 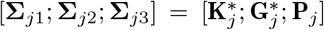 and ***δ*** = (*ω*^2^, *σ*^2^, *τ*^2^), respectively. In practice, we replace the left hand side of Eq. (9) with the realized value 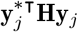. Note that many choices of **H** will yield unbiased estimates for (*ω*^2^, *σ*^2^, *τ*^2^), but different choices of **H** can affect statistical efficiency of the estimates. The set of moment matching equations in MQS is generated by using the covariance matrices corresponding to the variance components in place of the arbitrary **H**. This system of equations then can be rewritten as the following matrix multiplication

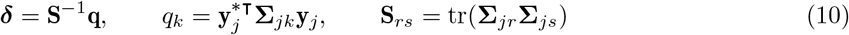

where **q** is a 3-dimensional vector and **S** is a 3 × 3 dimensional matrix with *k*, *r*, *s* ∈ {1, 2, 3} being indices to represent the different variance components. If we subset just to compute an estimate for the marginal epistatic variance component (i.e., for the second index), then Eq. (10) reduces to the following formula

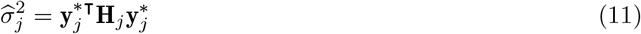

where the variant-specific matrix 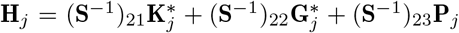 is now used in place of the arbitrary **H**.

#### Univariate hypothesis testing

In general, there are two ways to compute *P*-values in the MAPIT framework^81^. The first option uses a two-sided z-score or normal test. This particular test only requires the variance component estimate 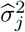 from Eq. (11) and its corresponding standard error, which is approximated in MQS approach by

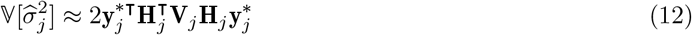

where 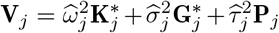. The second option for deriving *P*-values in the MAPIT framework uses an exact test which is based on the fact that the MQS variance component estimate follows a mixture of chi-square distributions under the null hypothesis. This is derived from both the normality assumption on **y**\* and the quadratic form of the statistic in Eq. (11). Namely, 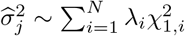 where 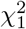 are chi-square random variables with one degree of freedom and (λ_1_,…, λ_*N*_) are the eigenvalues of the matrix

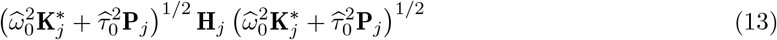

with 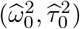 being the MQS estimates of 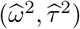 under the null hypothesis. Several approaches have been suggested to obtain *P*-values under a mixture of chi-square distributions, including the Davies method^97^ (see Data and Software Availability). In practice, while the Davies method is an exact test and is expected to produce calibrated *P*-values, it can become computationally intensive since it scales cubically in the number of individuals *N*. On the other hand, while the normal test only scales quadratically in *N* because of the variance approximation in Eq. (12), it has been shown to lead to mis-calibrated *P*-values for datasets with small sample sizes. As result, MAPIT uses a hybrid procedure which uses the normal test by default, and then applies the Davies method when the *P*-value from the normal test is below the threshold of 0.05^81^.

### Derivation of the multivariate marginal epistasis test

The “multivariate MArginal ePIstasis Test” (mvMAPIT) is a multi-outcome generalization of the MAPIT framework which aims to improve upon the identification of variants that are involved in genetic interactions by leveraging the correlation structure between multiple traits. Once again, consider a GWA study with *N* individuals this time who have been measured for *D* different phenotypes. We will denote these sets of outcomes via a *D* × *N* dimensional matrix 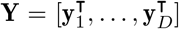 with **y**_*d*_ denoting an *N*-dimensional phenotypic vector for the *d*-th trait. Given the *j*-th variant of interest, we specify the mvMAPIT approach as the following multivariate linear mixed model (mvLMM)^86^

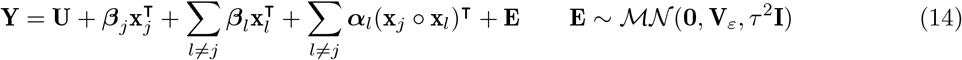

where, in addition to previous notation, **U** is a *D* × *N* dimensional matrix which contains population-level intercepts that are the same for all individuals within each trait; ***β***_*j*_ and ***β***_*l*_ are *D*-dimensional vectors of additive effects for the *j*-th and *l*-th genotypic vectors; ***α***_*l*_ is a *D*-dimensional vector of coefficients for the interaction effects between the *j*-th and *l*-th SNPs spanning all traits; and **E** denotes an *D* × *N* matrix of residual errors that is assumed to follow a matrix-variate normal distribution with mean **0**, within column covariance **V**_*ε*_ among the *D* traits, and independent within row covariance (scaled by *τ*^2^) among the *N* individuals in the study.

Similar to the univariate setting, we need to make additional probabilistic assumptions to ensure model identifiability when Eq. (14) is an underdetermined linear system. To that end, let **B** = [***β***_*l*_]_*l*≠*j*_ and **A** = [***α***_*l*_]_*l*≠*j*_ denote the collection of coefficients not involving the *j*-th variant of interest. Here, we will assume that these *D* × (*J* – 1) effect size matrices also follow matrix-variate normal distributions where 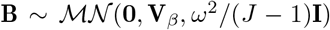 and 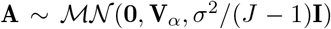, respectively. Note that this formulation is largely similar to the univariate case except with the additional property that the phenotypes being studied share some genetic covariance through **V**_*β*_ and **V**_*α*_. This assumption, coupled with the affine transformation property of matrix normal distributions, allows for us to equivalently represent the mvMAPIT model in Eq. (14) as the following multivariate variance component model

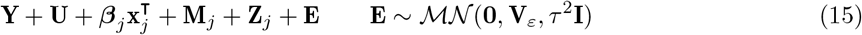

where 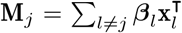 with 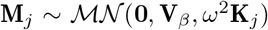 represents the combined additive effects from all other variants across the *D* traits and 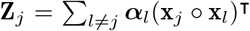 with 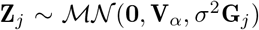 encodes all pairwise interaction terms involving the *j*-th SNP across the *D* traits. Here, the term **Z**_*j*_ becomes the main focus of model inference.

In this study, we demonstrate the utility of mvMAPIT while analyzing *D* = 2 traits at a time, but note that the framework can easily be applied to more phenotypes. Additional traits require more resources both in terms of compute time and memory. For each point estimate, mvMAPIT performs matrix operations that scale quadratically with sample size. The software also needs to store covariance matrices corresponding to the number of random effects in the model. Both these added costs scale as *D*(*D* + 1)/2 for *D* traits. When higher order interactions are included, the additional burden on resources come from requiring to store additional covariance matrices as well as projecting these covariance matrices onto the space orthogonal to the variant of interest and the population intercept. The time complexity of the projection scales as DN^2^ with again *N* being the number of samples in the data.

### Hypothesis testing in the mvMAPIT framework

The goal of identifying variants with marginal epistatic effects in the mvMAPIT framework still comes down to assessing the null hypothesis *H*_0_: *σ*^2^ =0. However, parameter estimation in mvLMMs can present substantial computational challenges. For example, one common way in the literature to rewrite the model specified in Eq. (15) is to vectorize (or stack) the columns of each matrix in the regression such that **y** = vec(**Y**), ***μ*** = vec(**U**), **m**_*j*_ = vec(**M**_*j*_), **z**_*j*_ = vec(**Z**_*j*_), and **ε** = vec(**E**). Under this reformulation, we could simply follow the procedures in Eqs. (8)–(13) to find significant variance components; but since 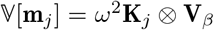 and 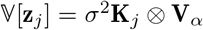 are each *ND* × *ND* dimensions (via the Kronecker product ⊗), the per-iterative computation time for performing hypothesis testing on each *j*-th SNP would now increase both with the number of individuals (*N*) and with the number of phenotypes (*D*). This could make model fitting infeasible for large biobanks with only two traits. As an alternative, we present a combinatorial approach which first fits univariate MAPIT models and then combines the resulting *P*-values with those stemming from a “covariance statistic” which looks for shared marginal epistatic effects between all pairwise combinations of the *D* traits. Importantly, our combinatorial approach does not make assumptions about the covariance structure between traits, which would need to be known (or assumed) in the Kronecker formulation.

To implement the multivariate marginal epistasis test, we follow a similar strategy used in the univariate MAPIT model and right multiply Eq. (15) by a variant-specific projection 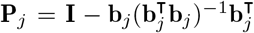 which maps the model onto a column space that is orthogonal to the population-level intercepts and the genotypic vector **x**_*j*_. This results in a simplified mvLMM of the following form

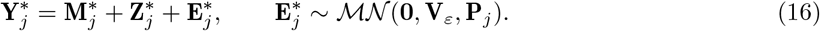

where, in addition to previous notation, 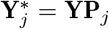; 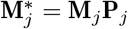; 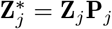, and 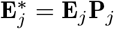, respectively. Probabilistically, this transformation assumes 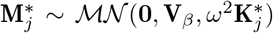 with 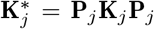; and 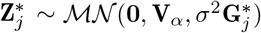 with 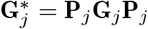. The joint analysis of multiple outcomes requires a generalization of the MQS algorithm to also include moment estimates for the covariance components between traits. Without loss of generality, we will let 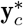 and 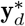 be the c-th and d-th rows of the measured phenotypic matrix 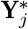, respectively. The general MQS estimates for the marginal epistatic effect is a generalization of Eq. (11) which is given in the following quadratic form

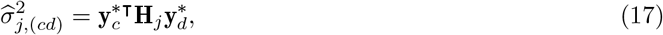

where **H**_*j*_ is as previously defined in the univariate MAPIT case and the indices span between the *c*, *d* ∈ 1,…, *D* phenotypes. Here, when *c* = *d*, the above is exactly equal to Eq. (11); however, when *c* ≠ *d*, then Eq. (17) takes on a bilinear form where 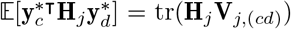 with 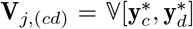 being the covariance between any two traits of interest. The corresponding standard error of the bilinear covariance component can then be estimated via the following approximation^96^

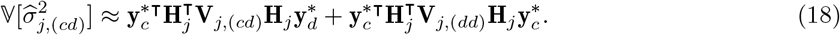

Once again, notice that when *c* = *d*, the term **V**_*j*,(*cd*)_ = **V**_*j*,(*dd*)_ and the above approximation in Eq. (18) is equal to Eq. (12).

The combinatorial hypothesis procedure that is used in mvMAPIT occurs in three key steps.

1. In the first step, the model fits univariate models for all *D* traits of interests (i.e., using Eqs. (8)–(13) from the MAPIT model or equivalently Eqs. (17) and (18) with *c* = *d*). Here, we use the proposed hybrid testing approach where we first implement a normal test by default, and then apply the exact Davies method when the *P*-value from the normal test is below the nominal significance threshold of 0.05^81^.
2. In the second step, we derive *P*-values for the covariance components (i.e., using Eqs. (17) and (18) when *c* ≠ *d*) with a normal test. As we have shown in the main text, the *P*-values derived for the covariance components with the asymptotic normal approximation tend to be slightly deflated under the null hypothesis. While this leads to generally conservative behavior with respect to type I error control, the downside is that the test may result in reduced power under the alternative, especially after multiple correction for datasets with small sample sizes or for traits that have low genetical correlation. In these cases, deriving an exact test to obtain more calibrated *P*-values could be done; however, we do not explore this line of work here.
3. In the third and final step, mvMAPIT combines the *P*-values from the first two steps into an overall marginal epistatic *P*-value. Assume that we only have *D* = 2 traits. In this case, we would have T = 3 sets of *P*-values (two marginal sets and one covariance set). The mvMAPIT software carries out the *P*-value combining procedure in two different ways. The first assumes that each of the *t* = 1,…, *T* tests are (effectively) independent and implements Fisher’s method^98^ which combines *P*-values into a single chi-square test statistic using the formula 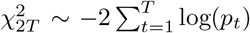 where *p_t_* denotes the *P*-value from the *t*-th test. In Fisher’s method, the χ^2^ test statistic will be large when *P*-values tend to be small (i.e., when the null hypothesis is not true for every test). The second approach assumes an unknown dependency structure between each of the *T* tests and computes a harmonic mean^99^ *P*-value where 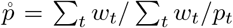. Here, 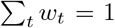 are weights which we uniformly set to be *w_t_* = 1/*T* for all *P*-values.

In practice, epistatic effects are assumed to make small contributions to the overall broad-sense heritability of complex traits^50–52^. As a result, detecting associated variants that significantly contribute to non-additive variation can be difficult. Intuitively, this combinatorial approach is meant to aggregate over the signal identified in both the marginal and covariance tests to improve power. In the main text, we show that both of Fisher’s method and the harmonic mean approach are well calibrated under the null hypothesis (i.e., only additive effects for all traits analyzed) and increase the ability to detect marginal epistatic variants under the alternative.

### Simulation studies

To test the utility of the mvMAPIT framework, we modified a frequently used simulation scheme^12,81^ to generate collections of synthetic quantitative traits under multiple genetic architectures using real genotypes from chromosome 22 of the control samples in the Wellcome Trust Case Control Consortium (WTCCC) 1 study. After preprocessing, considering this particular group of individuals and SNPs resulted in a dataset consisting of *N* = 2,938 individuals and *J* = 5,747 markers. In these simulations, we randomly choose 1,000 causal SNPs to directly affect *D* = 2 phenotypes. We generate these synthetic traits via the following general multivariate linear model:

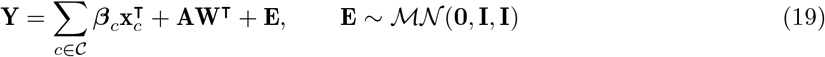

where **Y** is an *D* × *N* matrix containing all the phenotypes; 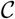 represents the set of 1,000 causal SNPs; **x**_*c*_ is the genotype for the *c*-th causal SNP encoded as 0, 1, or 2 copies of a reference allele; ***β***_*c*_ is a *D*-dimensional vector and represent the additive effect sizes for the *c*-th SNP in the *D* traits; **W** is an *N* × *M* matrix which holds pairwise interactions (i.e., Hadamard products) between some subset of causal SNPs; **A** = [***α***_1_,…, ***α***_*M*_] is a *D* × *M* matrix of interaction effect sizes with ***α***_*m*_ being *D*-dimensional epistatic coefficients for the *m*-th interaction in the *d*-th trait; and **E** is an *D* × *N* matrix of normally distributed environmental noise.

In these studies, we assume that the total phenotypic variances for both traits in **Y** are set to be 1. The additive and interaction effect sizes for causal SNPs are randomly drawn from matrix normal distributions where we control the correlation of effects between traits. This simplifies to us drawing coefficients as

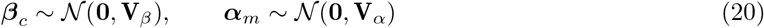

where **V**_*β*_ and **V**_*α*_ are *D* × *D* covariance matrices for additive effects and pairwise interactions between the phenotypes. Once these coefficients are sampled, we rescale them so that they explain a fixed proportion of the broad-sense heritability *H*^2^. Similarly, the environmental noise matrix is rescaled such that it explains 1 – *H*^2^. When generating synthetic traits, we assume that the additive effects make up *ρ*% of the broad-sense heritability while the pairwise interactions make up the remaining (1 – *ρ*)%. Alternatively, we say that the proportion of the heritability explained by additivity is *ρH*^2^, while the proportion of phenotypic variance explained by pairwise interactions is (1 – *ρ*)*H*^2^. Setting *ρ* =1 represents the null model where the variation of a trait is driven by solely additive effects. Here, we use the same simulation strategy used in previous studies^12,81^ where we divide the causal variants into three groups where:

- 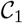 is a small number of SNPs with additive and epistatic effects;
- 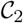 is a larger number of SNPs with additive and epistatic effects;
- 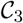 is a large number of SNPs with only additive effects.

Here, the epistatic causal SNPs interact between sets, so that SNPs in 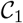 with SNPs in the 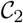, but do not interact with variants in their own group (with the same rule applies to the second group). With this set up, one can think of the SNPs assigned to 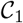 as being the “hub nodes” in an interaction network. Note that we use this setup because it has been shown that the ability to detect two interacting variants depends on the proportion of phenotypic variance that they marginally explain. For example, in our case, this means that power is expected to depend on 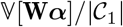 and 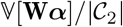 for groups 1 and 2, respectively, where 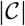 denotes the cardinality of the set. Given different parameters for the generative model in Eq. (19), we simulate data mirroring a wide range of genetic architectures by varying the following parameters:

- broad-sense heritability: *H*^2^ = 0.3 and 0.6;
- proportion of phenotypic variation that is explained by additive effects: *ρ* = 0.5, 0.8, and 1;
- causal SNPs in each of the three groups: 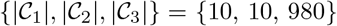 and {10, 20, 970};
- correlation between additive effects: *v*_*β*,12_ = 0, 0.8, and 1;
- correlation between epistatic effects: *v*_*α*,12_ = 0 and 0.8.

All figures and tables show the mean performances (and standard errors) for each parameter combination across 100 simulated replicates.

### Preprocessing of the heterogenous stock of mice dataset

As part of the analyses, this work makes use of GWA data from the Wellcome Trust Centre for Human Genetics^89–91^ (http://mtweb.cs.ucl.ac.uk/mus/www/mouse/index.shtml). The genotypes from this study were downloaded directly using the `BGLR-R` package ^136^. This study contains *N* = 1,814 heterogenous stock of mice from 85 families (all descending from eight inbred progenitor strains)^89,90^, and 131 quantitative traits that are classified into 6 broad categories including behavior, diabetes, asthma, immunology, haematology, and biochemistry. Phenotypic measurements for these mice can be found freely available online to download (details can be found at http://mtweb.cs.ucl.ac.uk/mus/www/mouse/HS/index.shtml and https://github.com/lcrawlab/mvMAPIT). In the main text, we focused on 15 hematological phenotypes including: atypical lymphocytes (ALY; `Haem.ALYabs`), basophils (BAS; `Haem.BASabs`), hematocrit (HCT; `Haem.HCT`), hemoglobin (HGB; `Haem.HGB`), large immature cells (LIC; `Haem.LICabs`), lymphocytes (LYM; `Haem.LYMabs`), mean corpuscular hemoglobin (MCH; `Haem.MCH`), mean corpuscular volume (MCV; `Haem.MCV`), monocytes (MON; `Haem.MONabs`), mean platelet volume (MPV; `Haem.MPV`), neutrophils (NEU; `Haem.NEUabs`), plateletcrit (PCT; `Haem.PCT`), platelets (PLT; `Haem.PLT`), red blood cell count (RBC; `Haem.RBC`), red cell distribution width (RDW; `Haem.RDW`), and white blood cell count (WBC; `Haem.WBC`). All phenotypes were previously corrected for sex, age, body weight, season, year, and cage effects^89,90^. For individuals with missing genotypes, we imputed values by the mean genotype of that SNP in their corresponding family. Only polymorphic SNPs with minor allele frequency above 5% were kept for the analyses. This left a total of *J* = 10,227 autosomal SNPs that were available for all mice.

### Data and software availability

Source code, tutorials, and tutorials for implementing the “multivariate MArginal ePIstasis Test” are publicly available as an R package which is available online at https://github.com/lcrawlab/mvMAPIT. We use the `CompQuadForm` R package^137^ to compute *P*-values from the Davies method. The Davies method can sometimes yield a *P*-value equal exactly to 0 when the true *P*-value is extremely small^137^. If this is of concern, one can compute the *P*-values for MAPIT using Kuonen’s saddlepoint method^138^ or Satterthwaite’s approximation equation^139^. In the current implementation of mvMAPIT, the saddlepoint approximation is performed if the Davies method returns with error. We wrote our own function to combine *P*-values using Fisher’s method which is largely inspired by functions in the `metap` R package^140^. We use the `harmonicmeanp` R package^141,142^ to combine *P*-values using the harmonic mean. Full package documentation can be found at https://lcrawlab.github.io/mvMAPIT/. Data to reproduce figures for the broadly neutralizing antibodies as well as the mice study can be found at https://doi.org/10.7910/DVN/WPFIGU^143^.

Data about the binding affinity landscapes for neutralizing antibodies were downloaded directly from Phillips et al.^88^. Information about mice dataset from the Wellcome Trust Centre for Human Genetics^89–91^ can be found at http://mtweb.cs.ucl.ac.uk/mus/www/mouse/index.shtml. The genotypes from this study were downloaded using the `BGLR-R` package ^136^. Details about the mice phenotypes can be found http://mtweb.cs.ucl.ac.uk/mus/www/mouse/HS/index.shtml and hematological traits can be downloaded from the mvMAPIT package. In the real data analyses, SNPs were mapped to the closest neighboring genes using the Mouse Genome Informatics database (http://www.informatics.jax.org)^106^.

## Supplementary Figures

**Figure S1.**
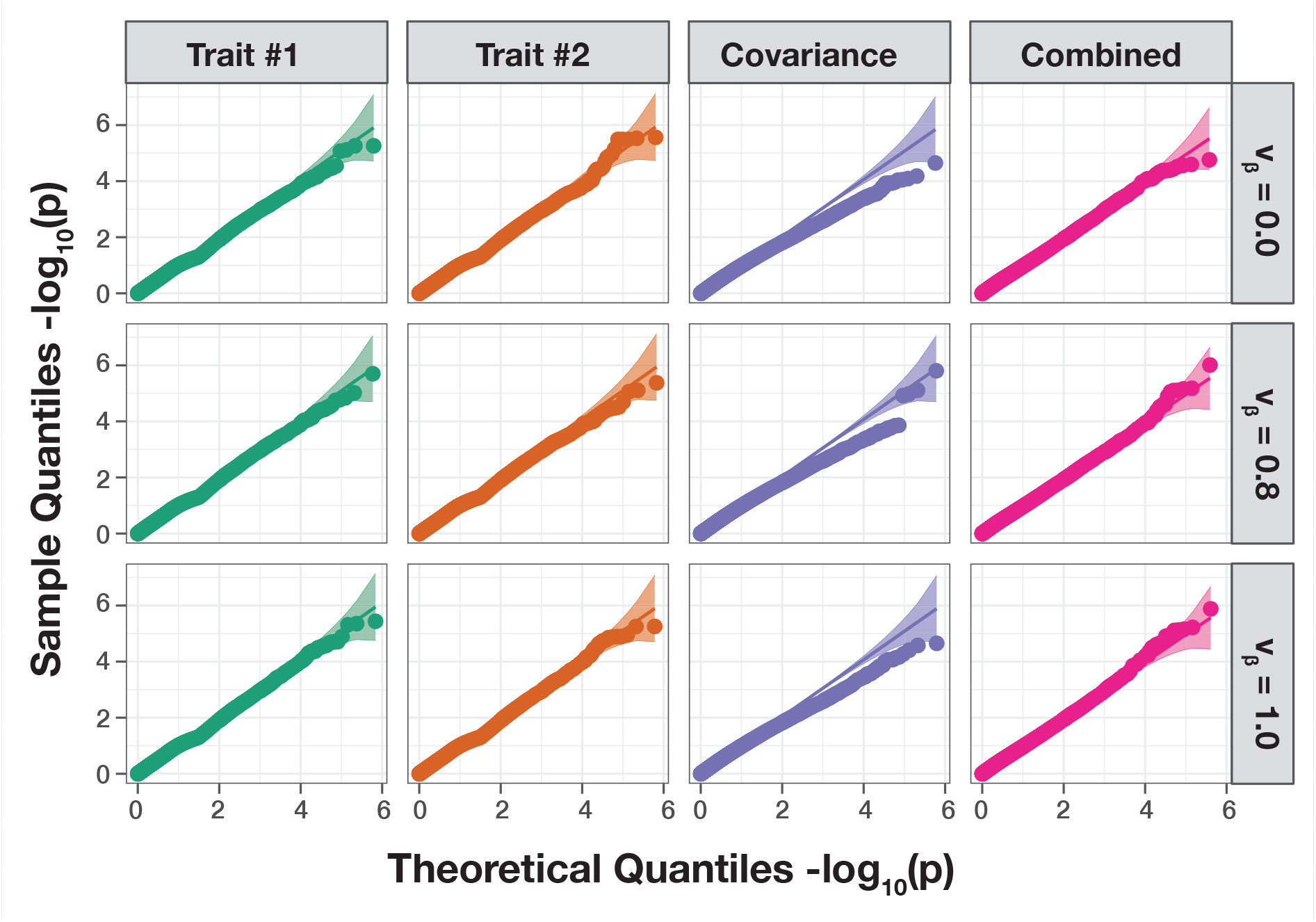
The mvMAPIT framework using Fisher’s method produces well-calibrated *P*-values when traits are generated by only additive effects (sample size *N* = 1,000 individuals). In these simulations, quantitative traits are simulated to have narrow-sense heritability *h*^2^ = 0.6 with an architecture made up of only additive genetic variation. Each row of quantile-quantile (QQ) plots corresponds to a setting where the additive genetic effects for a causal SNP have different correlation structures across traits. In these simulations, we consider scenarios where we have traits with independent additive effects (*v_β_* = 0), traits with highly correlated additive effects (*v_β_* = 0.8), and traits with perfectly correlated additive effects (*v_β_* = 1). The first two columns show *P*-values resulting from the univariate MAPIT test on “trait #1” and “trait #2”, respectively. The third column depicts the “covariance” *P*-values which corresponds to assessing the pairwise interactions affecting both traits is. Lastly, the fourth column shows the final “combined” *P*-value which combines the *P*-values from the first three columns using Fisher’s method. The 95% confidence interval for the null hypothesis of no marginal epistatic effects is shown in grey. Each plot combines results from 100 simulated replicates.

**Figure S2.**
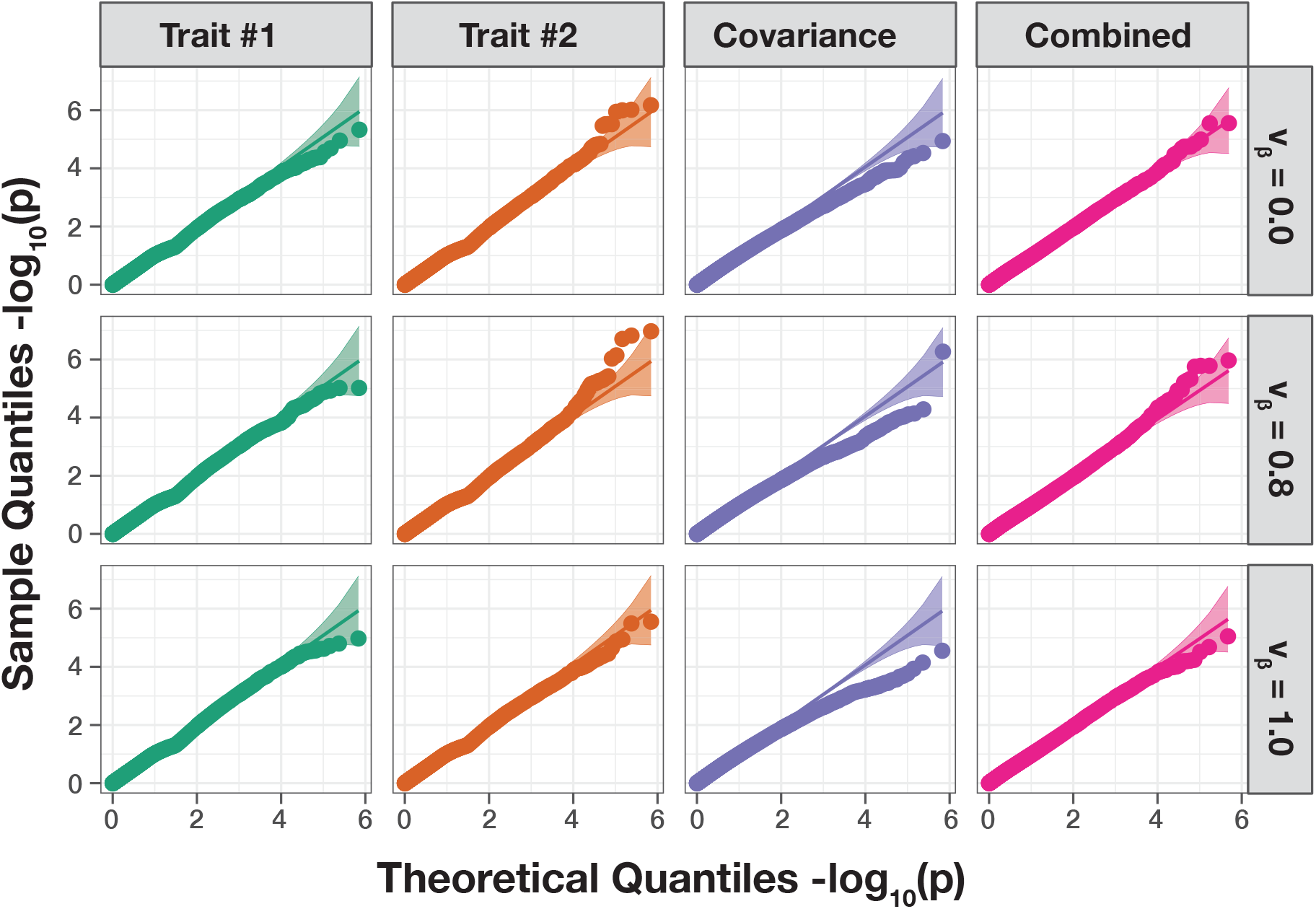
The mvMAPIT framework using Fisher’s method produces well-calibrated *P*-values when traits are generated by only additive effects (sample size *N* = 1,750 individuals). In these simulations, quantitative traits are simulated to have narrow-sense heritability *h*^2^ = 0.6 with an architecture made up of only additive genetic variation. Each row of quantile-quantile (QQ) plots corresponds to a setting where the additive genetic effects for a causal SNP have different correlation structures across traits. In these simulations, we consider scenarios where we have traits with independent additive effects (*v_β_* = 0), traits with highly correlated additive effects (*v_β_* = 0.8), and traits with perfectly correlated additive effects (*v_β_* = 1). The first two columns show *P*-values resulting from the univariate MAPIT test on “trait #1” and “trait #2”, respectively. The third column depicts the “covariance” *P*-values which corresponds to assessing the pairwise interactions affecting both traits is. Lastly, the fourth column shows the final “combined” *P*-value which combines the *P*-values from the first three columns using Fisher’s method. The 95% confidence interval for the null hypothesis of no marginal epistatic effects is shown in grey. Each plot combines results from 100 simulated replicates.

**Figure S3.**
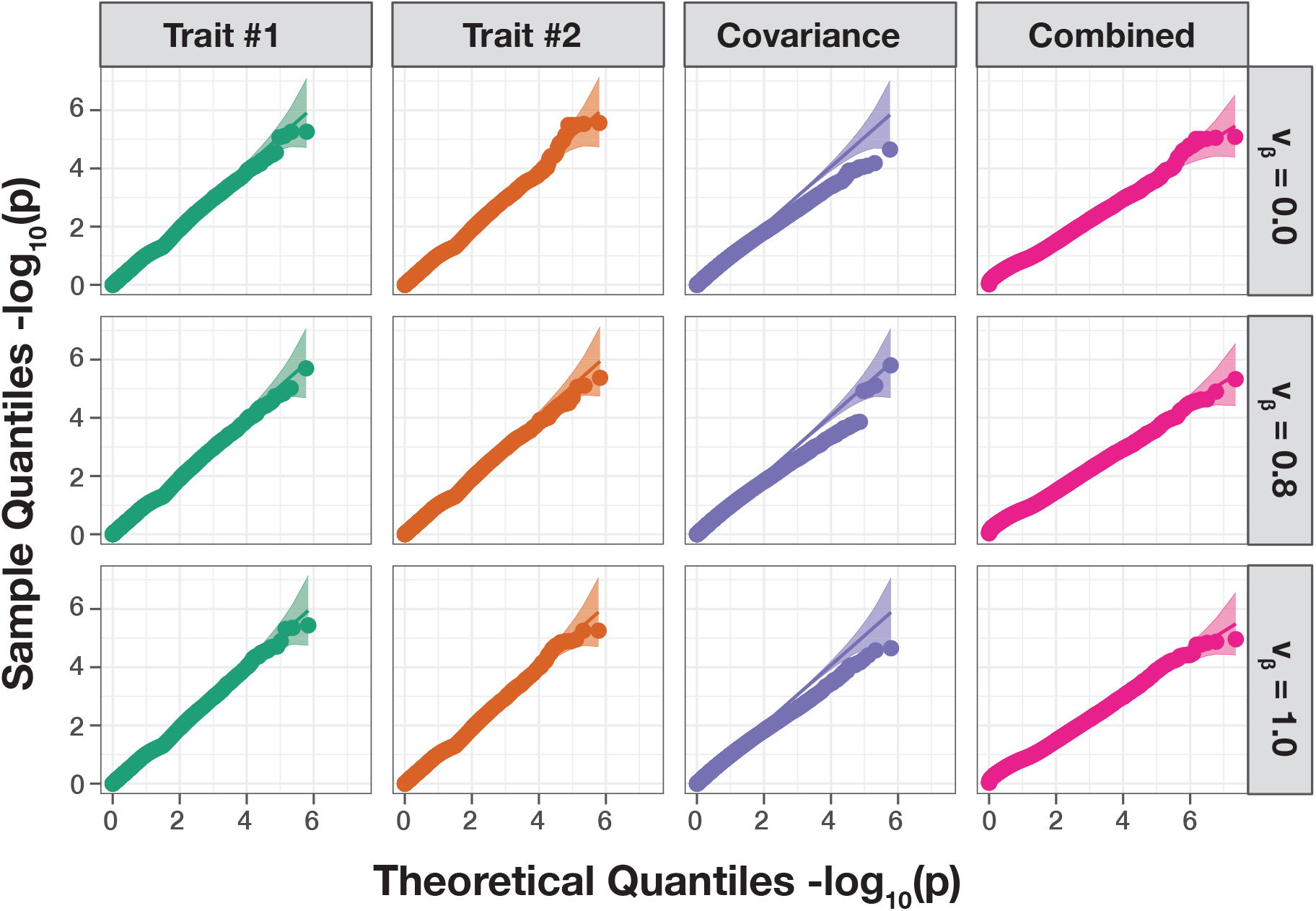
The mvMAPIT framework using the harmonic mean produces well-calibrated *P*-values when traits are generated by only additive effects (sample size *N* = 1,000 individuals). In these simulations, quantitative traits are simulated to have narrow-sense heritability *h*^2^ = 0.6 with an architecture made up of only additive genetic variation. Each row of quantile-quantile (QQ) plots corresponds to a setting where the additive genetic effects for a causal SNP have different correlation structures across traits. In these simulations, we consider scenarios where we have traits with independent additive effects (*v_β_* = 0), traits with highly correlated additive effects (*v_β_* = 0.8), and traits with perfectly correlated additive effects (*v_β_* = 1). The first two columns show *P*-values resulting from the univariate MAPIT test on “trait #1” and “trait #2”, respectively. The third column depicts the “covariance” *P*-values which corresponds to assessing the pairwise interactions affecting both traits is. Lastly, the fourth column shows the final “combined” *P*-value which combines the *P*-values from the first three columns using Fisher’s method. The 95% confidence interval for the null hypothesis of no marginal epistatic effects is shown in grey. Each plot combines results from 100 simulated replicates.

**Figure S4.**
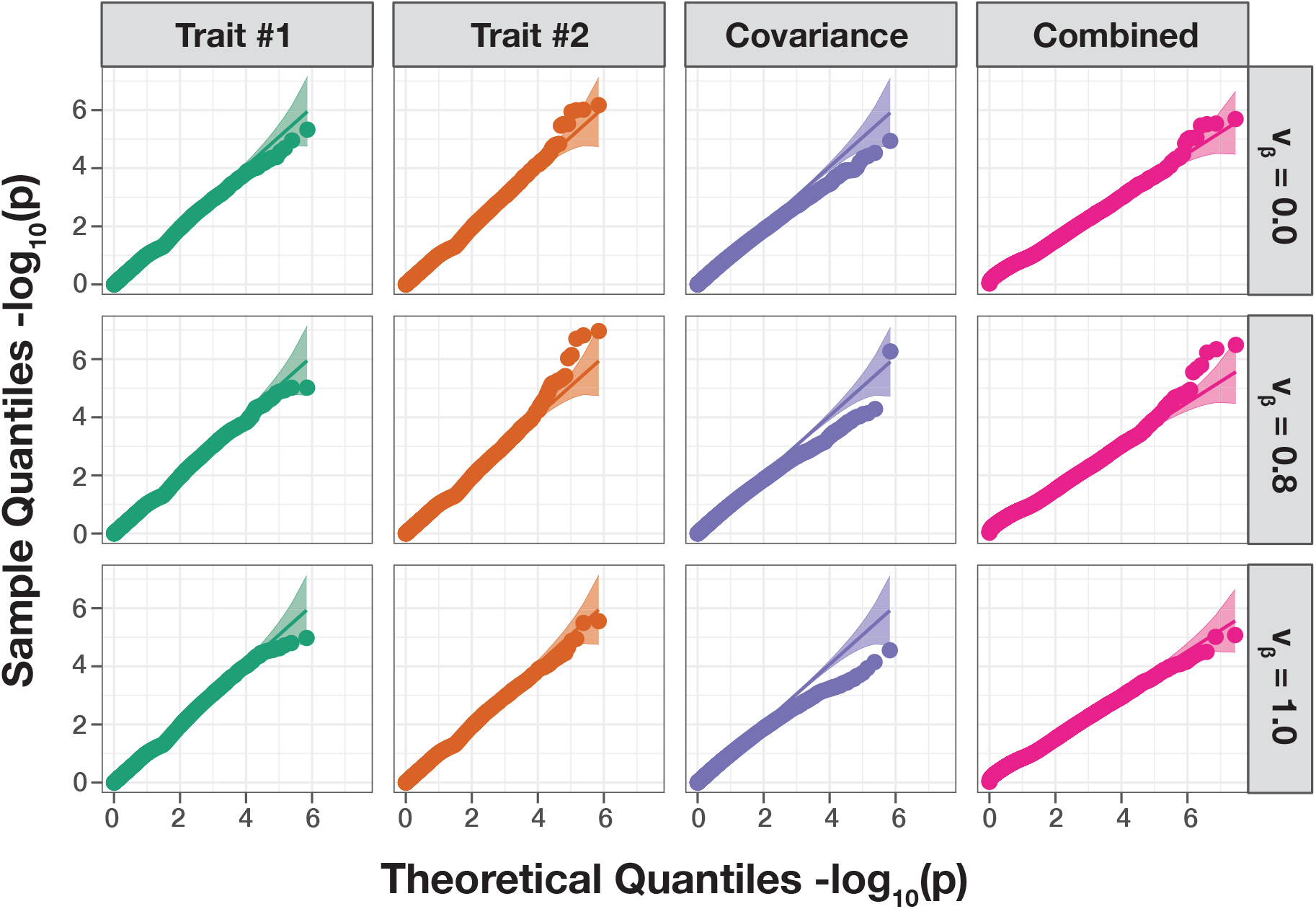
The mvMAPIT framework using the harmonic mean produces well-calibrated *P*-values when traits are generated by only additive effects (sample size *N* = 1,750 individuals). In these simulations, quantitative traits are simulated to have narrow-sense heritability *h*^2^ = 0.6 with an architecture made up of only additive genetic variation. Each row of quantile-quantile (QQ) plots corresponds to a setting where the additive genetic effects for a causal SNP have different correlation structures across traits. In these simulations, we consider scenarios where we have traits with independent additive effects (*v_β_* = 0), traits with highly correlated additive effects (*v_β_* = 0.8), and traits with perfectly correlated additive effects (*v_β_* = 1). The first two columns show *P*-values resulting from the univariate MAPIT test on “trait #1” and “trait #2”, respectively. The third column depicts the “covariance” *P*-values which corresponds to assessing the pairwise interactions affecting both traits is. Lastly, the fourth column shows the final “combined” *P*-value which combines the *P*-values from the first three columns using Fisher’s method. The 95% confidence interval for the null hypothesis of no marginal epistatic effects is shown in grey. Each plot combines results from 100 simulated replicates.

**Figure S5.**
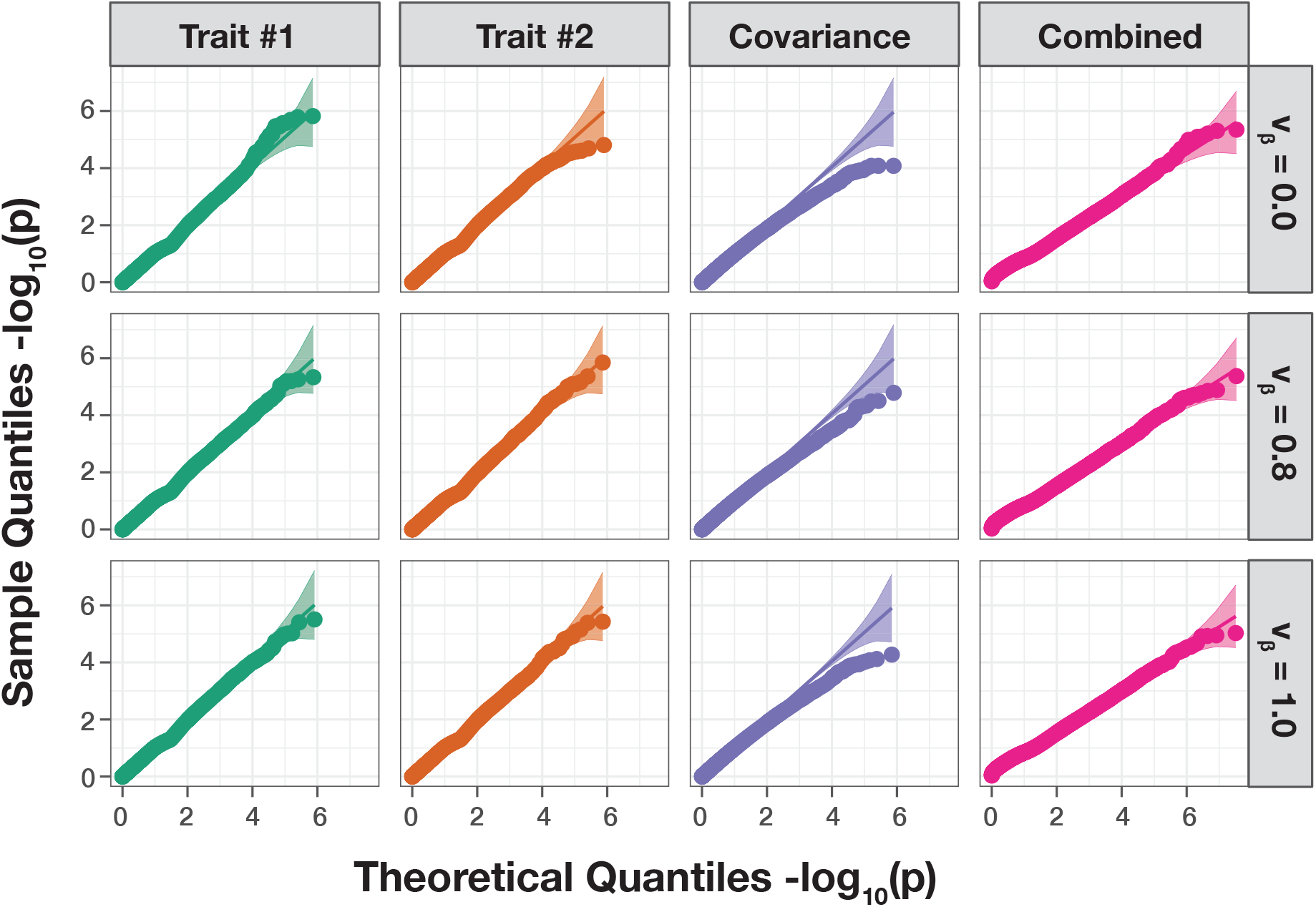
The mvMAPIT framework using the harmonic mean produces well-calibrated *P*-values when traits are generated by only additive effects (sample size *N* = 2,500 individuals). In these simulations, quantitative traits are simulated to have narrow-sense heritability *h*^2^ = 0.6 with an architecture made up of only additive genetic variation. Each row of quantile-quantile (QQ) plots corresponds to a setting where the additive genetic effects for a causal SNP have different correlation structures across traits. In these simulations, we consider scenarios where we have traits with independent additive effects (*v_β_* = 0), traits with highly correlated additive effects (*v_β_* = 0.8), and traits with perfectly correlated additive effects (*v_β_* = 1). The first two columns show *P*-values resulting from the univariate MAPIT test on “trait #1” and “trait #2”, respectively. The third column depicts the “covariance” *P*-values which corresponds to assessing the pairwise interactions affecting both traits is. Lastly, the fourth column shows the final “combined” *P*-value which combines the *P*-values from the first three columns using Fisher’s method. The 95% confidence interval for the null hypothesis of no marginal epistatic effects is shown in grey. Each plot combines results from 100 simulated replicates.

**Figure S6.**
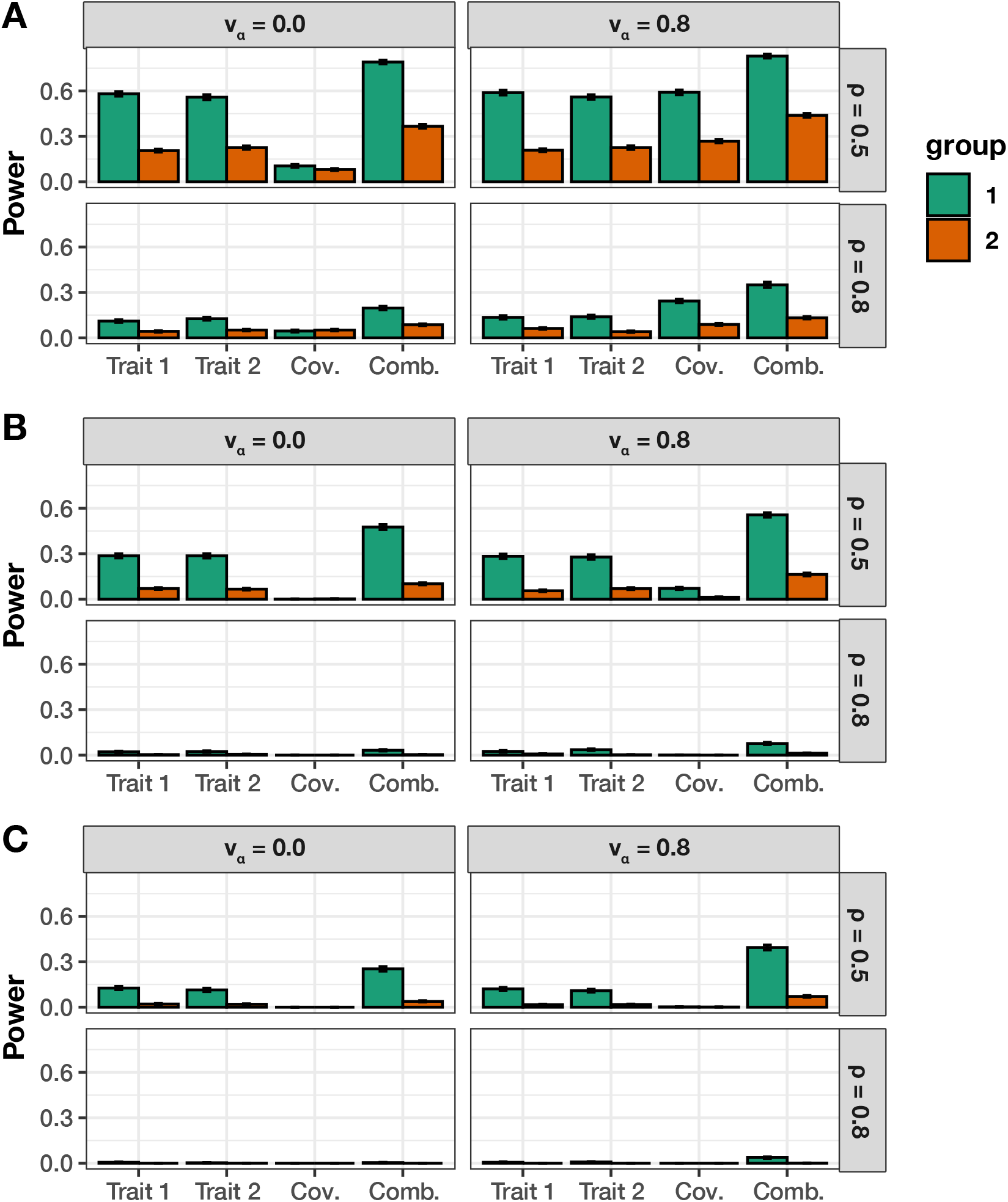
Empirical power of mvMAPIT with Fisher’s method to detect group #1 (10) and group #2 (20) epistatic variants across complex traits with moderate broad-sense heritability. In these simulations, both quantitative traits are simulated to have broad-sense heritability *H*^2^ = 0.6 with architectures made up of both additive and epistatic effects. The parameter *ρ* = {0.5, 0.8} is used to determine the portion of broad-sense heritability contributed by additive effects. Each column corresponds to a setting where the epistatic effects for interactive pairs have different correlation structures across traits. In these simulations, we consider scenarios where we have traits with independent epistatic effects (*v_α_* = 0) and traits with highly correlated epistatic effects (*v_α_* = 0.8). This plot shows the empirical power of mvMAPIT at significance levels **(A)** *P* = 5 × 10^−2^, **(B)** *P* = 5 × 10^−4^, and **(C)** *P* = 1 × 10^−5^, respectively. Group #1 and #2 causal markers are colored in green and orange, respectively. For comparison, the “trait #1” and “trait #2” bars correspond to the univariate MAPIT model, the “cov” bars corresponds to power contributed by the covariance test, and “comb” details power from the overall association identified by mvMAPIT in the combination approach. Results are based on 100 simulations per parameter combination and the horizontal bars represent standard errors.

**Figure S7.**
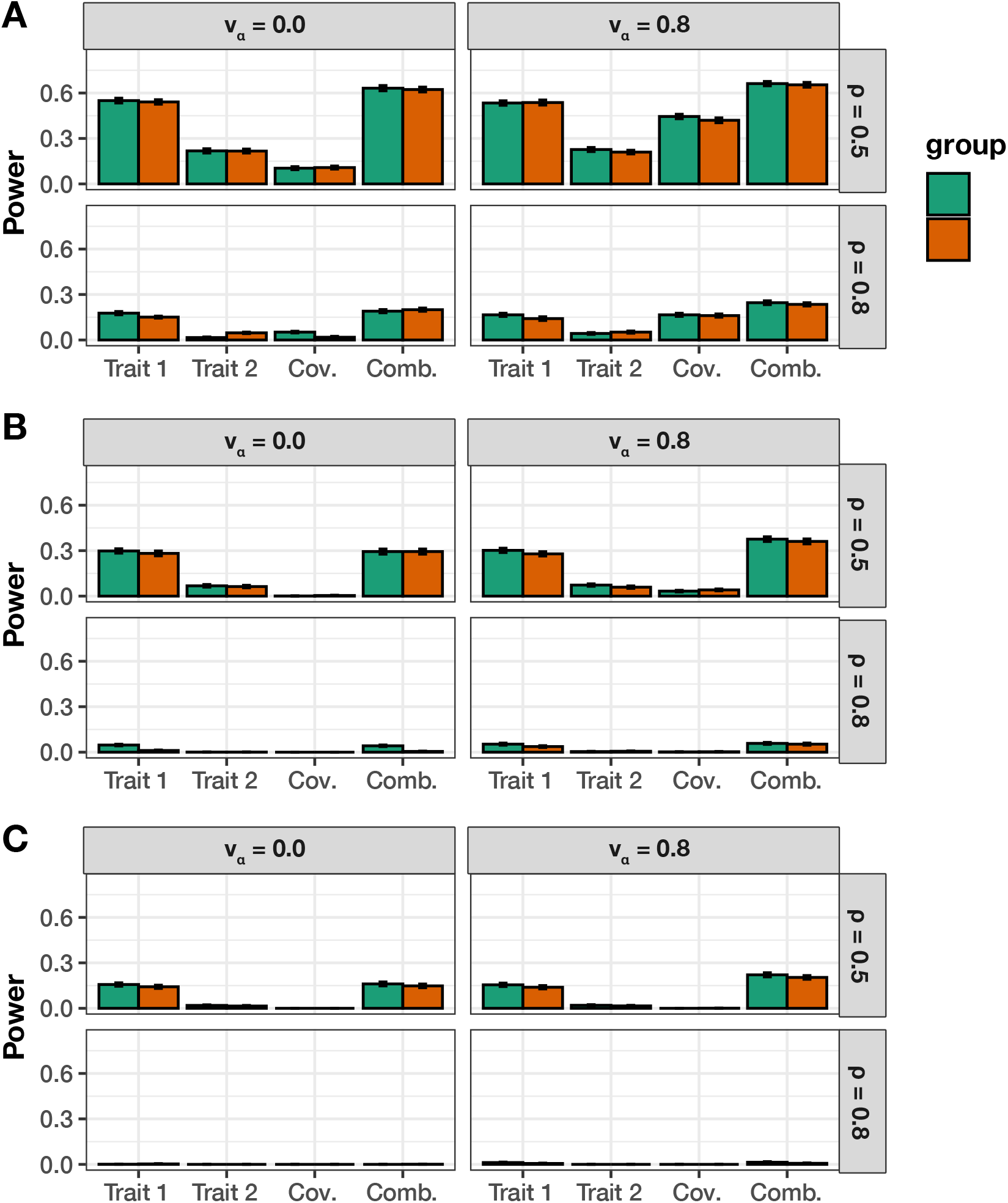
Empirical power of mvMAPIT with Fisher’s method to detect group #1 (10) and group #2 (10) epistatic variants across complex traits with different levels of broad-sense heritability. In these simulations, one of the quantitative traits has a moderate broad-sense heritability *H*^2^ = 0.6, while the other has heritability *H*^2^ = 0.3. Both traits have architectures made up of both additive and epistatic effects. The parameter *ρ* = {0.5, 0.8} is used to determine the portion of broad-sense heritability contributed by additive effects. Each column corresponds to a setting where the epistatic effects for interactive pairs have different correlation structures across traits. In these simulations, we consider scenarios where we have traits with independent epistatic effects (*v_α_* = 0) and traits with highly correlated epistatic effects (*v_α_* = 0.8). This plot shows the empirical power of mvMAPIT at significance levels **(A)** *P* = 5 × 10^−2^, **(B)** *P* = 5 × 10^−4^, and **(C)** *P* = 1 × 10^−5^, respectively. Group #1 and #2 causal markers are colored in green and orange, respectively. For comparison, the “trait #1” and “trait #2” bars correspond to the univariate MAPIT model, the “cov” bars corresponds to power contributed by the covariance test, and “comb” details power from the overall association identified by mvMAPIT in the combination approach. Results are based on 100 simulations per parameter combination and the horizontal bars represent standard errors.

**Figure S8.**
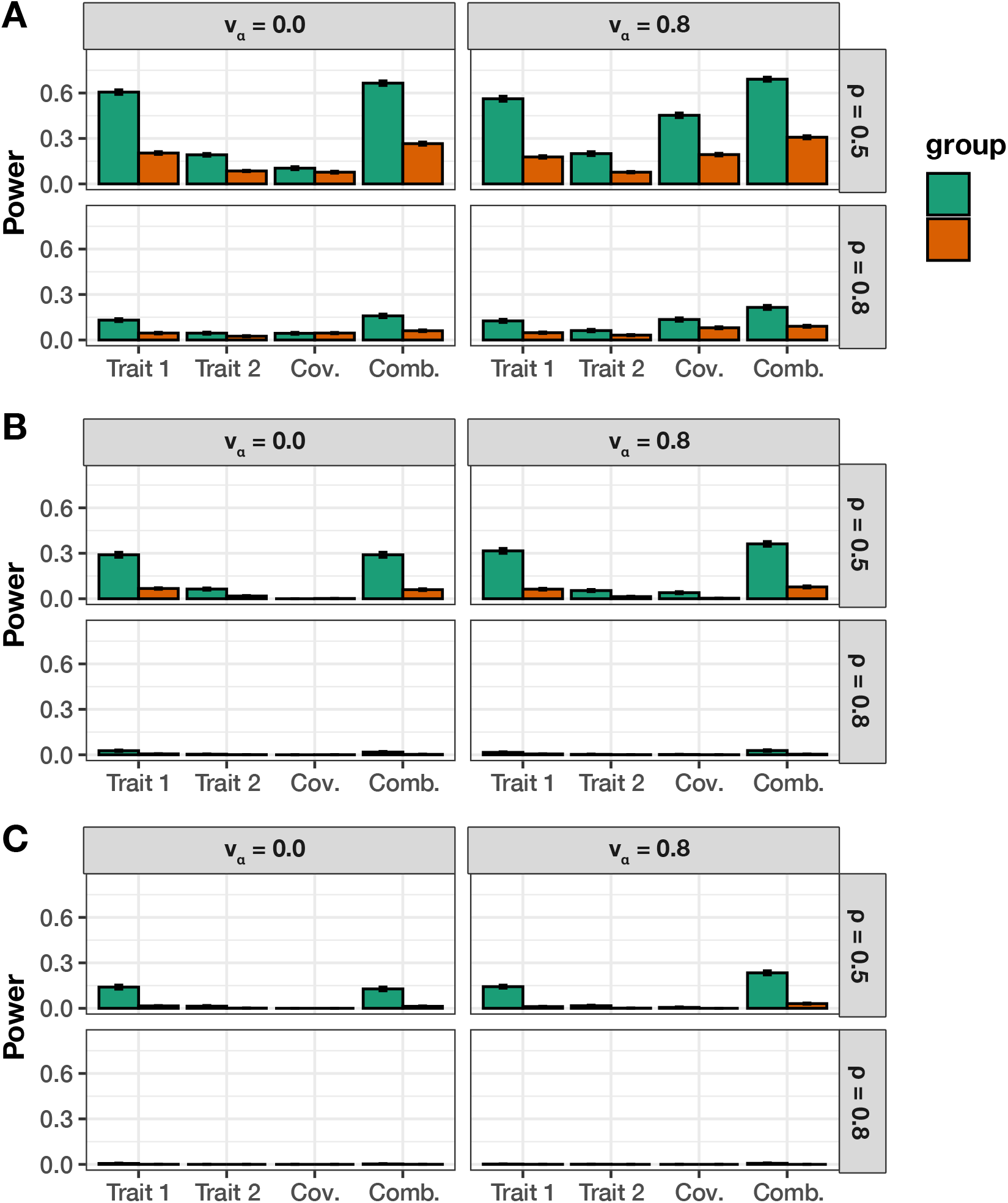
Empirical power of mvMAPIT with Fisher’s method to detect group #1 (10) and group #2 (20) epistatic variants across complex traits with different levels of broad-sense heritability. In these simulations, one of the quantitative traits has a moderate broad-sense heritability *H*^2^ = 0.6, while the other has heritability *H*^2^ = 0.3. Both traits have architectures made up of both additive and epistatic effects. The parameter *ρ* = {0.5, 0.8} is used to determine the portion of broad-sense heritability contributed by additive effects. Each column corresponds to a setting where the epistatic effects for interactive pairs have different correlation structures across traits. In these simulations, we consider scenarios where we have traits with independent epistatic effects (*v_α_* = 0) and traits with highly correlated epistatic effects (*v_α_* = 0.8). This plot shows the empirical power of mvMAPIT at significance levels **(A)** *P* = 5 × 10^−2^, **(B)** *P* = 5 × 10^−4^, and **(C)** *P* = 1 × 10^−5^, respectively. Group #1 and #2 causal markers are colored in green and orange, respectively. For comparison, the “trait #1” and “trait #2” bars correspond to the univariate MAPIT model, the “cov” bars corresponds to power contributed by the covariance test, and “comb” details power from the overall association identified by mvMAPIT in the combination approach. Results are based on 100 simulations per parameter combination and the horizontal bars represent standard errors.

**Figure S9.**
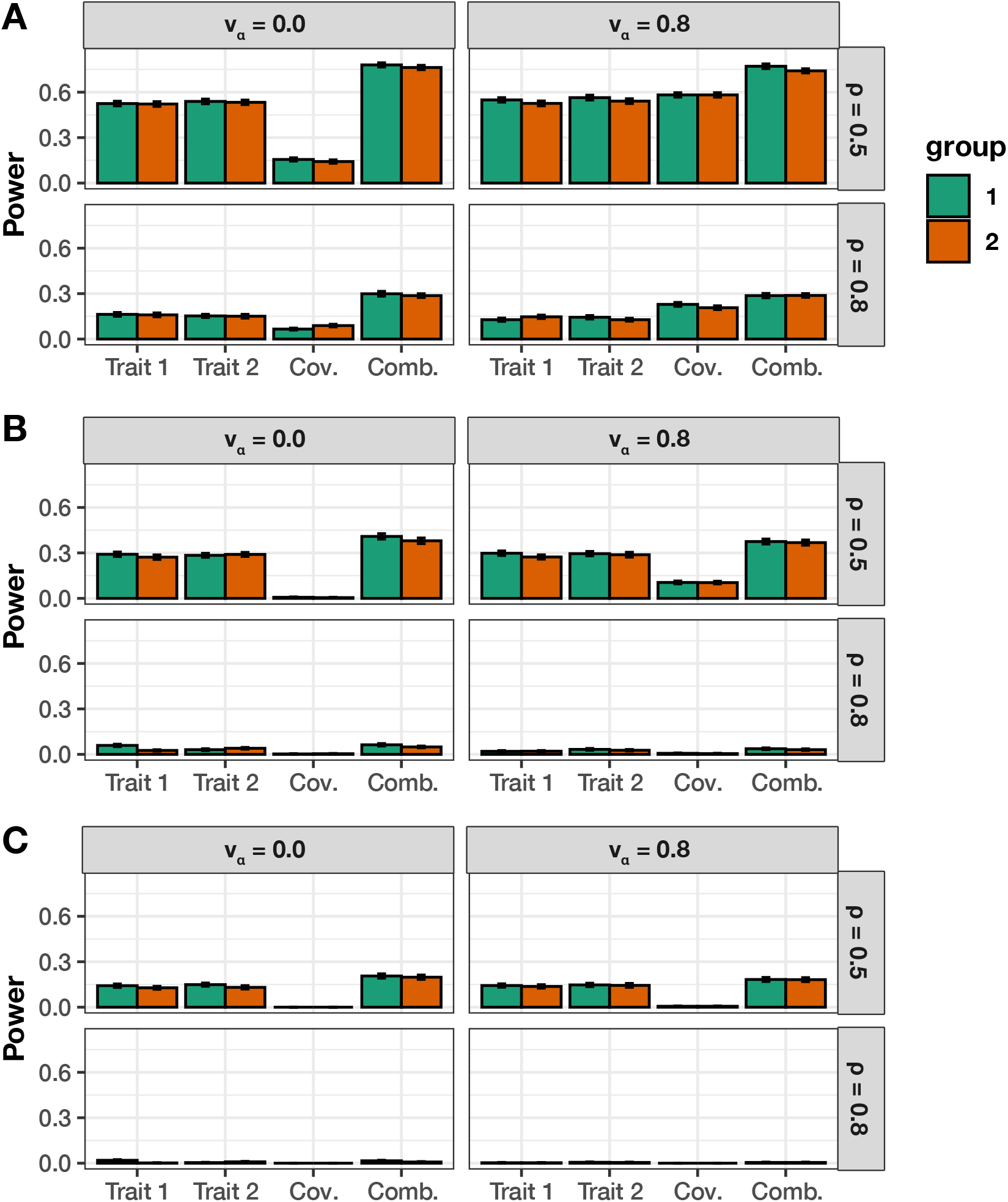
Empirical power of mvMAPIT with the harmonic mean combination approach to detect group #1 (10) and group #2 (10) epistatic variants across complex traits with moderate broad-sense heritability. In these simulations, both quantitative traits are simulated to have broad-sense heritability *H*^2^ = 0.6 with architectures made up of both additive and epistatic effects. The parameter ρ = {0.5, 0.8} is used to determine the portion of broad-sense heritability contributed by additive effects. Each column corresponds to a setting where the epistatic effects for interactive pairs have different correlation structures across traits. In these simulations, we consider scenarios where we have traits with independent epistatic effects (*v_α_* = 0) and traits with highly correlated epistatic effects (*v_α_* = 0.8). This plot shows the empirical power of mvMAPIT at significance levels **(A)** *P* = 5 × 10^−2^, **(B)** *P* = 5 × 10 ^4^, and **(C)** *P* = 1 × 10 ^5^, respectively. Group #1 and #2 causal markers are colored in green and orange, respectively. For comparison, the “trait #1” and “trait #2” bars correspond to the univariate MAPIT model, the “cov” bars corresponds to power contributed by the covariance test, and “comb” details power from the overall association identified by mvMAPIT in the combination approach. Results are based on 100 simulations per parameter combination and the horizontal bars represent standard errors.

**Figure S10.**
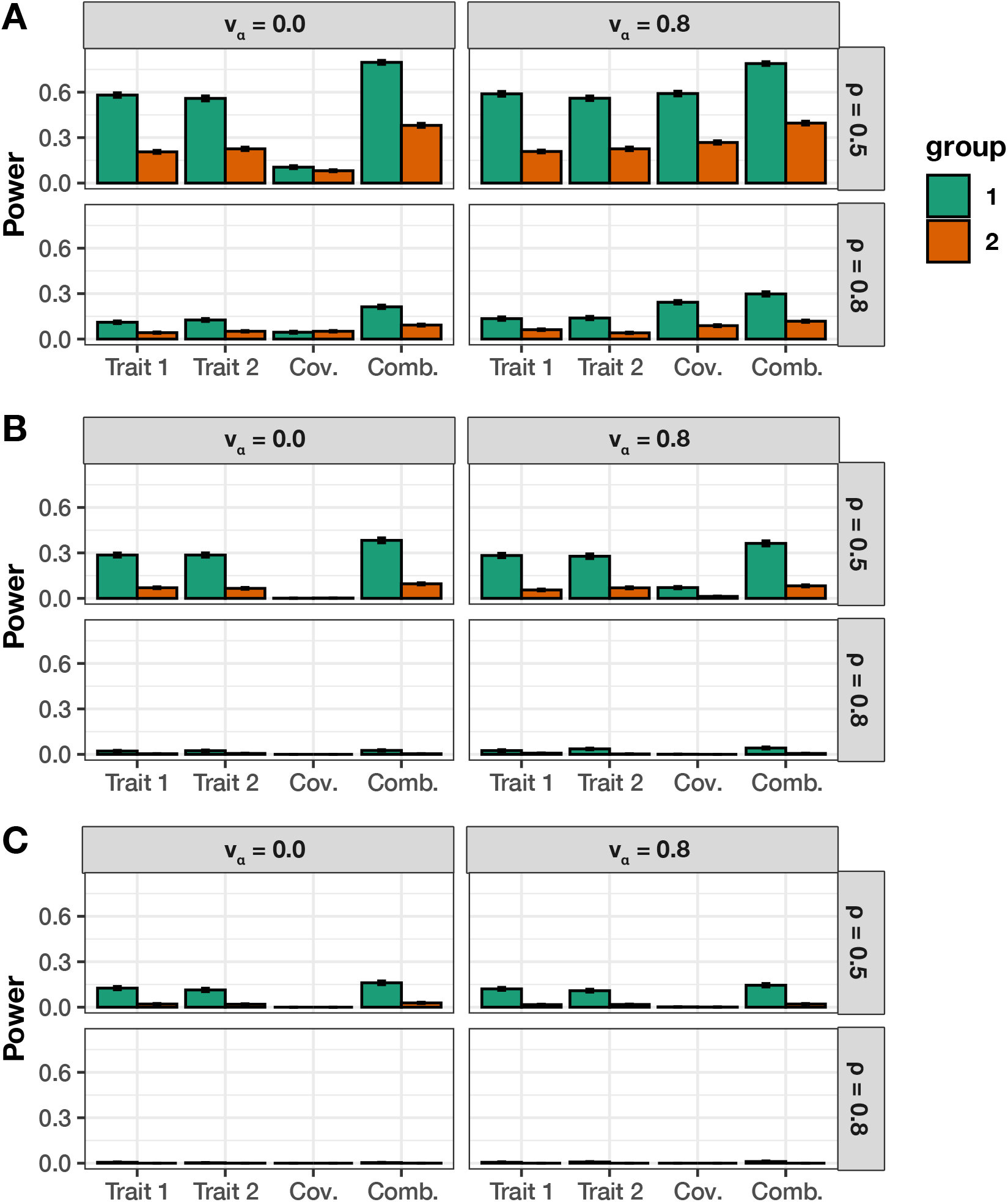
Empirical power of mvMAPIT with the harmonic mean combination approach to detect group #1 (10) and group #2 (20) epistatic variants across complex traits with moderate broad-sense heritability. In these simulations, both quantitative traits are simulated to have broad-sense heritability *H*^2^ = 0.6 with architectures made up of both additive and epistatic effects. The parameter *ρ* = {0.5, 0.8} is used to determine the portion of broad-sense heritability contributed by additive effects. Each column corresponds to a setting where the epistatic effects for interactive pairs have different correlation structures across traits. In these simulations, we consider scenarios where we have traits with independent epistatic effects (*v_α_* = 0) and traits with highly correlated epistatic effects (*v_α_* = 0.8). This plot shows the empirical power of mvMAPIT at significance levels **(A)** *P* = 5 × 10^−2^, **(B)** *P* = 5 × 10 ^4^, and **(C)** *P* = 1 × 10 ^5^, respectively. Group #1 and #2 causal markers are colored in green and orange, respectively. For comparison, the “trait #1” and “trait #2” bars correspond to the univariate MAPIT model, the “cov” bars corresponds to power contributed by the covariance test, and “comb” details power from the overall association identified by mvMAPIT in the combination approach. Results are based on 100 simulations per parameter combination and the horizontal bars represent standard errors.

**Figure S11.**
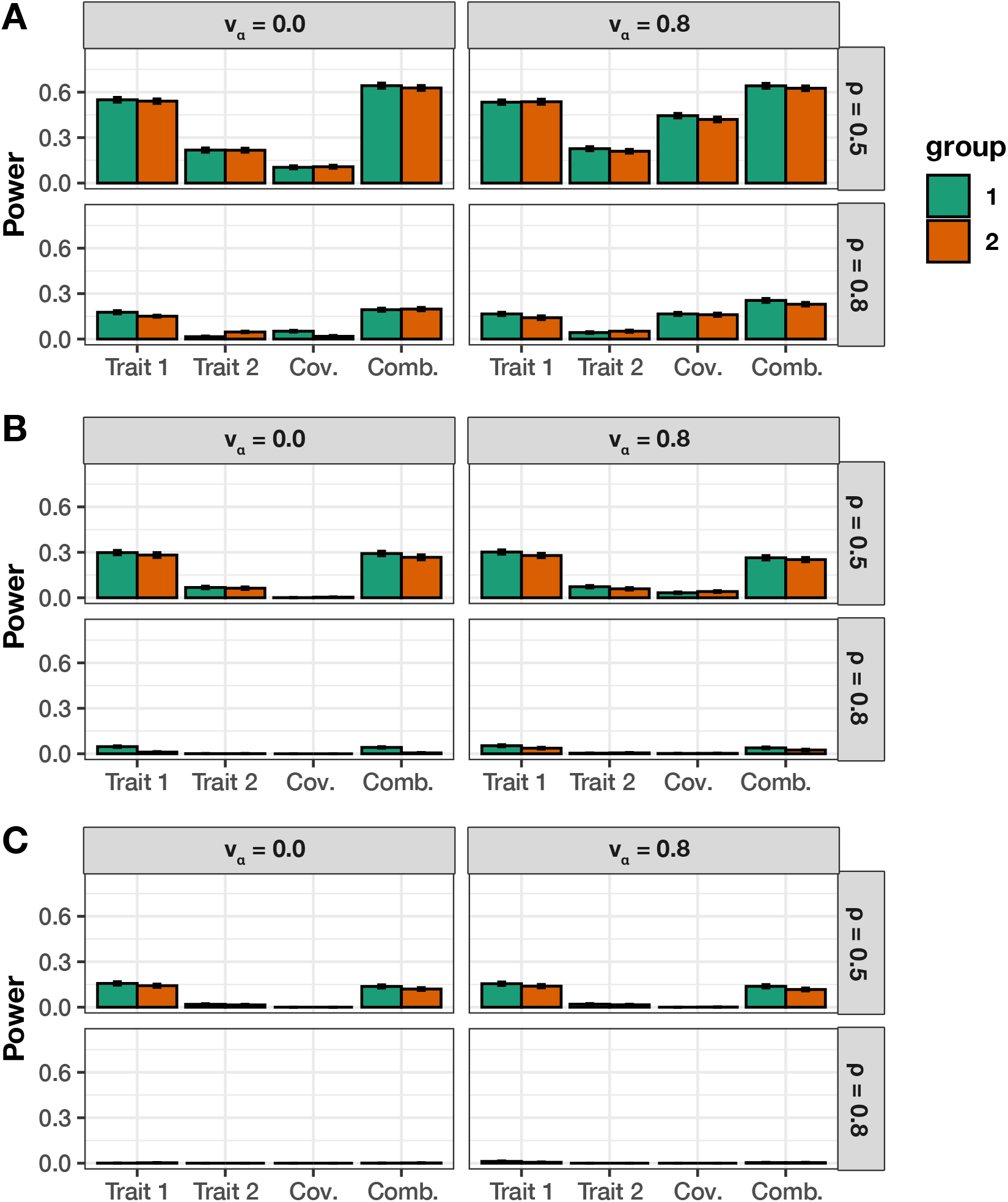
Empirical power of mvMAPIT with the harmonic mean combination approach to detect group #1 (10) and group #2 (10) epistatic variants across complex traits with different levels of broad-sense heritability. In these simulations, one of the quantitative traits has a moderate broad-sense heritability *H*^2^ = 0.6, while the other has heritability *H*^2^ = 0.3. Both traits have architectures made up of both additive and epistatic effects. The parameter *ρ* = {0.5, 0.8} is used to determine the portion of broad-sense heritability contributed by additive effects. Each column corresponds to a setting where the epistatic effects for interactive pairs have different correlation structures across traits. In these simulations, we consider scenarios where we have traits with independent epistatic effects (*v_α_* = 0) and traits with highly correlated epistatic effects (*v_α_* = 0.8). This plot shows the empirical power of mvMAPIT at significance levels **(A)** *P* = 5 × 10^−2^, **(B)** *P* = 5 × 10^−4^, and **(C)** *P* = 1 × 10^−5^, respectively. Group #1 and #2 causal markers are colored in green and orange, respectively. For comparison, the “trait #1” and “trait #2” bars correspond to the univariate MAPIT model, the “cov” bars corresponds to power contributed by the covariance test, and “comb” details power from the overall association identified by mvMAPIT in the combination approach. Results are based on 100 simulations per parameter combination and the horizontal bars represent standard errors.

**Figure S12.**
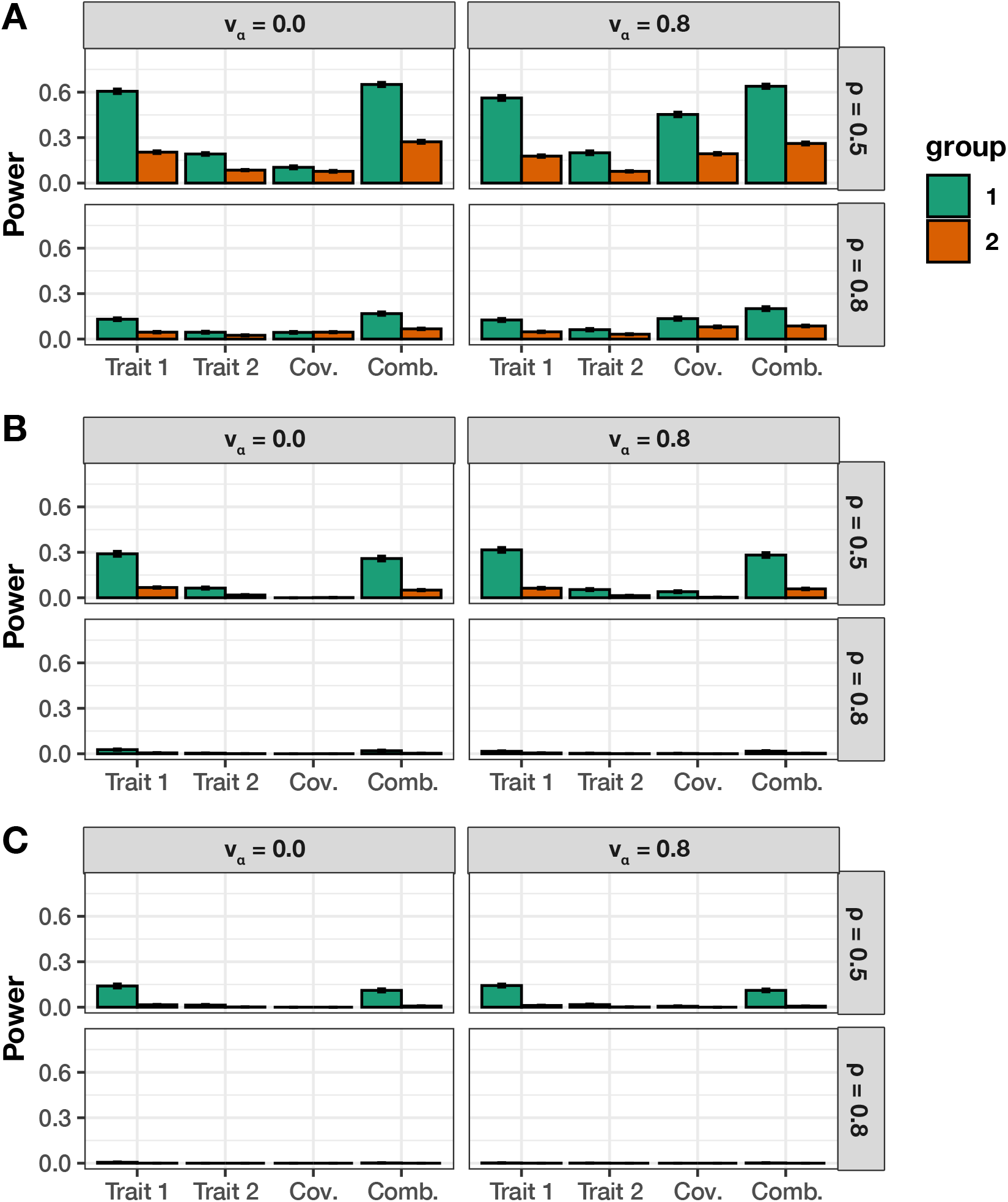
Empirical power of mvMAPIT with the harmonic mean combination approach to detect group #1 (10) and group #2 (20) epistatic variants across complex traits with different levels of broad-sense heritability. In these simulations, one of the quantitative traits has a moderate broad-sense heritability *H*^2^ = 0.6, while the other has heritability *H*^2^ = 0.3. Both traits have architectures made up of both additive and epistatic effects. The parameter *ρ* = {0.5, 0.8} is used to determine the portion of broad-sense heritability contributed by additive effects. Each column corresponds to a setting where the epistatic effects for interactive pairs have different correlation structures across traits. In these simulations, we consider scenarios where we have traits with independent epistatic effects (*v_α_* = 0) and traits with highly correlated epistatic effects (*v_α_* = 0.8). This plot shows the empirical power of mvMAPIT at significance levels **(A)** *P* = 5 × 10^−2^, **(B)** *P* = 5 × 10^−4^, and **(C)** *P* = 1 × 10^−5^, respectively. Group #1 and #2 causal markers are colored in green and orange, respectively. For comparison, the “trait #1” and “trait #2” bars correspond to the univariate MAPIT model, the “cov” bars corresponds to power contributed by the covariance test, and “comb” details power from the overall association identified by mvMAPIT in the combination approach. Results are based on 100 simulations per parameter combination and the horizontal bars represent standard errors.

**Figure S13.**
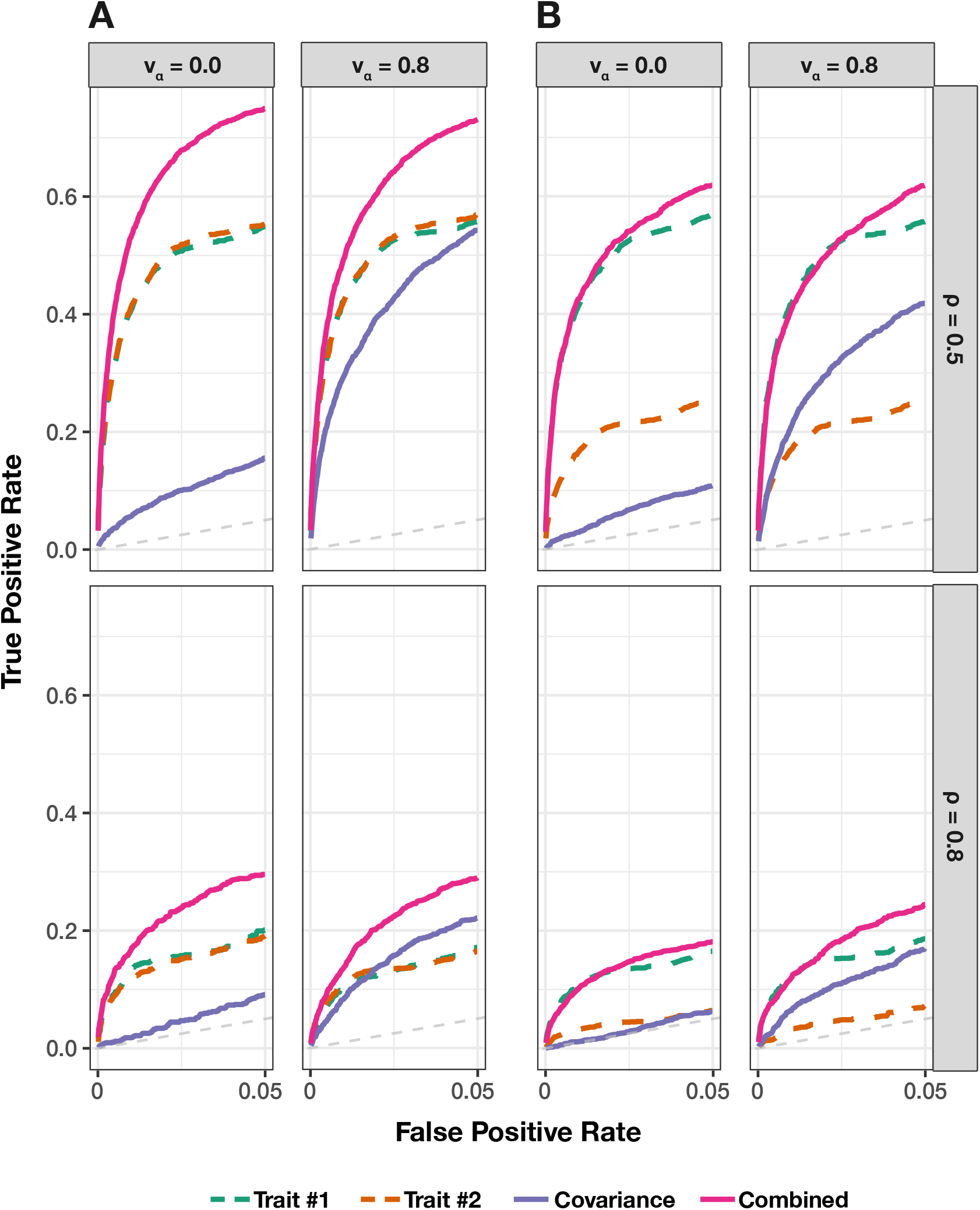
Receiver operating characteristic (ROC) curves comparing the ability of mvMAPIT using the harmonic mean to the univariate MAPIT model in detecting group #1 (10) and group #2 (10) epistatic variants across complex traits. In panel **(A)** both traits have broad-sense heritability *H*^2^ = 0.6; while in panel **(B)** one of traits has broad-sense heritability *H*^2^ = 0.6 and the other has heritability *H*^2^ = 0.3. Across the rows, the parameter *ρ* = {0.5, 0.8} is used to determine the portion of broad-sense heritability contributed by additive effects. Each column corresponds to settings where the epistatic effects across traits are independent (*v_α_* = 0) or highly correlated (*v_α_* = 0.8). For comparison, the “trait #1” and “trait #2” dotted lines correspond to the univariate MAPIT model, the “covariance” solid purple line corresponds to power contributed by the covariance test, and the “combined” pink line shows power from the overall association identified by mvMAPIT in the multivariate approach. Note that the upper limit of the x-axis (i.e., false positive rate) has been truncated at 0.05. All results are based on 100 simulated replicates.

**Figure S14.**
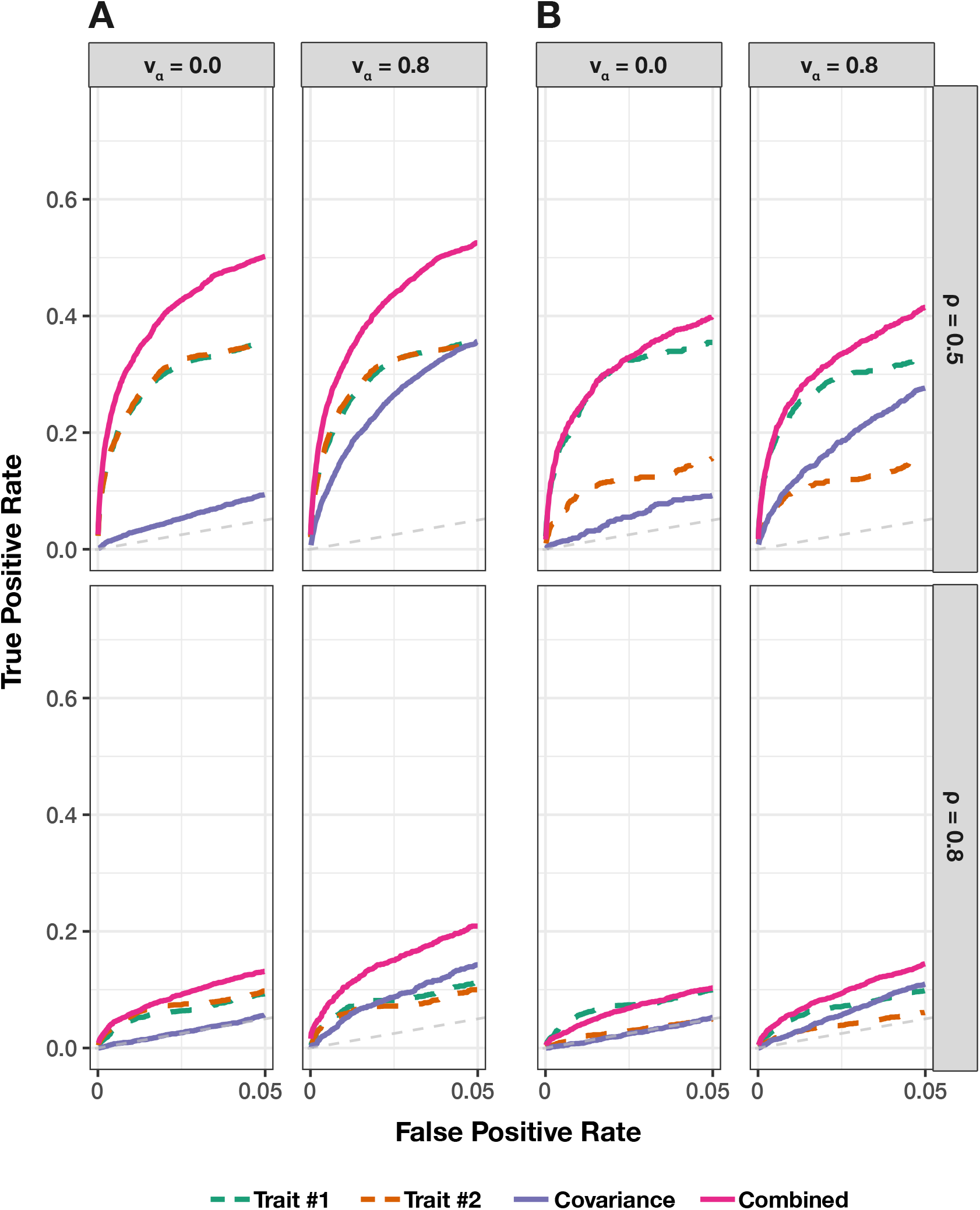
Receiver operating characteristic (ROC) curves comparing the ability of mvMAPIT with Fisher’s method to the univariate MAPIT model in detecting group #1 (10) and group #2 (20) epistatic variants across complex traits. In panel **(A)** both traits have broad-sense heritability *H*^2^ = 0.6; while in panel **(B)** one of traits has broad-sense heritability *H*^2^ = 0.6 and the other has heritability *H*^2^ = 0.3. Across the rows, the parameter p = {0.5, 0.8} is used to determine the portion of broad-sense heritability contributed by additive effects. Each column corresponds to settings where the epistatic effects across traits are independent (*v_α_* = 0) or highly correlated (*v_α_* = 0.8). For comparison, the “trait #1” and “trait #2” dotted lines correspond to the univariate MAPIT model, the “covariance” solid purple line corresponds to power contributed by the covariance test, and the “combined” pink line shows power from the overall association identified by mvMAPIT in the multivariate approach. Note that the upper limit of the x-axis (i.e., false positive rate) has been truncated at 0.05. All results are based on 100 simulated replicates.

**Figure S15.**
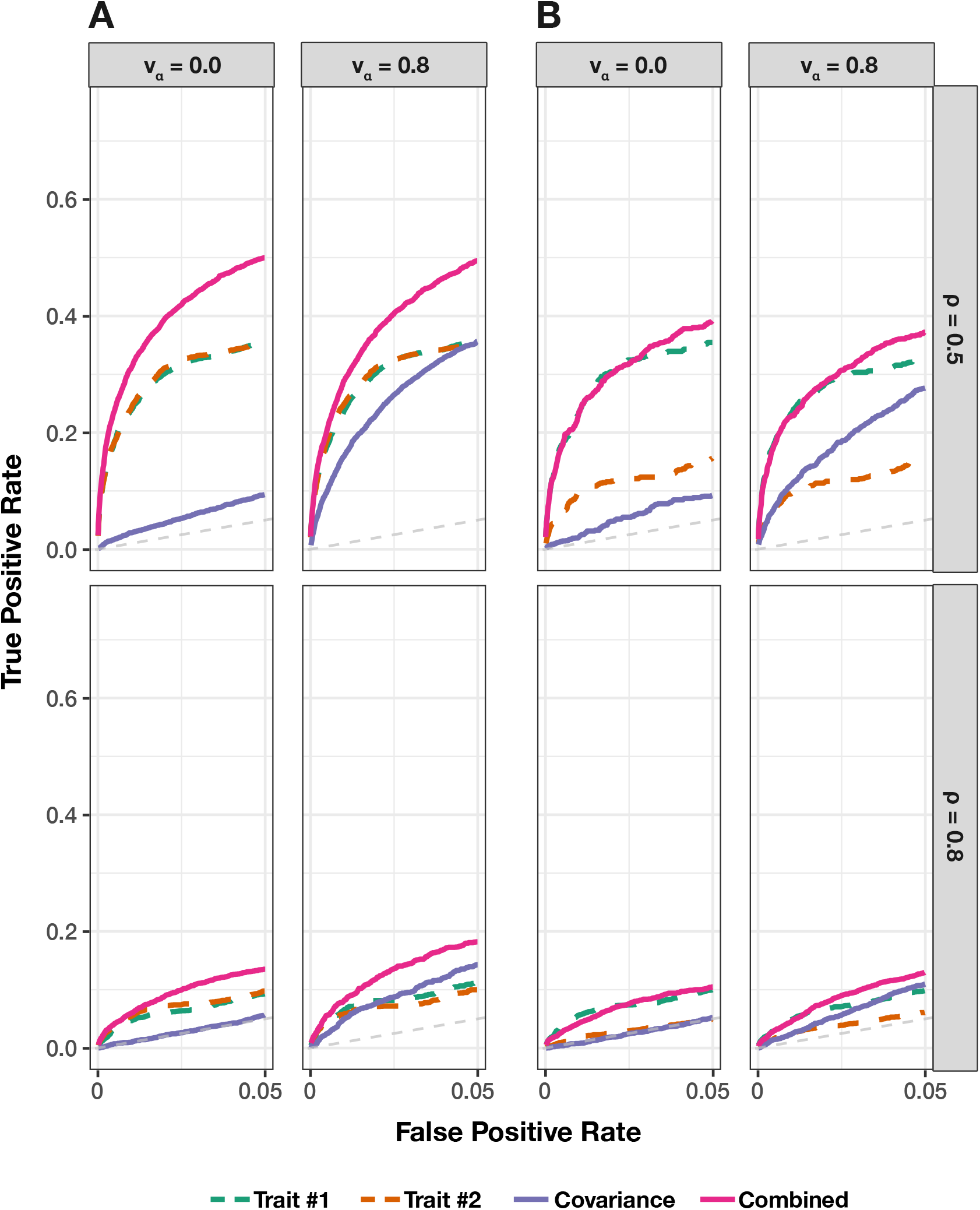
Receiver operating characteristic (ROC) curves comparing the ability of mvMAPIT using the harmonic mean to the univariate MAPIT model in detecting group #1 (10) and group #2 (20) epistatic variants across complex traits. In panel **(A)** both traits have broad-sense heritability *H*^2^ = 0.6; while in panel **(B)** one of traits has broad-sense heritability *H*^2^ = 0.6 and the other has heritability *H*^2^ = 0.3. Across the rows, the parameter *ρ* = {0.5, 0.8} is used to determine the portion of broad-sense heritability contributed by additive effects. Each column corresponds to settings where the epistatic effects across traits are independent (*v_α_* = 0) or highly correlated (*v_α_* = 0.8). For comparison, the “trait #1” and “trait #2” dotted lines correspond to the univariate MAPIT model, the “covariance” solid purple line corresponds to power contributed by the covariance test, and the “combined” pink line shows power from the overall association identified by mvMAPIT in the multivariate approach. Note that the upper limit of the x-axis (i.e., false positive rate) has been truncated at 0.05. All results are based on 100 simulated replicates.

**Figure S16.**
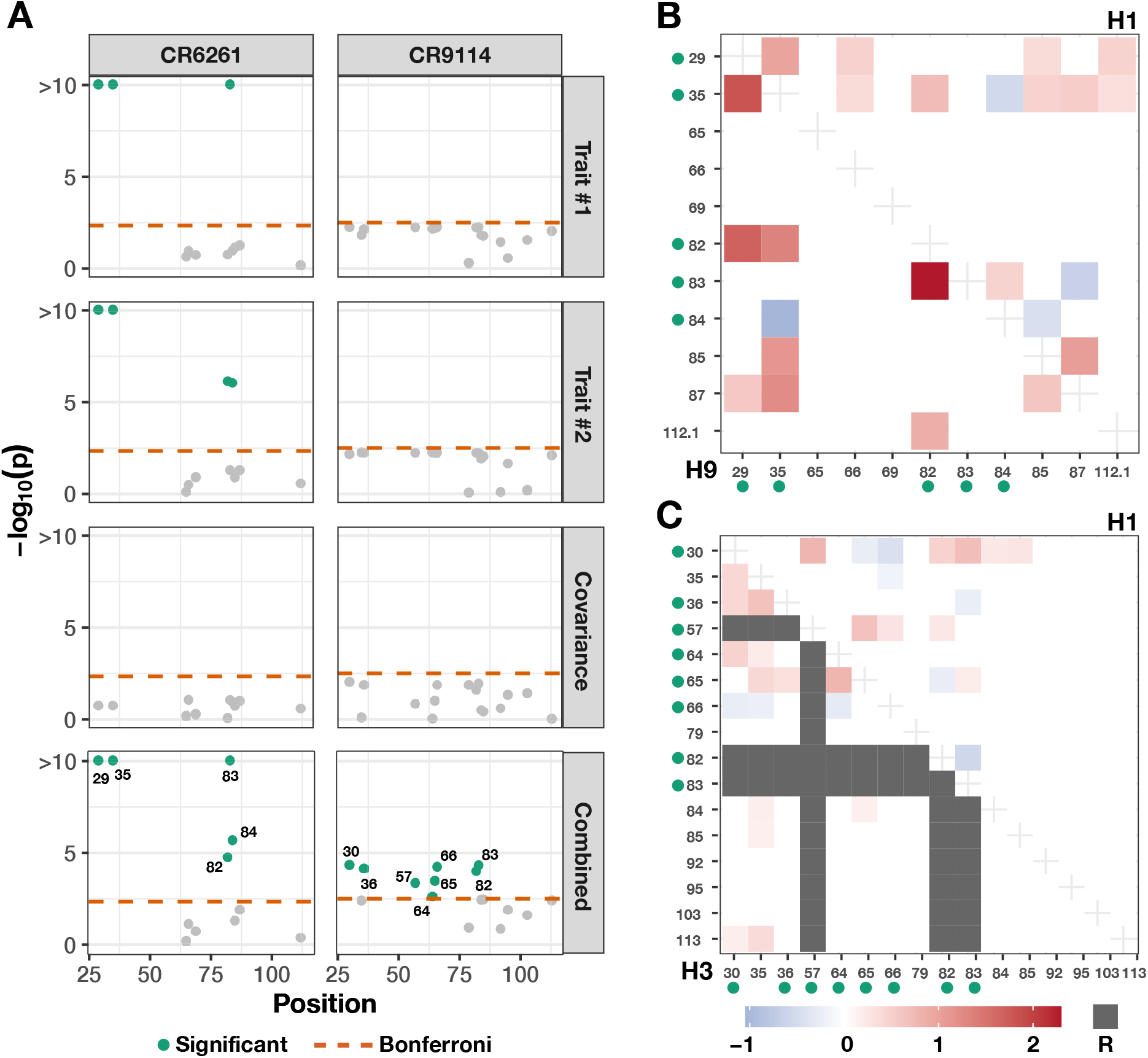
Applying mvMAPIT with the harmonic mean to broadly neutralizing antibodies recovers heavy-chain mutations known to be involved in epistatic interactions that affect binding against two influenza strains. These results are based on protein sequence data from Phillips et al. ^88^ who generated a nearly combinatorially complete library for two broadly neutralizing anti-influenza antibodies (bnAbs), CR6261 and CR9114. For each antibody, we assess binding affinity to different influenza strains. For CR6261, traits #1 and #2 are binding measurements to the antigens *H*_1_ and *H*_9_; while, for CR9114, we assess the same measurement for *H*_1_ and *H*_3_. Panel **(A)** shows Manhattan plots for the different sets of *P*-values computed during the mvMAPIT analysis. The red horizontal lines indicate a chain-wide Bonferroni corrected significance threshold (P = 4.55 × 10^−3^ for CR6261 and *P* = 3.13 × 10^−3^ for CR9114, respectively). The green colored dots are positions that have significant marginal epistatic effects after multiple correction. Panels **(B)** and **(C)** reproduce exhaustive search results originally reported by Phillips et al.^88^. The green dots next to the mutation labels on the axes are the residues that are significant in the multivariate MAPIT analysis and correspond to panel **(A)**. The shaded regions in panel **(B)** are the regression coefficients for pairwise interactions between positions when assessing binding of CR6261with *H*_1_ (upper right triangle) and *H*_9_ (lower left triangle). Similarly, panel **(C)** shows the same information when assessing binding of CR9114 with *H*_1_ (upper right triangle) and *H*_3_ (lower left triangle). Required mutations (indicated by R) are plotted in gray and left out of the analysis^88^.

**Figure S17.**
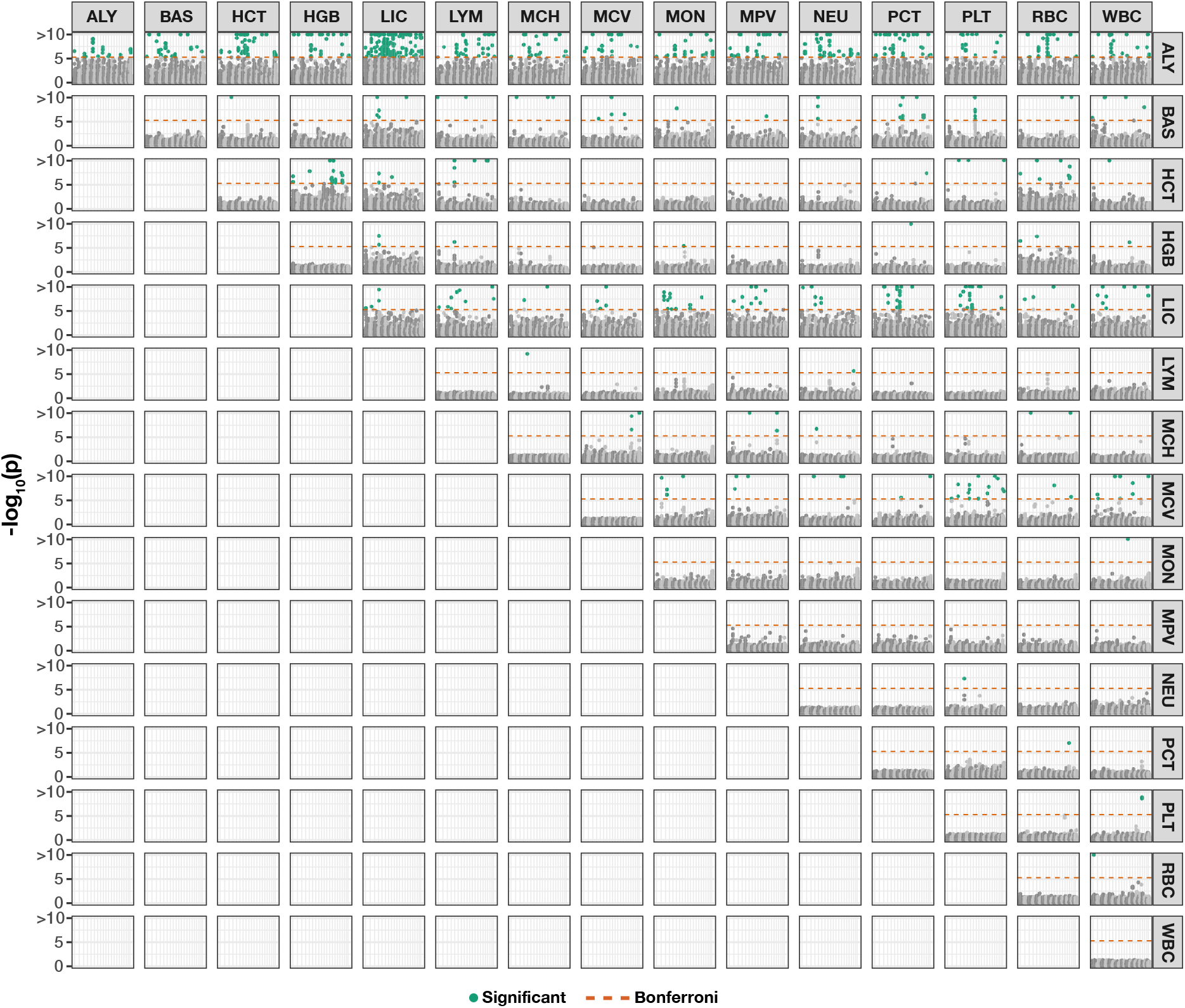
Manhattan plot of genome-wide interaction study for all trait pairs in the heterogenous stock of mice dataset from the Wellcome Trust Centre for Human Genetics^89–91^ using mvMAPIT with Fisher’s method. The columns correspond to trait #1 in the analysis while the rows denote trait #2. Results on the diagonal correspond to results from running a univariate MAPIT model. The results on the off-diagonals show the combined *P*-values from mvMAPIT. The red horizontal lines indicate a genome-wide Bonferroni corrected significance threshold (*P* = 4.83 × 10^−6^). The green colored dots are SNPs that have significant marginal epistatic effects after multiple correction. Full names for the abbreviations of each trait can be found in the main text (Materials and Methods).

**Figure S18.**
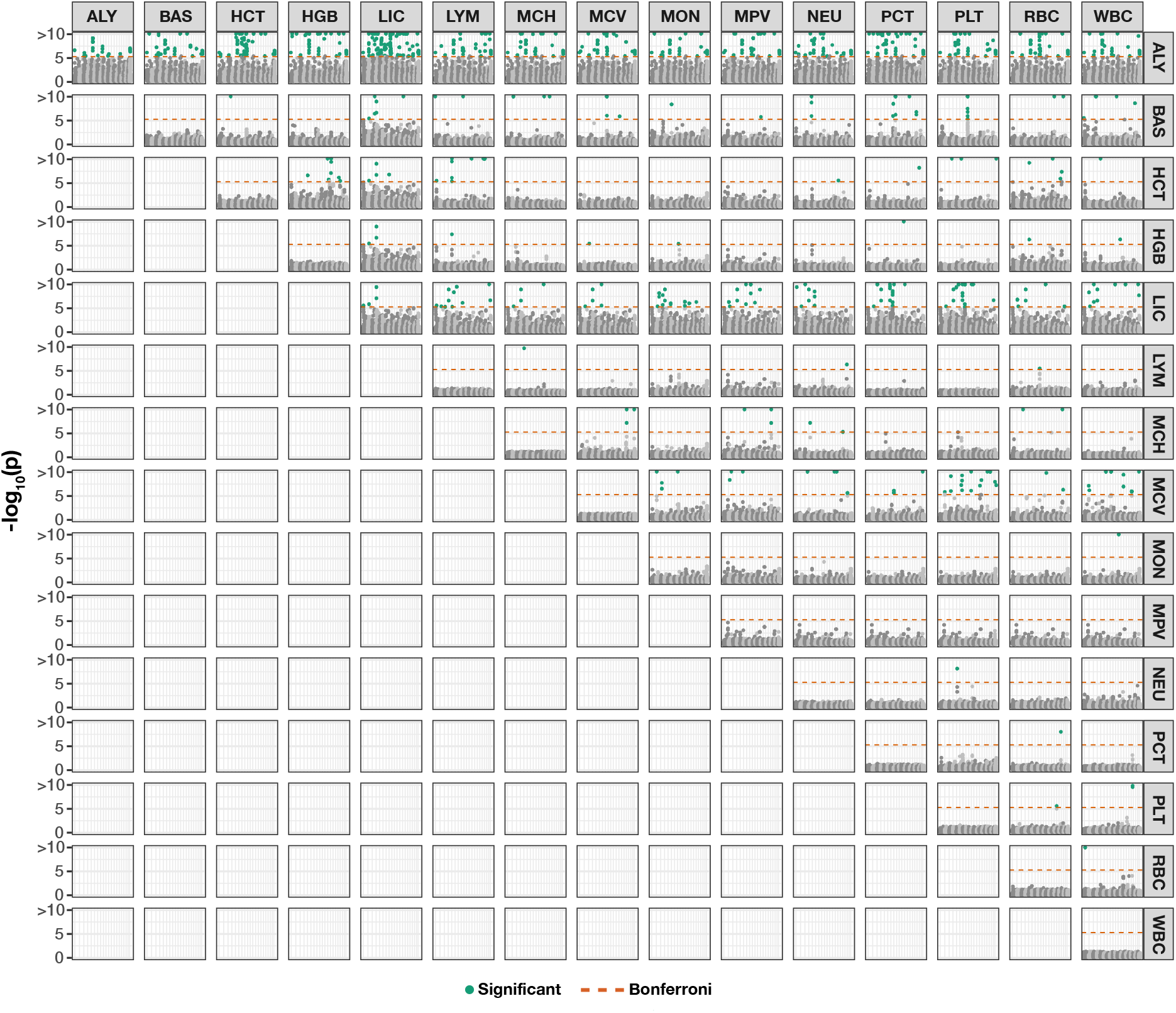
Manhattan plot of genome-wide interaction study for all trait pairs in the heterogenous stock of mice dataset from the Wellcome Trust Centre for Human Genetics^89–91^ using mvMAPIT with the harmonic mean. The columns correspond to trait #1 in the analysis while the rows denote trait #2. Results on the diagonal correspond to results from running a univariate MAPIT model. The results on the off-diagonals show the combined *P*-values from mvMAPIT. The red horizontal lines indicate a genome-wide Bonferroni corrected significance threshold (*P* = 4.83 × 10^−6^). The green colored dots are SNPs that have significant marginal epistatic effects after multiple correction. Full names for the abbreviations of each trait can be found in the main text (Materials and Methods).

**Figure S19.**
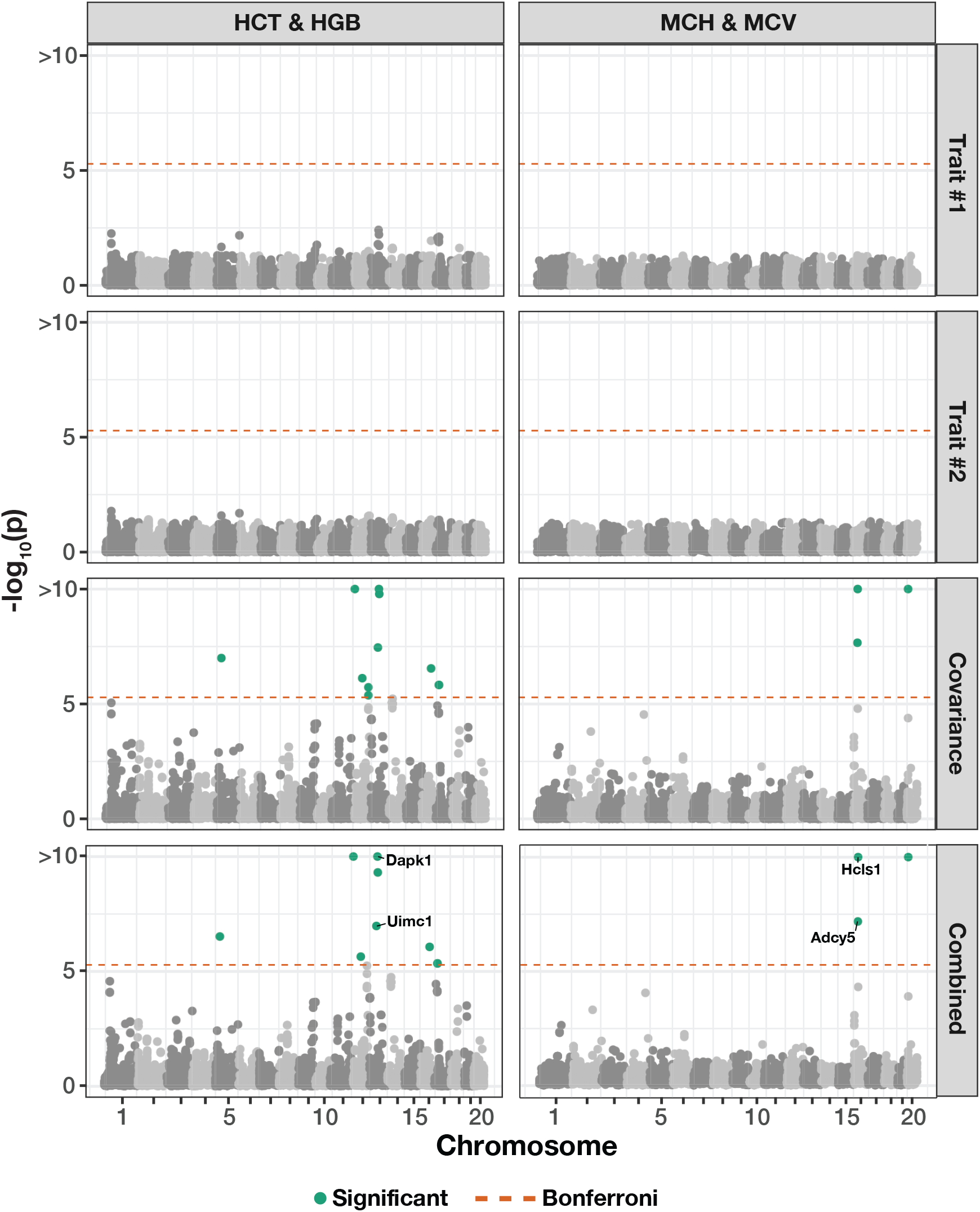
Manhattan plot of genome-wide interaction study for two pairs of hematology traits in the heterogenous stock of mice dataset from the Wellcome Trust Centre for Human Genetics^89–91^ using mvMAPIT with the harmonic mean. The trait pairs in this figure include hematocrit (HCT) and hemoglobin (HGB) in the left column and mean corpuscular hemoglobin (MCH) and mean corpuscular volume (MCV) in the right column. Here, we depict the *P*-values computed during each step of the mvMAPIT modeling pipeline. The red horizontal lines indicate a genome-wide Bonferroni corrected significance threshold (*P* = 4.83 × 10^−6^). The green colored dots are SNPs that have significant marginal epistatic effects after multiple test correction. Significant SNPs were mapped to the closest neighboring genes using the Mouse Genome Informatics database (http://www.informatics.jax.org)^106^.

**Figure S20.**
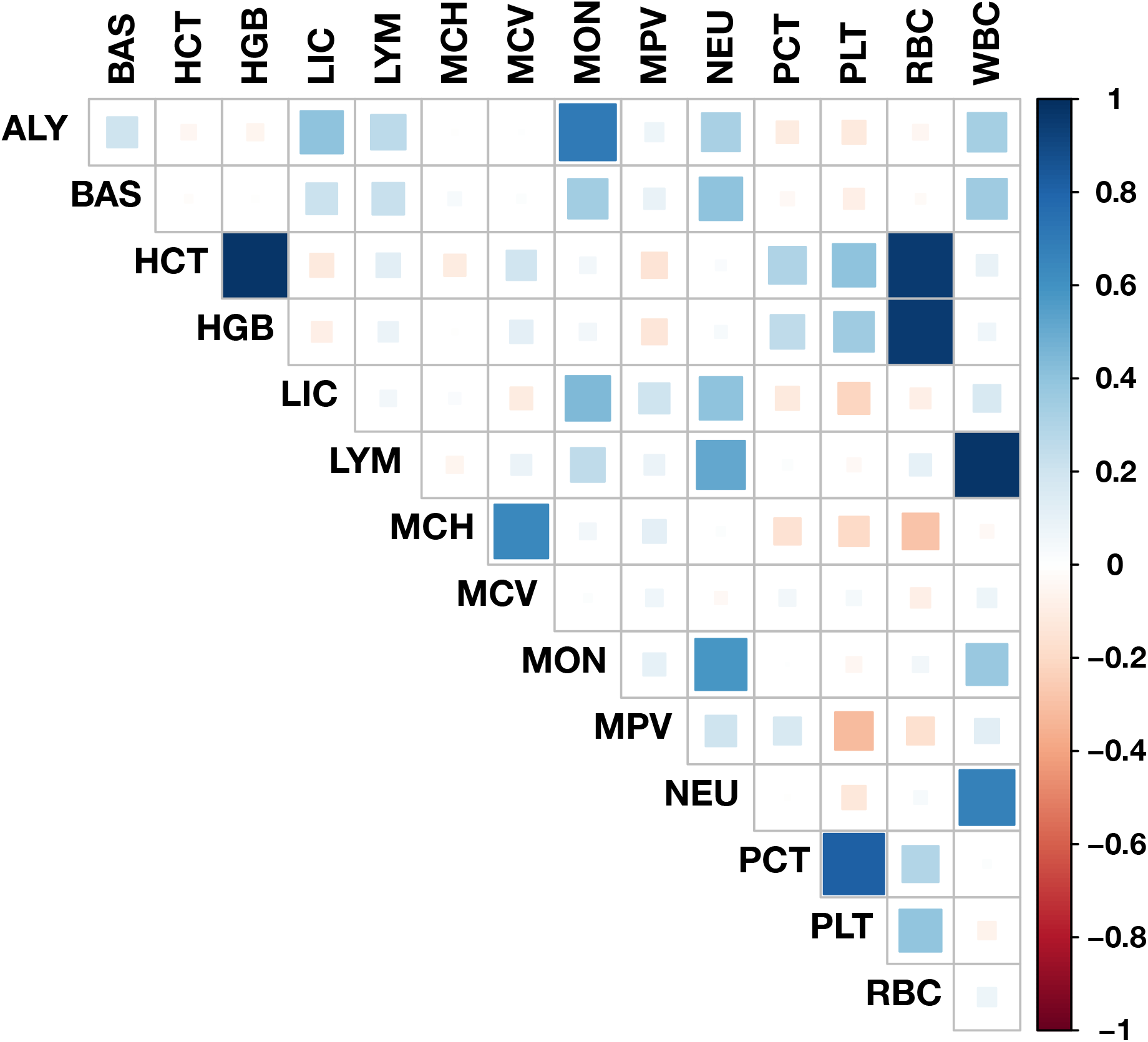
Empirical correlations for all trait pairs in the heterogenous stock of mice dataset from the Wellcome Trust Centre for Human Genetics^89–91^. Full names for the abbreviations of each trait can be found in the main text (Materials and Methods).

## Supplementary Tables

**Table S1.** The mvMAPIT framework using Fisher’s method preserves type I error rates under the null model when traits are generated by only additive effects (sample size *N* = 1,000 individuals). In these simulations, quantitative traits are simulated to have narrow-sense heritability *h*^2^ = 0.6 with an architecture made up of only additive genetic variation. Each row corresponds to a setting where the additive genetic effects for a causal SNP have different correlation structures across traits. In these simulations, we consider scenarios where we have traits with independent additive effects (*v_β_* = 0), traits with highly correlated additive effects (*v_β_* = 0.8), and traits with perfectly correlated additive effects (*v_β_* = 1). We assess the calibration of the *P*-values that are produced by mvMAPIT during each of the three key steps in its combinatorial hypothesis testing procedure (see Materials and Methods). We show type I error rates resulting from *P*-values taken from the “univariate” test on each trait independently, the “covariance” *P*-values which corresponds to assessing the pairwise interactions affecting both traits, and the final “combined” *P*-value. Results are summarized over 100 simulated replicates. Values in the parentheses are the standard deviations across replicates.

| | Add. Effect Corr. | $P = 0.05$ | $P = 0.01$ | $P = 0.001$ |
| --- | --- | --- | --- | --- |
| Univariate | $v_\beta = 0.0$ | 0.030 ($1 \times 10^{-2}$) | 0.007 ($3 \times 10^{-3}$) | 0.0007 ($6 \times 10^{-4}$) |
| | $v_\beta = 0.8$ | 0.030 ($1 \times 10^{-2}$) | 0.007 ($2 \times 10^{-3}$) | 0.0007 ($6 \times 10^{-4}$) |
| | $v_\beta = 1.0$ | 0.030 ($1 \times 10^{-2}$) | 0.007 ($2 \times 10^{-3}$) | 0.0007 ($6 \times 10^{-4}$) |
| Covariance | $v_\beta = 0.0$ | 0.040 ($1 \times 10^{-2}$) | 0.005 ($2 \times 10^{-3}$) | 0.0002 ($4 \times 10^{-4}$) |
| | $v_\beta = 0.8$ | 0.040 ($1 \times 10^{-2}$) | 0.005 ($2 \times 10^{-3}$) | 0.0002 ($3 \times 10^{-4}$) |
| | $v_\beta = 1.0$ | 0.040 ($1 \times 10^{-2}$) | 0.005 ($2 \times 10^{-3}$) | 0.0003 ($3 \times 10^{-4}$) |
| Combined | $v_\beta = 0.0$ | 0.030 ($1 \times 10^{-2}$) | 0.006 ($2 \times 10^{-3}$) | 0.0005 ($7 \times 10^{-4}$) |
| | $v_\beta = 0.8$ | 0.040 ($1 \times 10^{-2}$) | 0.006 ($2 \times 10^{-3}$) | 0.0005 ($4 \times 10^{-4}$) |
| | $v_\beta = 1.0$ | 0.040 ($1 \times 10^{-2}$) | 0.006 ($2 \times 10^{-3}$) | 0.0006 ($6 \times 10^{-4}$) |

**Table S2.** The mvMAPIT framework using Fisher’s method preserves type I error rates under the null model when traits are generated by only additive effects (sample size *N* = 1,750 individuals). In these simulations, quantitative traits are simulated to have narrow-sense heritability *h*^2^ = 0.6 with an architecture made up of only additive genetic variation. Each row corresponds to a setting where the additive genetic effects for a causal SNP have different correlation structures across traits. In these simulations, we consider scenarios where we have traits with independent additive effects (*v_β_* = 0), traits with highly correlated additive effects (*v_β_* = 0.8), and traits with perfectly correlated additive effects (*v_β_* = 1). We assess the calibration of the *P*-values that are produced by mvMAPIT during each of the three key steps in its combinatorial hypothesis testing procedure (see Materials and Methods). We show type I error rates resulting from *P*-values taken from the “univariate” test on each trait independently, the “covariance” *P*-values which corresponds to assessing the pairwise interactions affecting both traits, and the final “combined” *P*-value. Results are summarized over 100 simulated replicates. Values in the parentheses are the standard deviations across replicates.

| | Add. Effect Corr. | $P = 0.05$ | $P = 0.01$ | $P = 0.001$ |
| --- | --- | --- | --- | --- |
| Univariate | $v_\beta = 0.0$ | 0.030 ($1 \times 10^{-2}$) | 0.008 ($2 \times 10^{-3}$) | 0.0007 ($5 \times 10^{-4}$) |
| | $v_\beta = 0.8$ | 0.030 ($1 \times 10^{-2}$) | 0.008 ($2 \times 10^{-3}$) | 0.0009 ($7 \times 10^{-4}$) |
| | $v_\beta = 1.0$ | 0.030 ($1 \times 10^{-2}$) | 0.008 ($3 \times 10^{-3}$) | 0.0009 ($9 \times 10^{-4}$) |
| Covariance | $v_\beta = 0.0$ | 0.040 ($1 \times 10^{-2}$) | 0.006 ($2 \times 10^{-3}$) | 0.0003 ($4 \times 10^{-4}$) |
| | $v_\beta = 0.8$ | 0.040 ($1 \times 10^{-2}$) | 0.006 ($2 \times 10^{-3}$) | 0.0002 ($3 \times 10^{-4}$) |
| | $v_\beta = 1.0$ | 0.040 ($1 \times 10^{-2}$) | 0.006 ($2 \times 10^{-3}$) | 0.0002 ($3 \times 10^{-4}$) |
| Combined | $v_\beta = 0.0$ | 0.040 ($1 \times 10^{-2}$) | 0.007 ($2 \times 10^{-3}$) | 0.0006 ($5 \times 10^{-4}$) |
| | $v_\beta = 0.8$ | 0.040 ($1 \times 10^{-2}$) | 0.007 ($2 \times 10^{-3}$) | 0.0007 ($8 \times 10^{-4}$) |
| | $v_\beta = 1.0$ | 0.040 ($1 \times 10^{-2}$) | 0.007 ($2 \times 10^{-3}$) | 0.0006 ($6 \times 10^{-4}$) |

**Table S3.** The mvMAPIT framework using the harmonic mean preserves type I error rates under the null model when traits are generated by only additive effects (sample size *N* = 1,000 individuals). In these simulations, quantitative traits are simulated to have narrowsense heritability *h*^2^ = 0.6 with an architecture made up of only additive genetic variation. Each row corresponds to a setting where the additive genetic effects for a causal SNP have different correlation structures across traits. In these simulations, we consider scenarios where we have traits with independent additive effects (*v_β_* = 0), traits with highly correlated additive effects (*v_β_* = 0.8), and traits with perfectly correlated additive effects (*v_β_* = 1). We assess the calibration of the *P*-values that are produced by mvMAPIT during each of the three key steps in its combinatorial hypothesis testing procedure (see Materials and Methods). We show type I error rates resulting from *P*-values taken from the “univariate” test on each trait independently, the “covariance” *P*-values which corresponds to assessing the pairwise interactions affecting both traits, and the final “combined” *P*-value. Results are summarized over 100 simulated replicates. Values in the parentheses are the standard deviations across replicates.

| | Add. Effect Corr. | $P = 0.05$ | $P = 0.01$ | $P = 0.001$ |
| --- | --- | --- | --- | --- |
| Univariate | $v_\beta = 0.0$ | 0.030 ($1 \times 10^{-2}$) | 0.007 ($3 \times 10^{-3}$) | 0.0007 ($6 \times 10^{-4}$) |
| | $v_\beta = 0.8$ | 0.030 ($1 \times 10^{-2}$) | 0.007 ($2 \times 10^{-3}$) | 0.0007 ($6 \times 10^{-4}$) |
| | $v_\beta = 1.0$ | 0.030 ($1 \times 10^{-2}$) | 0.007 ($2 \times 10^{-3}$) | 0.0007 ($6 \times 10^{-4}$) |
| Covariance | $v_\beta = 0.0$ | 0.040 ($1 \times 10^{-2}$) | 0.005 ($2 \times 10^{-3}$) | 0.0002 ($4 \times 10^{-4}$) |
| | $v_\beta = 0.8$ | 0.040 ($1 \times 10^{-2}$) | 0.005 ($2 \times 10^{-3}$) | 0.0002 ($3 \times 10^{-4}$) |
| | $v_\beta = 1.0$ | 0.040 ($1 \times 10^{-2}$) | 0.005 ($2 \times 10^{-3}$) | 0.0003 ($3 \times 10^{-4}$) |
| Combined | $v_\beta = 0.0$ | 0.040 ($1 \times 10^{-2}$) | 0.006 ($2 \times 10^{-3}$) | 0.0005 ($5 \times 10^{-4}$) |
| | $v_\beta = 0.8$ | 0.040 ($1 \times 10^{-2}$) | 0.006 ($2 \times 10^{-3}$) | 0.0004 ($4 \times 10^{-4}$) |
| | $v_\beta = 1.0$ | 0.040 ($1 \times 10^{-2}$) | 0.006 ($2 \times 10^{-3}$) | 0.0005 ($5 \times 10^{-4}$) |

**Table S4.** The mvMAPIT framework using the harmonic mean preserves type I error rates under the null model when traits are generated by only additive effects (sample size *N* = 1,750 individuals). In these simulations, quantitative traits are simulated to have narrowsense heritability *h*^2^ = 0.6 with an architecture made up of only additive genetic variation. Each row corresponds to a setting where the additive genetic effects for a causal SNP have different correlation structures across traits. In these simulations, we consider scenarios where we have traits with independent additive effects (*v_β_* = 0), traits with highly correlated additive effects (*v_β_* = 0.8), and traits with perfectly correlated additive effects (*v_β_* = 1). We assess the calibration of the *P*-values that are produced by mvMAPIT during each of the three key steps in its combinatorial hypothesis testing procedure (see Materials and Methods). We show type I error rates resulting from *P*-values taken from the “univariate” test on each trait independently, the “covariance” *P*-values which corresponds to assessing the pairwise interactions affecting both traits, and the final “combined” *P*-value. Results are summarized over 100 simulated replicates. Values in the parentheses are the standard deviations across replicates.

| | Add. Effect Corr. | $P = 0.05$ | $P = 0.01$ | $P = 0.001$ |
| --- | --- | --- | --- | --- |
| Univariate | $v_\beta = 0.0$ | 0.030 ($1 \times 10^{-2}$) | 0.008 ($2 \times 10^{-3}$) | 0.0007 ($5 \times 10^{-4}$) |
| | $v_\beta = 0.8$ | 0.030 ($1 \times 10^{-2}$) | 0.008 ($2 \times 10^{-3}$) | 0.0009 ($7 \times 10^{-4}$) |
| | $v_\beta = 1.0$ | 0.030 ($1 \times 10^{-2}$) | 0.008 ($3 \times 10^{-3}$) | 0.0009 ($9 \times 10^{-4}$) |
| Covariance | $v_\beta = 0.0$ | 0.040 ($1 \times 10^{-2}$) | 0.006 ($2 \times 10^{-3}$) | 0.0003 ($4 \times 10^{-4}$) |
| | $v_\beta = 0.8$ | 0.040 ($1 \times 10^{-2}$) | 0.006 ($2 \times 10^{-3}$) | 0.0002 ($3 \times 10^{-4}$) |
| | $v_\beta = 1.0$ | 0.040 ($1 \times 10^{-2}$) | 0.006 ($2 \times 10^{-3}$) | 0.0002 ($3 \times 10^{-4}$) |
| Combined | $v_\beta = 0.0$ | 0.040 ($1 \times 10^{-2}$) | 0.008 ($2 \times 10^{-3}$) | 0.0006 ($5 \times 10^{-4}$) |
| | $v_\beta = 0.8$ | 0.040 ($1 \times 10^{-2}$) | 0.008 ($2 \times 10^{-3}$) | 0.0007 ($8 \times 10^{-4}$) |
| | $v_\beta = 1.0$ | 0.040 ($1 \times 10^{-2}$) | 0.007 ($2 \times 10^{-3}$) | 0.0005 ($5 \times 10^{-4}$) |

**Table S5.** The mvMAPIT framework using the harmonic mean preserves type I error rates under the null model when traits are generated by only additive effects (sample size *N* = 2,500 individuals). In these simulations, quantitative traits are simulated to have narrowsense heritability *h*^2^ = 0.6 with an architecture made up of only additive genetic variation. Each row corresponds to a setting where the additive genetic effects for a causal SNP have different correlation structures across traits. In these simulations, we consider scenarios where we have traits with independent additive effects (*v_β_* = 0), traits with highly correlated additive effects (*v_β_* = 0.8), and traits with perfectly correlated additive effects (*v_β_* = 1). We assess the calibration of the *P*-values that are produced by mvMAPIT during each of the three key steps in its combinatorial hypothesis testing procedure (see Materials and Methods). We show type I error rates resulting from *P*-values taken from the “univariate” test on each trait independently, the “covariance” *P*-values which corresponds to assessing the pairwise interactions affecting both traits, and the final “combined” *P*-value. Results are summarized over 100 simulated replicates. Values in the parentheses are the standard deviations across replicates.

| | Add. Effect Corr. | $P = 0.05$ | $P = 0.01$ | $P = 0.001$ |
| --- | --- | --- | --- | --- |
| Univariate | $v_\beta = 0.0$ | 0.030 ($1 \times 10^{-2}$) | 0.009 ($2 \times 10^{-3}$) | 0.0010 ($9 \times 10^{-4}$) |
| | $v_\beta = 0.8$ | 0.030 ($1 \times 10^{-2}$) | 0.009 ($2 \times 10^{-3}$) | 0.0009 ($7 \times 10^{-4}$) |
| | $v_\beta = 1.0$ | 0.030 ($1 \times 10^{-2}$) | 0.009 ($3 \times 10^{-3}$) | 0.0009 ($7 \times 10^{-4}$) |
| Covariance | $v_\beta = 0.0$ | 0.040 ($1 \times 10^{-2}$) | 0.006 ($2 \times 10^{-3}$) | 0.0003 ($4 \times 10^{-4}$) |
| | $v_\beta = 0.8$ | 0.040 ($1 \times 10^{-2}$) | 0.007 ($2 \times 10^{-3}$) | 0.0004 ($5 \times 10^{-4}$) |
| | $v_\beta = 1.0$ | 0.040 ($1 \times 10^{-2}$) | 0.006 ($2 \times 10^{-3}$) | 0.0003 ($4 \times 10^{-4}$) |
| Combined | $v_\beta = 0.0$ | 0.040 ($1 \times 10^{-2}$) | 0.008 ($2 \times 10^{-3}$) | 0.0007 ($6 \times 10^{-4}$) |
| | $v_\beta = 0.8$ | 0.040 ($1 \times 10^{-2}$) | 0.008 ($2 \times 10^{-3}$) | 0.0007 ($6 \times 10^{-4}$) |
| | $v_\beta = 1.0$ | 0.040 ($1 \times 10^{-2}$) | 0.008 ($2 \times 10^{-3}$) | 0.0005 ($6 \times 10^{-4}$) |

**Table S6. Complete summary of the marginal epistatic results after applying the mvMAPIT framework to protein sequence data from a nearly combinatorially complete library of two broadly neutralizing anti-influenza antibodies.** Here, data is from Phillips et al.^88^ who generated a nearly combinatorially complete library for two broadly neutralizing anti-influenza antibodies (bnAbs), CR6261 and CR9114. In the first column, we list the antibody being analyzed. In the second column, we give their corresponding residue. In the third and fourth columns, we list all the pairwise antigen combinations done in the analysis. In the remaining columns, we give the results stemming from univariate analyses on antigens #1 and #2, respectively, the covariance (cov) test, and the overall *P*-values derived by mvMAPIT using both Fisher’s method and the harmonic mean. Tutorials for how to take these results and recreate the Manhattan plots shown in Figures 5 and S16 can be found in the mvMAPIT GitHub repository (see Materials and Methods). (XLSX)

**Table S7. Complete summary of the marginal epistatic results after applying the mvMAPIT framework to 15 hematology traits in the heterogenous stock of mice dataset from the Wellcome Trust Centre for Human Genetics^89–91^.** In the first column, we list the ID of each SNP. In the second and third columns, we give their corresponding chromosome and basepair according to the mouse assembly NCBI build 34 (accessed from Shifman et al.^135^). In the fourth and fifth columns, we list all the pairwise trait combinations done in the analysis. In the remaining columns, we give the results stemming from univariate analyses on traits #1 and #2, respectively, the covariance (cov) test, and the overall *P*-values derived by mvMAPIT using both Fisher’s method and the harmonic mean. Tutorials for how to take these results and recreate the Manhattan plots shown in Figures S17 and S18 can be found in the mvMAPIT GitHub repository (see Materials and Methods). (XLSX)

## References

[1] Stephan Ripke, Benjamin M. Neale, Aiden Corvin, James T. R. Walters, Kai-How Farh, Peter A. Holmans, Phil Lee, Brendan Bulik-Sullivan, David A. Collier, Hailiang Huang, Tune H. Pers, Ingrid Agartz, Esben Agerbo, Margot Albus, Madeline Alexander, Farooq Amin, Silviu A. Bacanu, Martin Begemann, Richard A. Belliveau Jr, Judit Bene, Sarah E. Bergen, Elizabeth Bevilacqua, Tim B. Bigdeli, Donald W. Black, Richard Bruggeman, Nancy G. Buccola, Randy L. Buckner, William Byerley, Wiepke Cahn, Guiqing Cai, Dominique Campion, Rita M. Cantor, Vaughan J. Carr, Noa Carrera, Stanley V. Catts, Kimberly D. Chambert, Raymond C. K. Chan, Ronald Y. L. Chen, Eric Y. H. Chen, Wei Cheng, Eric F. C. Cheung, Siow Ann Chong, C. Robert Cloninger, David Cohen, Nadine Cohen, Paul Cormican, Nick Craddock, James J. Crowley, David Curtis, Michael Davidson, Kenneth L. Davis, Franziska Degenhardt, Jurgen Del Favero, Ditte Demontis, Dimitris Dikeos, Timothy Dinan, Srdjan Djurovic, Gary Donohoe, Elodie Drapeau, Jubao Duan, Frank Dudbridge, Naser Durmishi, Peter Eichhammer, Johan Eriksson, Valentina Escott-Price, Laurent Essioux, Ayman H. Fanous, Martilias S. Farrell, Josef Frank, Lude Franke, Robert Freedman, Nelson B. Freimer, Marion Friedl, Joseph I. Friedman, Menachem Fromer, Giulio Genovese, Lyudmila Georgieva, Ina Giegling, Paola Giusti-Rodríguez, Stephanie Godard, Jacqueline I. Goldstein, Vera Golimbet, Srihari Gopal, Jacob Gratten, Lieuwe de Haan, Christian Hammer, Marian L. Hamshere, Mark Hansen, Thomas Hansen, Vahram Haroutunian, Annette M. Hartmann, Frans A. Henskens, Stefan Herms, Joel N. Hirschhorn, Per Hoffmann, Andrea Hofman, Mads V. Hollegaard, David M. Hougaard, Masashi Ikeda, Inge Joa, Antonio Julià, René S. Kahn, Luba Kalaydjieva, Sena Karachanak-Yankova, Juha Karjalainen, David Kavanagh, Matthew C. Keller, James L. Kennedy, Andrey Khrunin, Yunjung Kim, Janis Klovins, James A. Knowles, Bettina Konte, Vaidutis Kucinskas, Zita Ausrele Kucinskiene, Hana Kuzelova-Ptackova, Anna K. Kähler, Claudine Laurent, Jimmy Lee Chee Keong, S. Hong Lee, Sophie E. Legge, Bernard Lerer, Miaoxin Li, Tao Li, Kung-Yee Liang, Jeffrey Lieberman, Svetlana Limborska, Carmel M. Loughland, Jan Lubinski, Jouko Lönnqvist, Milan Macek Jr, Patrik K. E. Magnusson, Brion S. Maher, Wolfgang Maier, Jacques Mallet, Sara Marsal, Manuel Mattheisen, Morten Mattingsdal, Robert W. McCarley, Colm McDonald, Andrew M. McIntosh, Sandra Meier, Carin J. Meijer, Bela Melegh, Ingrid Melle, Raquelle I. Mesholam-Gately, Andres Metspalu, Patricia T. Michie, Lili Milani, Vihra Milanova, Younes Mokrab, Derek W. Morris, Ole Mors, Kieran C. Murphy, Robin M. Murray, Inez Myin-Germeys, Bertram Müller-Myhsok, Mari Nelis, Igor Nenadic, Deborah A. Nertney, Gerald Nestadt, Kristin K. Nicodemus, Liene Nikitina-Zake, Laura Nisenbaum, Annelie Nordin, Eadbhard O’Callaghan, Colm O’Dushlaine, F. Anthony O’Neill, Sang-Yun Oh, Ann Olincy, Line Olsen, Jim Van Os, Christos Pantelis, George N. Papadimitriou, Sergi Papiol, Elena Parkhomenko, Michele T. Pato, Tiina Paunio, Milica Pejovic-Milovancevic, Diana O. Perkins, Olli Pietiläinen, Jonathan Pimm, Andrew J. Pocklington, John Powell, Alkes Price, Ann E. Pulver, Shaun M. Purcell, Digby Quested, Henrik B. Rasmussen, Abraham Reichenberg, Mark A. Reimers, Alexander L. Richards, Joshua L. Roffman, Panos Roussos, Douglas M. Ruderfer, Veikko Salomaa, Alan R. Sanders, Ulrich Schall, Christian R. Schubert, Thomas G. Schulze, Sibylle G. Schwab, Edward M. Scolnick, Rodney J. Scott, Larry J. Seidman, Jianxin Shi, Engilbert Sigurdsson, Teimuraz Silagadze, Jeremy M. Silverman, Kang Sim, Petr Slominsky, Jordan W. Smoller, Hon-Cheong So, ChrisC. A. Spencer, Eli A. Stahl, Hreinn Stefansson, Stacy Steinberg, Elisabeth Stogmann, Richard E. Straub, Eric Strengman, Jana Strohmaier, T. Scott Stroup, Mythily Subramaniam, Jaana Suvisaari, Dragan M. Svrakic, Jin P. Szatkiewicz, Erik Söderman, Srinivas Thirumalai, Draga Toncheva, Sarah Tosato, Juha Veijola, John Waddington, Dermot Walsh, Dai Wang, Qiang Wang, Bradley T. Webb, Mark Weiser, Dieter B. Wildenauer, Nigel M. Williams, Stephanie Williams, Stephanie H. Witt, Aaron R. Wolen, Emily H. M. Wong, Brandon K. Wormley, Hualin Simon Xi, Clement C. Zai, Xuebin Zheng, Fritz Zimprich, Naomi R. Wray, Kari Stefansson, Peter M. Visscher, Wellcome Trust Case-Control Consortium, Rolf Adolfsson, Ole A. Andreassen, Douglas H. R. Blackwood, Elvira Bramon, Joseph D. Buxbaum, Anders D. Børglum, Sven Cichon, Ariel Darvasi, Enrico Domenici, Hannelore Ehrenreich, Tõnu Esko, Pablo V. Gejman, Michael Gill, Hugh Gurling, Christina M. Hultman, Nakao Iwata, Assen V. Jablensky, Erik G. Jönsson, Kenneth S. Kendler, George Kirov, Jo Knight, Todd Lencz, Douglas F. Levinson, Qingqin S. Li, Jianjun Liu, Anil K. Malhotra, Steven A. McCarroll, Andrew McQuillin, Jennifer L. Moran, Preben B. Mortensen, Bryan J. Mowry, Markus M. Nöthen, Roel A. Ophoff, Michael J. Owen, Aarno Palotie, Carlos N. Pato, Tracey L. Petryshen, Danielle Posthuma, Marcella Rietschel, Brien P. Riley, Dan Rujescu, Pak C. Sham, Pamela Sklar, David St Clair, Daniel R. Weinberger, Jens R. Wendland, Thomas Werge, Schizophrenia Working Group of the Psychiatric Genomics Consortium, and Psychosis Endophenotypes International Consortium. Biological insights from 108 schizophrenia-associated genetic loci. Nature, 511(7510):421–427, 2014. ISSN 1476-4687. doi: 10.1038/nature13595. URL https://www.nature.com/articles/nature13595. Number: 7510 Publisher: Nature Publishing Group.

[2] David Ellinghaus, Luke Jostins, Sarah L. Spain, Adrian Cortes, Jörn Bethune, Buhm Han, Yu Rang Park, Soumya Raychaudhuri, Jennie G. Pouget, Matthias Hübenthal, Trine Folseraas, Yunpeng Wang, Tonu Esko, Andres Metspalu, Harm-Jan Westra, Lude Franke, Tune H. Pers, Rinse K. Weersma, Valerie Collij, Mauro D’Amato, Jonas Halfvarson, Anders Boeck Jensen, Wolfgang Lieb, Franziska Degenhardt, Andreas J. Forstner, Andrea Hofmann, Stefan Schreiber, Ulrich Mrowietz, Brian D. Juran, Konstantinos N. Lazaridis, Søren Brunak, Anders M. Dale, Richard C. Trembath, Stephan Weidinger, Michael Weichenthal, Eva Ellinghaus, James T. Elder, Jonathan N. W. N. Barker, Ole A. Andreassen, Dermot P. McGovern, Tom H. Karlsen, Jeffrey C. Barrett, Miles Parkes, Matthew A. Brown, and Andre Franke. Analysis of five chronic inflammatory diseases identifies 27 new associations and highlights disease-specific patterns at shared loci. Nature Genetics, 48(5): 510–518, 2016. ISSN 1546-1718. doi: 10.1038/ng.3528. URL https://www.nature.com/articles/ng.3528. Number: 5 Publisher: Nature Publishing Group.

[3] Christian Fuchsberger, Jason Flannick, Tanya M. Teslovich, Anubha Mahajan, Vineeta Agarwala, Kyle J. Gaulton, Clement Ma, Pierre Fontanillas, Loukas Moutsianas, Davis J. McCarthy, Manuel A. Rivas, John R. B. Perry, Xueling Sim, Thomas W. Blackwell, Neil R. Robertson, N. William Rayner, Pablo Cingolani, Adam E. Locke, Juan Fernandez Tajes, Heather M. Highland, Josee Dupuis, Peter S. Chines, Cecilia M. Lindgren, Christopher Hartl, Anne U. Jackson, Han Chen, Jeroen R. Huyghe, Martijn van de Bunt, Richard D. Pearson, Ashish Kumar, Martina Müller-Nurasyid, Niels Grarup, Heather M. Stringham, Eric R. Gamazon, Jaehoon Lee, Yuhui Chen, Robert A. Scott, Jennifer E. Below, Peng Chen, Jinyan Huang, Min Jin Go, Michael L. Stitzel, Dorota Pasko, Stephen C. J. Parker, Tibor V. Varga, Todd Green, Nicola L. Beer, Aaron G. Day-Williams, Teresa Ferreira, Tasha Fingerlin, Momoko Horikoshi, Cheng Hu, Iksoo Huh, Mohammad Kamran Ikram, Bong-Jo Kim, Yongkang Kim, Young Jin Kim, Min-Seok Kwon, Juyoung Lee, Selyeong Lee, Keng-Han Lin, Taylor J. Maxwell, Yoshihiko Nagai, Xu Wang, Ryan P. Welch, Joon Yoon, Weihua Zhang, Nir Barzilai, Benjamin F. Voight, Bok-Ghee Han, Christopher P. Jenkinson, Teemu Kuulasmaa, Johanna Kuusisto, Alisa Manning, Maggie C. Y. Ng, Nicholette D. Palmer, Beverley Balkau, Alena Stančáková, Hanna E. Abboud, Heiner Boeing, Vilmantas Giedraitis, Dorairaj Prabhakaran, Omri Gottesman, James Scott, Jason Carey, Phoenix Kwan, George Grant, Joshua D. Smith, Benjamin M. Neale, Shaun Purcell, Adam S. Butterworth, Joanna M. M. Howson, Heung Man Lee, Yingchang Lu, Soo-Heon Kwak, Wei Zhao, John Danesh, Vincent K. L. Lam, Kyong Soo Park, Danish Saleheen, Wing Yee So, Claudia H. T. Tam, Uzma Afzal, David Aguilar, Rector Arya, Tin Aung, Edmund Chan, Carmen Navarro, Ching-Yu Cheng, Domenico Palli, Adolfo Correa, Joanne E. Curran, Denis Rybin, Vidya S. Farook, Sharon P. Fowler, Barry I. Freedman, Michael Griswold, Daniel Esten Hale, Pamela J. Hicks, Chiea-Chuen Khor, Satish Kumar, Benjamin Lehne, Dorothée Thuillier, Wei Yen Lim, Jianjun Liu, Yvonne T. van der Schouw, Marie Loh, Solomon K. Musani, Sobha Puppala, William R. Scott, Loïc Yengo, Sian-Tsung Tan, Herman A. Taylor Jr, Farook Thameem, Gregory Wilson, Tien Yin Wong, Pål Rasmus Njølstad, Jonathan C. Levy, Massimo Mangino, Lori L. Bonnycastle, Thomas Schwarzmayr, João Fadista, Gabriela L. Surdulescu, Christian Herder, Christopher J. Groves, Thomas Wieland, Jette Bork-Jensen, Ivan Brandslund, Cramer Christensen, Heikki A. Koistinen, Alex S. F. Doney, Leena Kinnunen, Tõnu Esko, Andrew J. Farmer, Liisa Hakaste, Dylan Hodgkiss, Jasmina Kravic, Valeriya Lyssenko, Mette Hollensted, Marit E. Jørgensen, Torben Jørgensen, Claes Ladenvall, Johanne Marie Justesen, Annemari Käräjämäki, Jennifer Kriebel, Wolfgang Rathmann, Lars Lannfelt, Torsten Lauritzen, Narisu Narisu, Allan Linneberg, Olle Melander, Lili Milani, Matt Neville, Marju Orho-Melander, Lu Qi, Qibin Qi, Michael Roden, Olov Rolandsson, Amy Swift, Anders H. Rosengren, Kathleen Stirrups, Andrew R. Wood, Evelin Mihailov, Christine Blancher, Mauricio O. Carneiro, Jared Maguire, Ryan Poplin, Khalid Shakir, Timothy Fennell, Mark DePristo, Martin Hrabé de Angelis, Panos Deloukas, Anette P. Gjesing, Goo Jun, Peter Nilsson, Jacquelyn Murphy, Robert Onofrio, Barbara Thorand, Torben Hansen, Christa Meisinger, Frank B. Hu, Bo Isomaa, Fredrik Karpe, Liming Liang, Annette Peters, Cornelia Huth, Stephen P. O’Rahilly, Colin N. A. Palmer, Oluf Pedersen, Rainer Rauramaa, Jaakko Tuomilehto, Veikko Salomaa, Richard M. Watanabe, Ann-Christine Syvänen, Richard N. Bergman, Dwaipayan Bharadwaj, Erwin P. Bottinger, Yoon Shin Cho, Giriraj R. Chandak, Juliana C. N. Chan, Kee Seng Chia, Mark J. Daly, Shah B. Ebrahim, Claudia Langenberg, Paul Elliott, Kathleen A. Jablonski, Donna M. Lehman, Weiping Jia, Ronald C. W. Ma, Toni I. Pollin, Manjinder Sandhu, Nikhil Tandon, Philippe Froguel, Inês Barroso, Yik Ying Teo, Eleftheria Zeggini, Ruth J. F. Loos, Kerrin S. Small, Janina S. Ried, Ralph A. DeFronzo, Harald Grallert, Benjamin Glaser, Andres Metspalu, Nicholas J. Wareham, Mark Walker, Eric Banks, Christian Gieger, Erik Ingelsson, Hae Kyung Im, Thomas Illig, Paul W. Franks, Gemma Buck, Joseph Trakalo, David Buck, Inga Prokopenko, Reedik Mägi, Lars Lind, Yossi Farjoun, Katharine R. Owen, Anna L. Gloyn, Konstantin Strauch, Tiinamaija Tuomi, Jaspal Singh Kooner, Jong-Young Lee, Taesung Park, Peter Donnelly, Andrew D. Morris, Andrew T. Hattersley, Donald W. Bowden, Francis S. Collins, Gil Atzmon, John C. Chambers, Timothy D. Spector, Markku Laakso, Tim M. Strom, Graeme I. Bell, John Blangero, Ravindranath Duggirala, E. Shyong Tai, Gilean McVean, Craig L. Hanis, James G. Wilson, Mark Seielstad, Timothy M. Frayling, James B. Meigs, Nancy J. Cox, Rob Sladek, Eric S. Lander, Stacey Gabriel, Noël P. Burtt, Karen L. Mohlke, Thomas Meitinger, Leif Groop, Goncalo Abecasis, Jose C. Florez, Laura J. Scott, Andrew P. Morris, Hyun Min Kang, Michael Boehnke, David Altshuler, and Mark I. McCarthy. The genetic architecture of type 2 diabetes. Nature, 536(7614):41–47, 2016. ISSN 1476-4687. doi: 10.1038/nature18642. URL https://www.nature.com/articles/nature18642. Number: 7614 Publisher: Nature Publishing Group.

[4] Peter M. Visscher, Matthew A. Brown, Mark I. McCarthy, and Jian Yang. Five years of GWAS discovery. American Journal of Human Genetics, 90(1):7–24, 2012. ISSN 0002-9297. doi: 10.1016/j.ajhg.2011.11.029. URL https://www.ncbi.nlm.nih.gov/pmc/articles/PMC3257326/.

[5] Peter M. Visscher, Naomi R. Wray, Qian Zhang, Pamela Sklar, Mark I. McCarthy, Matthew A. Brown, and Jian Yang. 10 years of GWAS discovery: Biology, function, and translation. The American Journal of Human Genetics, 101(1):5–22, 2017. ISSN 0002-9297, 1537-6605. doi: 10.1016/j.ajhg.2017.06.005. URL https://www.cell.com/ajhg/abstract/S0002-9297(17)30240-9. Publisher: Elsevier.

[6] Ruth J. F. Loos. 15 years of genome-wide association studies and no signs of slowing down. Nature Communications, 11(1):5900, 2020. ISSN 2041-1723. doi: 10.1038/s41467-020-19653-5. URL https://www.nature.com/articles/s41467-020-19653-5. Number: 1 Publisher: Nature Publishing Group.

[7] Elena S. Gusareva, Jean-Claude Twizere, Kristel Sleegers, Pierre Dourlen, Jose F. Abisambra, Shelby Meier, Ryan Cloyd, Blaine Weiss, Bart Dermaut, Kyrylo Bessonov, Sven J. van der Lee, Minerva M. Carrasquillo, Yuriko Katsumata, Majid Cherkaoui, Bob Asselbergh, M. Arfan Ikram, Richard Mayeux, Lindsay A. Farrer, Jonathan L. Haines, Margaret A. Pericak-Vance, Gerard D. Schellenberg, Genetic and Environmental Risk in Alzheimer’s Disease 1 consortium (GERAD1), Alzheimer’s Disease Genetics Consortium (ADGC), European Alzheimer Disease Initiative Investigators (EADI1 Consortium), Rebecca Sims, Julie Williams, Philippe Amouyel, Cornelia M. van Duijn, Nilüfer Ertekin-Taner, Christine Van Broeckhoven, Franck Dequiedt, David W. Fardo, Jean-Charles Lambert, and Kristel Van Steen. Male-specific epistasis between WWC1 and TLN2 genes is associated with alzheimer’s disease. Neurobiology of Aging, 72:188.e3–188.e12, 2018. ISSN 1558-1497. doi: 10.1016/j.neurobiolaging.2018.08.001.

[8] Yohei Kirino, George Bertsias, Yoshiaki Ishigatsubo, Nobuhisa Mizuki, Ilknur Tugal-Tutkun, Emire Seyahi, Yilmaz Ozyazgan, F. Sevgi Sacli, Burak Erer, Hidetoshi Inoko, Zeliha Emrence, Atilla Cakar, Neslihan Abaci, Duran Ustek, Colleen Satorius, Atsuhisa Ueda, Mitsuhiro Takeno, Yoonhee Kim, Geryl M. Wood, Michael J. Ombrello, Akira Meguro, Ahmet Gül, Elaine F. Remmers, and Daniel L. Kastner. Genome-wide association analysis identifies new susceptibility loci for behçet’s disease and epistasis between HLA-b*51 and ERAP1. Nature Genetics, 45(2):202–207, 2013. ISSN 1546-1718. doi: 10.1038/ng.2520.

[9] Sha Tao, Junjie Feng, Timothy Webster, Guangfu Jin, Fang-Chi Hsu, Shyh-Huei Chen, Seong-Tae Kim, Zhong Wang, Zheng Zhang, Siqun L. Zheng, William B. Isaacs, Jianfeng Xu, and Jielin Sun. Genome-wide two-locus epistasis scans in prostate cancer using two european populations. Human Genetics, 131(7):1225–1234, 2012. ISSN 1432-1203. doi: 10.1007/s00439-012-1148-4.

[10] Wen-Hua Wei, Gib Hemani, Attila Gyenesei, Veronique Vitart, Pau Navarro, Caroline Hayward, Claudia P. Cabrera, Jennifer E. Huffman, Sara A. Knott, Andrew A. Hicks, Igor Rudan, Peter P. Pramstaller, Sarah H. Wild, James F. Wilson, Harry Campbell, Nicholas D. Hastie, Alan F. Wright, and Chris S. Haley. Genome-wide analysis of epistasis in body mass index using multiple human populations. European journal of human genetics: EJHG, 20(8):857–862, 2012. ISSN 1476-5438. doi: 10.1038/ejhg.2012.17.

[11] Gang Chen, Futao Zhang, Wenda Xue, Ruyan Wu, Haiming Xu, Kesheng Wang, and Jun Zhu. An association study revealed substantial effects of dominance, epistasis and substance dependence co-morbidity on alcohol dependence symptom count. Addiction Biology, 22(6):1475–1485, 2017. ISSN 1369-1600. doi: 10.1111/adb.12402.

[12] Gregory Darnell, Samuel Pattillo Smith, Dana Udwin, Sohini Ramachandran, and Lorin Crawford. Partitioning tagged non-additive genetic effects in summary statistics provides evidence of pervasive epistasis in complex traits, 2022. URL https://www.biorxiv.org/content/10.1101/2022.07.21.501001v3. Pages: 2022.07.21.501001 Section: New Results.

[13] A. C. Peripato, R. A. De Brito, S. R. Matioli, L. S. Pletscher, T. T. Vaughn, and J. M. Cheverud. Epistasis affecting litter size in mice. Journal of Evolutionary Biology, 17(3):593–602, 2004. ISSN 1420-9101. doi: 10.1111/j.1420-9101.2004.00702.x.

[14] Amy Hin Yan Tong, Guillaume Lesage, Gary D. Bader, Huiming Ding, Hong Xu, Xiaofeng Xin, James Young, Gabriel F. Berriz, Renee L. Brost, Michael Chang, YiQun Chen, Xin Cheng, Gordon Chua, Helena Friesen, Debra S. Goldberg, Jennifer Haynes, Christine Humphries, Grace He, Shamiza Hussein, Lizhu Ke, Nevan Krogan, Zhijian Li, Joshua N. Levinson, Hong Lu, Patrice Ménard, Christella Munyana, Ainslie B. Parsons, Owen Ryan, Raffi Tonikian, Tania Roberts, Anne-Marie Sdicu, Jesse Shapiro, Bilal Sheikh, Bernhard Suter, Sharyl L. Wong, Lan V. Zhang, Hongwei Zhu, Christopher G. Burd, Sean Munro, Chris Sander, Jasper Rine, Jack Greenblatt, Matthias Peter, Anthony Bretscher, Graham Bell, Frederick P. Roth, Grant W. Brown, Brenda Andrews, Howard Bussey, and Charles Boone. Global Mapping of the Yeast Genetic Interaction Network. Science, 303(5659):808–813, February 2004. doi: 10.1126/science.1091317.

[15] Rachel B. Brem, John D. Storey, Jacqueline Whittle, and Leonid Kruglyak. Genetic interactions between polymorphisms that affect gene expression in yeast. Nature, 436(7051):701–703, August 2005. ISSN 1476-4687. doi: 10.1038/nature03865.

[16] Adam M. Deutschbauer and Ronald W. Davis. Quantitative trait loci mapped to single-nucleotide resolution in yeast. Nature Genetics, 37(12):1333–1340, December 2005. ISSN 1546-1718. doi: 10.1038/ng1674.

[17] Juergen Kroymann and Thomas Mitchell-Olds. Epistasis and balanced polymorphism influencing complex trait variation. Nature, 435(7038):95–98, May 2005. ISSN 1476-4687. doi: 10.1038/nature03480.

[18] Sean R. Collins, Maya Schuldiner, Nevan J. Krogan, and Jonathan S. Weissman. A strategy for extracting and analyzing large-scale quantitative epistatic interaction data. Genome Biology, 7(7): R63, July 2006. ISSN 1474-760X. doi: 10.1186/gb-2006-7-7-r63.

[19] Ben Lehner, Catriona Crombie, Julia Tischler, Angelo Fortunato, and Andrew G. Fraser. Systematic mapping of genetic interactions in Caenorhabditis elegans identifies common modifiers of diverse signaling pathways. Nature Genetics, 38(8):896–903, August 2006. ISSN 1546-1718. doi: 10.1038/ng1844.

[20] Robert P. St Onge, Ramamurthy Mani, Julia Oh, Michael Proctor, Eula Fung, Ronald W. Davis, Corey Nislow, Frederick P. Roth, and Guri Giaever. Systematic pathway analysis using high-resolution fitness profiling of combinatorial gene deletions. Nature Genetics, 39(2):199–206, February 2007. ISSN 1546-1718. doi: 10.1038/ng1948.

[21] Adam M. Wentzell, Heather C. Rowe, Bjarne Gram Hansen, Carla Ticconi, Barbara Ann Halkier, and Daniel J. Kliebenstein. Linking Metabolic QTLs with Network and cis-eQTLs Controlling Biosynthetic Pathways. PLOS Genetics, 3(9):e162, September 2007. ISSN 1553-7404. doi: 10.1371/journal.pgen.0030162.

[22] Haifeng Shao, Lindsay C. Burrage, David S. Sinasac, Annie E. Hill, Sheila R. Ernest, William O’Brien, Hayden-William Courtland, Karl J. Jepsen, Andrew Kirby, E. J. Kulbokas, Mark J. Daly, Karl W. Broman, Eric S. Lander, and Joseph H. Nadeau. Genetic architecture of complex traits: Large phenotypic effects and pervasive epistasis. Proceedings of the National Academy of Sciences, 105(50):19910–19914, December 2008. doi: 10.1073/pnas.0810388105.

[23] Jonathan Flint and Trudy F.C. Mackay. Genetic architecture of quantitative traits in mice, flies, and humans. Genome Research, 19(5):723–733, May 2009. ISSN 1088-9051. doi: 10.1101/gr.086660.108.

[24] Justin Gerke, Kim Lorenz, and Barak Cohen. Genetic Interactions Between Transcription Factors Cause Natural Variation in Yeast. Science, 323(5913):498–501, January 2009. doi: 10.1126/science.1166426.

[25] Michael Costanzo, Anastasia Baryshnikova, Jeremy Bellay, Yungil Kim, Eric D. Spear, Carolyn S. Sevier, Huiming Ding, Judice L.Y. Koh, Kiana Toufighi, Sara Mostafavi, Jeany Prinz, Robert P. St. Onge, Benjamin VanderSluis, Taras Makhnevych, Franco J. Vizeacoumar, Solmaz Alizadeh, Sondra Bahr, Renee L. Brost, Yiqun Chen, Murat Cokol, Raamesh Deshpande, Zhijian Li, Zhen-Yuan Lin, Wendy Liang, Michaela Marback, Jadine Paw, Bryan-Joseph San Luis, Ermira Shuteriqi, Amy Hin Yan Tong, Nydia van Dyk, Iain M. Wallace, Joseph A. Whitney, Matthew T. Weirauch, Guoqing Zhong, Hongwei Zhu, Walid A. Houry, Michael Brudno, Sasan Ragibizadeh, Balázs Papp, Csaba Pál, Frederick P. Roth, Guri Giaever, Corey Nislow, Olga G. Troyanskaya, Howard Bussey, Gary D. Bader, Anne-Claude Gingras, Quaid D. Morris, Philip M. Kim, Chris A. Kaiser, Chad L. Myers, Brenda J. Andrews, and Charles Boone. The Genetic Landscape of a Cell. Science, 327(5964):425–431, January 2010. doi: 10.1126/science.1180823.

[26] Xionglei He, Wenfeng Qian, Zhi Wang, Ying Li, and Jianzhi Zhang. Prevalent positive epistasis in Escherichia coli and Saccharomyces cerevisiae metabolic networks. Nature Genetics, 42(3):272–276, March 2010. ISSN 1546-1718. doi: 10.1038/ng.524.

[27] Thomas Horn, Thomas Sandmann, Bernd Fischer, Elin Axelsson, Wolfgang Huber, and Michael Boutros. Mapping of signaling networks through synthetic genetic interaction analysis by RNAi. Nature Methods, 8(4):341–346, April 2011. ISSN 1548-7105. doi: 10.1038/nmeth.1581.

[28] Joseph P Jarvis and James M Cheverud. Mapping the Epistatic Network Underlying Murine Reproductive Fatpad Variation. Genetics, 187(2):597–610, February 2011. ISSN 1943-2631. doi: 10.1534/genetics.110.123505.

[29] Larry Leamy, Ryan Gordon, and Daniel Pomp. Sex-, Diet-, and Cancer-Dependent Epistatic Effects on Complex Traits in Mice. Frontiers in Genetics, 2, 2011. ISSN 1664-8021.

[30] Mats Pettersson, Francois Besnier, Paul B. Siegel, and Örjan Carlborg. Replication and Explorations of High-Order Epistasis Using a Large Advanced Intercross Line Pedigree. PLOS Genetics, 7(7):e1002180, July 2011. ISSN 1553-7404. doi: 10.1371/journal.pgen.1002180.

[31] Balázs Szappanos, Károly Kovács, Béla Szamecz, Frantisek Honti, Michael Costanzo, Anastasia Baryshnikova, Gabriel Gelius-Dietrich, Martin J. Lercher, Márk Jelasity, Chad L. Myers, Brenda J. Andrews, Charles Boone, Stephen G. Oliver, Csaba Pál, and Balázs Papp. An integrated approach to characterize genetic interaction networks in yeast metabolism. Nature Genetics, 43(7):656–662, July 2011. ISSN 1546-1718. doi: 10.1038/ng.846.

[32] Bryn E Gaertner, Michelle D Parmenter, Matthew V Rockman, Leonid Kruglyak, and Patrick C Phillips. More Than the Sum of Its Parts: A Complex Epistatic Network Underlies Natural Variation in Thermal Preference Behavior in Caenorhabditis elegans. Genetics, 192(4):1533–1542, December 2012. ISSN 1943-2631. doi: 10.1534/genetics.112.142877.

[33] Joshua S. Bloom, Ian M. Ehrenreich, Wesley T. Loo, Thúy-Lan Võ Lite, and Leonid Kruglyak. Finding the sources of missing heritability in a yeast cross. Nature, 494(7436):234–237, February 2013. ISSN 1476-4687. doi: 10.1038/nature11867.

[34] Sudarshan Chari and Ian Dworkin. The Conditional Nature of Genetic Interactions: The Consequences of Wild-Type Backgrounds on Mutational Interactions in a Genome-Wide Modifier Screen. PLOS Genetics, 9(8):e1003661, August 2013. ISSN 1553-7404. doi: 10.1371/journal.pgen.1003661.

[35] Patrick J. Monnahan and John K. Kelly. Epistasis Is a Major Determinant of the Additive Genetic Variance in Mimulus guttatus. PLOS Genetics, 11(5):e1005201, May 2015. ISSN 1553-7404. doi: 10.1371/journal.pgen.1005201.

[36] Wen Huang and Trudy F. C. Mackay. The Genetic Architecture of Quantitative Traits Cannot Be Inferred from Variance Component Analysis. PLOS Genetics, 12(11):e1006421, November 2016. ISSN 1553-7404. doi: 10.1371/journal.pgen.1006421.

[37] Lorin Crawford, Kris C. Wood, Xiang Zhou, and Sayan Mukherjee. Bayesian approximate kernel regression with variable selection. Journal of the American Statistical Association, 113 (524):1710–1721, 2018. ISSN 0162-1459, 1537-274X. doi: 10.1080/01621459.2017.1361830. URL https://www.tandfonline.com/doi/full/10.1080/01621459.2017.1361830.

[38] Daniel M. Weinreich, Nigel F. Delaney, Mark A. DePristo, and Daniel L. Hartl. Darwinian evolution can follow only very few mutational paths to fitter proteins. Science, 312(5770):111–114, 2006. doi: 10.1126/science.1123539. URL https://www.science.org/doi/10.1126/science.1123539. Publisher: American Association for the Advancement of Science.

[39] Simon K. G. Forsberg, Joshua S. Bloom, Meru J. Sadhu, Leonid Kruglyak, and Örjan Carlborg. Accounting for genetic interactions improves modeling of individual quantitative trait phenotypes in yeast. Nature Genetics, 49(4):497–503, April 2017. ISSN 1546-1718. doi: 10.1038/ng.3800.

[40] Juannan Zhou, Mandy S. Wong, Wei-Chia Chen, Adrian R. Krainer, Justin B. Kinney, and David M. McCandlish. Higher-order epistasis and phenotypic prediction. Proceedings of the National Academy of Sciences, 119(39):e2204233119, 2022. doi: 10.1073/pnas.2204233119. URL https://www.pnas.org/doi/full/10.1073/pnas.2204233119. Publisher: Proceedings of the National Academy of Sciences.

[41] Frank J. Poelwijk, Michael Socolich, and Rama Ranganathan. Learning the pattern of epistasis linking genotype and phenotype in a protein. Nature Communications, 10(1):4213, September 2019. ISSN 2041-1723. doi: 10.1038/s41467-019-12130-8.

[42] Daniel E. Runcie, Jiayi Qu, Hao Cheng, and Lorin Crawford. MegaLMM: Mega-scale linear mixed models for genomic predictions with thousands of traits. Genome Biology, 22(1):213, July 2021. ISSN 1474-760X. doi: 10.1186/s13059-021-02416-w.

[43] Patricio R Muñoz, Marcio F R Resende, Jr, Salvador A Gezan, Marcos Deon Vilela Resende, Gustavo de los Campos, Matias Kirst, Dudley Huber, and Gary F Peter. Unraveling Additive from Nonadditive Effects Using Genomic Relationship Matrices. Genetics, 198(4):1759–1768, December 2014. ISSN 1943-2631. doi: 10.1534/genetics.114.171322.

[44] Yong Jiang and Jochen C. Reif. Modeling Epistasis in Genomic Selection. Genetics, 201(2):759–768, October 2015. ISSN 0016-6731, 1943-2631. doi: 10.1534/genetics.115.177907.

[45] Anthony D. Long and Charles H. Langley. The Power of Association Studies to Detect the Contribution of Candidate Genetic Loci to Variation in Complex Traits. Genome Research, 9(8):720–731, August 1999. ISSN 1088-9051.

[46] Jonathan Marchini, Peter Donnelly, and Lon R. Cardon. Genome-wide strategies for detecting multiple loci that influence complex diseases. Nature Genetics, 37(4):413–417, April 2005. ISSN 1546-1718. doi: 10.1038/ng1537.

[47] William G. Hill, Michael E. Goddard, and Peter M. Visscher. Data and Theory Point to Mainly Additive Genetic Variance for Complex Traits. PLOS Genetics, 4(2):e1000008, February 2008. ISSN 1553-7404. doi: 10.1371/journal.pgen.1000008.

[48] James F. Crow. On epistasis: Why it is unimportant in polygenic directional selection. Philosophical Transactions of the Royal Society B: Biological Sciences, 365(1544):1241–1244, April 2010. ISSN 0962-8436. doi: 10.1098/rstb.2009.0275.

[49] Or Zuk, Eliana Hechter, Shamil R. Sunyaev, and Eric S. Lander. The mystery of missing heritability: Genetic interactions create phantom heritability. Proceedings of the National Academy of Sciences, 109(4):1193–1198, January 2012. doi: 10.1073/pnas.1119675109.

[50] Valentin Hivert, Julia Sidorenko, Florian Rohart, Michael E. Goddard, Jian Yang, Naomi R. Wray, Loic Yengo, and Peter M. Visscher. Estimation of non-additive genetic variance in human complex traits from a large sample of unrelated individuals. The American Journal of Human Genetics, 108(5):786–798, May 2021. ISSN 0002-9297. doi: 10.1016/j.ajhg.2021.02.014.

[51] Ali Pazokitoroudi, Alec M. Chiu, Kathryn S. Burch, Bogdan Pasaniuc, and Sriram Sankararaman. Quantifying the contribution of dominance deviation effects to complex trait variation in biobank-scale data. The American Journal of Human Genetics, 108(5):799–808, May 2021. ISSN 0002-9297. doi: 10.1016/j.ajhg.2021.03.018.

[52] Pierrick Wainschtein, Deepti Jain, Zhili Zheng, L. Adrienne Cupples, Aladdin H. Shadyab, Barbara McKnight, Benjamin M. Shoemaker, Braxton D. Mitchell, Bruce M. Psaty, Charles Kooperberg, Ching-Ti Liu, Christine M. Albert, Dan Roden, Daniel I. Chasman, Dawood Darbar, Donald M. Lloyd-Jones, Donna K. Arnett, Elizabeth A. Regan, Eric Boerwinkle, Jerome I. Rotter, Jeffrey R. O’Connell, Lisa R. Yanek, Mariza de Andrade, Matthew A. Allison, Merry-Lynn N. McDonald, Mina K. Chung, Myriam Fornage, Nathalie Chami, Nicholas L. Smith, Patrick T. Ellinor, Ramachandran S. Vasan, Rasika A. Mathias, Ruth J. F. Loos, Stephen S. Rich, Steven A. Lubitz, Susan R. Heckbert, Susan Redline, Xiuqing Guo, Y.-D. Ida Chen, Cecelia A. Laurie, Ryan D. Hernandez, Stephen T. McGarvey, Michael E. Goddard, Cathy C. Laurie, Kari E. North, Leslie A. Lange, Bruce S. Weir, Loic Yengo, Jian Yang, and Peter M. Visscher. Assessing the contribution of rare variants to complex trait heritability from whole-genome sequence data. Nature Genetics, 54(3):263–273, March 2022. ISSN 1546-1718. doi: 10.1038/s41588-021-00997-7.

[53] Andrew Anand Brown, Alfonso Buil, Ana Viñuela, Tuuli Lappalainen, Hou-Feng Zheng, J Brent Richards, Kerrin S Small, Timothy D Spector, Emmanouil T Dermitzakis, and Richard Durbin. Genetic interactions affecting human gene expression identified by variance association mapping. eLife, 3:e01381, April 2014. ISSN 2050-084X. doi: 10.7554/eLife.01381.

[54] Gang Fang, Wen Wang, Vanja Paunic, Hamed Heydari, Michael Costanzo, Xiaoye Liu, Xiaotong Liu, Benjamin VanderSluis, Benjamin Oately, Michael Steinbach, Brian Van Ness, Eric E. Schadt, Nathan D. Pankratz, Charles Boone, Vipin Kumar, and Chad L. Myers. Discovering genetic interactions bridging pathways in genome-wide association studies. Nature Communications, 10(1): 4274, September 2019. ISSN 2041-1723. doi: 10.1038/s41467-019-12131-7.

[55] Joris van de Haar, Sander Canisius, Michael K. Yu, Emile E. Voest, Lodewyk F. A. Wessels, and Trey Ideker. Identifying Epistasis in Cancer Genomes: A Delicate Affair. Cell, 177(6):1375–1383, May 2019. ISSN 0092-8674. doi: 10.1016/j.cell.2019.05.005.

[56] Brooke Sheppard, Nadav Rappoport, Po-Ru Loh, Stephan J. Sanders, Noah Zaitlen, and Andy Dahl. A model and test for coordinated polygenic epistasis in complex traits. Proceedings of the National Academy of Sciences, 118(15):e1922305118, April 2021. doi: 10.1073/pnas.1922305118.

[57] Roshni A. Patel, Shaila A. Musharoff, Jeffrey P. Spence, Harold Pimentel, Catherine Tcheandjieu, Hakhamanesh Mostafavi, Nasa Sinnott-Armstrong, Shoa L. Clarke, Courtney J. Smith, Peter P. Durda, Kent D. Taylor, Russell Tracy, Yongmei Liu, W. Craig Johnson, Francois Aguet, Kristin G. Ardlie, Stacey Gabriel, Josh Smith, Deborah A. Nickerson, Stephen S. Rich, Jerome I. Rotter, Philip S. Tsao, Themistocles L. Assimes, and Jonathan K. Pritchard. Genetic interactions drive heterogeneity in causal variant effect sizes for gene expression and complex traits. The American Journal of Human Genetics, 109(7):1286–1297, July 2022. ISSN 0002-9297. doi: 10.1016/j.ajhg.2022.05.014.

[58] Evan E. Eichler, Jonathan Flint, Greg Gibson, Augustine Kong, Suzanne M. Leal, Jason H. Moore, and Joseph H. Nadeau. Missing heritability and strategies for finding the underlying causes of complex disease. Nature Reviews Genetics, 11(6):446–450, 2010. ISSN 1471-0064. doi: 10.1038/nrg2809. URL https://www.nature.com/articles/nrg2809. Number: 6 Publisher: Nature Publishing Group.

[59] Elena S. Gusareva, Minerva M. Carrasquillo, Céline Bellenguez, Elise Cuyvers, Samuel Colon, Neill R. Graff-Radford, Ronald C. Petersen, Dennis W. Dickson, Jestinah M. Mahachie John, Kyrylo Bessonov, Christine Van Broeckhoven, GERAD1 Consortium, Denise Harold, Julie Williams, Philippe Amouyel, Kristel Sleegers, Nilüfer Ertekin-Taner, Jean-Charles Lambert, and Kristel Van Steen. Genome-wide association interaction analysis for alzheimer’s disease. Neurobiology of Aging, 35(11):2436–2443, 2014. ISSN 1558-1497. doi: 10.1016/j.neurobiolaging.2014.05.014.

[60] Elena S. Gusareva and Kristel Van Steen. Practical aspects of genome-wide association interaction analysis. Human Genetics, 133(11):1343–1358, 2014. ISSN 1432-1203. doi: 10.1007/s00439-014-1480-y. URL https://doi.org/10.1007/s00439-014-1480-y.

[61] Amy Wanstrat and Edward Wakeland. The genetics of complex autoimmune diseases: non-MHC susceptibility genes. Nature Immunology, 2(9):802–809, 2001. ISSN 1529-2916. doi: 10.1038/ni0901-802. URL https://www.nature.com/articles/ni0901-802. Number: 9 Publisher: Nature Publishing Group.

[62] Shaun Purcell, Benjamin Neale, Kathe Todd-Brown, Lori Thomas, Manuel A. R. Ferreira, David Bender, Julian Maller, Pamela Sklar, Paul I. W. de Bakker, Mark J. Daly, and Pak C. Sham. PLINK: A tool set for whole-genome association and population-based linkage analyses. American Journal of Human Genetics, 81(3):559–575, 2007. ISSN 0002-9297. URL https://www.ncbi.nlm.nih.gov/pmc/articles/PMC1950838/.

[63] Thierry Schüpbach, Ioannis Xenarios, Sven Bergmann, and Karen Kapur. FastEpistasis: a high performance computing solution for quantitative trait epistasis. Bioinformatics, 26(11):1468–1469, 2010. ISSN 1367-4803. doi: 10.1093/bioinformatics/btq147. URL https://www.ncbi.nlm.nih.gov/pmc/articles/PMC2872003/.

[64] Li Ma, Andrew G. Clark, and Alon Keinan. Gene-based testing of interactions in association studies of quantitative traits. PLOS Genetics, 9(2):e1003321, 2013. ISSN 1553-7404. doi: 10.1371/journal.pgen.1003321. URL https://journals.plos.org/plosgenetics/article?id=10.1371/journal.pgen.1003321. Publisher: Public Library of Science.

[65] Snehit Prabhu and Itsik Pe’er. Ultrafast genome-wide scan for SNP–SNP interactions in common complex disease. Genome Research, 22(11):2230–2240, 2012. ISSN 1088-9051, 1549-5469. doi: 10.1101/gr.137885.112. URL https://genome.cshlp.org/content/22/11/2230. Company: Cold Spring Harbor Laboratory Press Distributor: Cold Spring Harbor Laboratory Press Institution: Cold Spring Harbor Laboratory Press Label: Cold Spring Harbor Laboratory Press Publisher: Cold Spring Harbor Lab.

[66] Juan Pablo Lewinger, John L. Morrison, Duncan C. Thomas, Cassandra E. Murcray, David V. Conti, Dalin Li, and W. James Gauderman. Efficient Two-Step Testing of Gene-Gene Interactions in Genome-Wide Association Studies. Genetic Epidemiology, 37(5):440–451, 2013. ISSN 1098-2272. doi: 10.1002/gepi.21720.

[67] Yingjie Guo, Honghong Cheng, Zhian Yuan, Zhen Liang, Yang Wang, and Debing Du. Testing Gene-Gene Interactions Based on a Neighborhood Perspective in Genome-wide Association Studies. Frontiers in Genetics, 12, 2021. ISSN 1664-8021.

[68] Xiang Wan, Can Yang, Qiang Yang, Hong Xue, Xiaodan Fan, Nelson L.S. Tang, and Weichuan Yu. BOOST: A fast approach to detecting gene-gene interactions in genome-wide case-control studies. American Journal of Human Genetics, 87(3):325–340, 2010. ISSN 0002-9297. doi: 10.1016/j.ajhg.2010.07.021. URL https://www.ncbi.nlm.nih.gov/pmc/articles/PMC2933337/.

[69] François Van Lishout, Jestinah M. Mahachie John, Elena S. Gusareva, Victor Urrea, Isabelle Cleynen, Emilie Théâtre, Benoît Charloteaux, Malu Luz Calle, Louis Wehenkel, and Kristel Van Steen. An efficient algorithm to perform multiple testing in epistasis screening. BMC Bioinformatics, 14(1):138, 2013. ISSN 1471-2105. doi: 10.1186/1471-2105-14-138. URL https://doi.org/10.1186/1471-2105-14-138.

[70] Yu Zhang and Jun S. Liu. Bayesian inference of epistatic interactions in case-control studies. Nature Genetics, 39(9):1167–1173, September 2007. ISSN 1546-1718. doi: 10.1038/ng2110.

[71] Yu Zhang, Jing Zhang, and Jun S. Liu. Block-based bayesian epistasis association mapping with application to WTCCC type 1 diabetes data. The Annals of Applied Statistics, 5(3):2052–2077, 2011. ISSN 1932-6157, 1941-7330. doi: 10.1214/11-AOAS469. URL https://projecteuclid.org/journals/annals-of-applied-statistics/volume-5/issue-3/Block-based-Bayesian-epistasis-association-mapping-with/10.1214/11-AOAS469.full. Publisher: Institute of Mathematical Statistics.

[72] Yang Guo, Zhiman Zhong, Chen Yang, Jiangfeng Hu, Yaling Jiang, Zizhen Liang, Hui Gao, and Jianxiao Liu. Epi-GTBN: An approach of epistasis mining based on genetic Tabu algorithm and Bayesian network. BMC Bioinformatics, 20(1):444, August 2019. ISSN 1471-2105. doi: 10.1186/s12859-019-3022-z.

[73] Wanwan Tang, Xuebing Wu, Rui Jiang, and Yanda Li. Epistatic Module Detection for Case-Control Studies: A Bayesian Model with a Gibbs Sampling Strategy. PLOS Genetics, 5(5):e1000464, May 2009. ISSN 1553-7404. doi: 10.1371/journal.pgen.1000464.

[74] Yu-Chuan Chang, June-Tai Wu, Ming-Yi Hong, Yi-An Tung, Ping-Han Hsieh, Sook Wah Yee, Kathleen M. Giacomini, Yen-Jen Oyang, Chien-Yu Chen, Michael W. Weiner, Paul Aisen, Ronald Petersen, Clifford R. Jack, Sara S. Mason, Colleen S. Albers, David Knopman, Kris Johnson, William Jagust, John Q. Trojanowki, Arthur W. Toga, Laurel Beckett, Robert C. Green, Martin R. Farlow, Ann Marie Hake, Brandy R. Matthews, Jared R. Brosch, Scott Herring, Cynthia Hunt, Leslie M. Shaw, Beau Ances, John C. Morris, Maria Carroll, Mary L. Creech, Erin Franklin, Mark A. Mintun, Stacy Schneider, Angela Oliver, Jeffrey Kaye, Joseph Quinn, Lisa Silbert, Betty Lind, Raina Carter, Sara Dolen, Lon S. Schneider, Sonia Pawluczyk, Mauricio Beccera, Liberty Teodoro, Bryan M. Spann, James Brewer, Helen Vanderswag, Adam Fleisher, Pierre Tariot, Anna Burke, Nadira Trncic, Stephanie Reeder, Judith L. Heidebrink, Joanne L. Lord, Rachelle S. Doody, Javier Villanueva-Meyer, Munir Chowdhury, Susan Rountree, Mimi Dang, Yaakov Stern, Lawrence S. Honig, Karen L. Bell, Daniel Marson, Randall Griffith, David Clark, David Geldmacher, John Brockington, Erik Roberson, Marissa Natelson Love, Hillel Grossman, Effie Mitsis, Raj C. Shah, Leyla de Toledo-Morrell, Ranjan Duara, Daniel Varon, Maria T. Greig, Peggy Roberts, Marilyn Albert, Chiadi Onyike, Daniel D’Agostino, Stephanie Kielb, James E. Galvin, Brittany Cerbone, Christina A. Michel, Dana M. Pogorelec, Henry Rusinek, Mony J. de Leon, Lidia Glodzik, Susan De Santi, P. Murali Doraiswamy, Jeffrey R. Petrella, Salvador Borges-Neto, Terence Z. Wong, Edward Coleman, Charles D. Smith, Greg Jicha, Peter Hardy, Partha Sinha, Elizabeth Oates, Gary Conrad, Anton P. Porsteinsson, Bonnie S. Goldstein, Kim Martin, Kelly M. Makino, M. Saleem Ismail, Connie Brand, Ruth A. Mulnard, Gaby Thai, Catherine Mc-AdamsOrtiz, Kyle Womack, Dana Mathews, Mary Quiceno, Allan I. Levey, James J. Lah, Janet S. Cellar, Jeffrey M. Burns, Russell H. Swerdlow, William M. Brooks, Liana Apostolova, Kathleen Tingus, Ellen Woo, Daniel H. S. Silverman, Po H. Lu, George Bartzokis, Neill R. Graff-Radford, Francine Parfitt, Tracy Kendall, Heather Johnson, Christopher H. van Dyck, Richard E. Carson, Martha G. MacAvoy, Pradeep Varma, Howard Chertkow, Howard Bergman, Chris Hosein, Sandra Black, Bojana Stefanovic, Curtis Caldwell, Ging-Yuek Robin Hsiung, Howard Feldman, Benita Mudge, Michele Assaly, Elizabeth Finger, Stephen Pasternack, Irina Rachisky, Dick Trost, Andrew Kertesz, Charles Bernick, Donna Munic, Marek-Marsel Mesulam, Kristine Lipowski, Sandra Weintraub, Borna Bonakdarpour, Diana Kerwin, Chuang-Kuo Wu, Nancy Johnson, Carl Sadowsky, Teresa Villena, Raymond Scott Turner, Kathleen Johnson, Brigid Reynolds, Reisa A. Sperling, Keith A. Johnson, Gad Marshall, Jerome Yesavage, Joy L. Taylor, Barton Lane, Allyson Rosen, Jared Tinklenberg, Marwan N. Sabbagh, Christine M. Belden, Sandra A. Jacobson, Sherye A. Sirrel, Neil Kowall, Ronald Killiany, Andrew E. Budson, Alexander Norbash, Patricia Lynn Johnson, Thomas O. Obisesan, Saba Wolday, Joanne Allard, Alan Lerner, Paula Ogrocki, Curtis Tatsuoka, Parianne Fatica, Evan Fletcher, Pauline Maillard, John Olichney, Charles DeCarli, Owen Carmichael, Smita Kittur, Michael Borrie, T.-Y. Lee, Rob Bartha, Sterling Johnson, Sanjay Asthana, Cynthia M. Carlsson, Steven G. Potkin, Adrian Preda, Dana Nguyen, Vernice Bates, Horacio Capote, Michelle Rainka, Douglas W. Scharre, Maria Kataki, Anahita Adeli, Earl A. Zimmerman, Dzintra Celmins, Alice D. Brown, Godfrey D. Pearlson, Karen Blank, Karen Anderson, Laura A. Flashman, Marc Seltzer, Mary L. Hynes, Robert B. Santulli, Kaycee M. Sink, Leslie Gordineer, Jeff D. Williamson, Pradeep Garg, Franklin Watkins, Brian R. Ott, Henry Querfurth, Geoffrey Tremont, Stephen Salloway, Paul Malloy, Stephen Correia, Howard J. Rosen, Bruce L. Miller, David Perry, Jacobo Mintzer, Kenneth Spicer, David Bachman, Nunzio Pomara, Raymundo Hernando, Antero Sarrael, Norman Relkin, Gloria Chaing, Michael Lin, Lisa Ravdin, Amanda Smith, Balebail Ashok Raj, Kristin Fargher, and for the Alzheimer’s Disease Neuroimaging Initiative. GenEpi: Gene-based epistasis discovery using machine learning. BMC Bioinformatics, 21(1):68, February 2020. ISSN 1471-2105. doi: 10.1186/s12859-020-3368-2.

[75] Paul Fergus, Casimiro Curbelo Montañez, Basma Abdulaimma, Paulo Lisboa, Carl Chalmers, and Beth Pineles. Utilizing Deep Learning and Genome Wide Association Studies for Epistatic-Driven Preterm Birth Classification in African-American Women. IEEE/ACM Transactions on Computational Biology and Bioinformatics, 17(2):668–678, March 2020. ISSN 1557-9964. doi: 10.1109/TCBB.2018.2868667.

[76] Akiko Nagai, Makoto Hirata, Yoichiro Kamatani, Kaori Muto, Koichi Matsuda, Yutaka Kiyohara, Toshiharu Ninomiya, Akiko Tamakoshi, Zentaro Yamagata, Taisei Mushiroda, Yoshinori Murakami, Koichiro Yuji, Yoichi Furukawa, Hitoshi Zembutsu, Toshihiro Tanaka, Yozo Ohnishi, Yusuke Nakamura, Masaki Shiono, Kazuo Misumi, Reiji Kaieda, Hiromasa Harada, Shiro Minami, Mitsuru Emi, Naoya Emoto, Hiroyuki Daida, Katsumi Miyauchi, Akira Murakami, Satoshi Asai, Mitsuhiko Moriyama, Yasuo Takahashi, Tomoaki Fujioka, Wataru Obara, Seijiro Mori, Hideki Ito, Satoshi Nagayama, Yoshio Miki, Akihide Masumoto, Akira Yamada, Yasuko Nishizawa, Ken Kodama, Hiromu Kutsumi, Yoshihisa Sugimoto, Yukihiro Koretsune, Hideo Kusuoka, Hideki Yanai, and Michiaki Kubo. Overview of the BioBank Japan Project: Study design and profile. Journal of Epidemiology, 27(3, Supplement):S2–S8, March 2017. ISSN 0917-5040. doi: 10.1016/j.je.2016.12.005.

[77] Clare Bycroft, Colin Freeman, Desislava Petkova, Gavin Band, Lloyd T. Elliott, Kevin Sharp, Allan Motyer, Damjan Vukcevic, Olivier Delaneau, Jared O’Connell, Adrian Cortes, Samantha Welsh, Alan Young, Mark Effingham, Gil McVean, Stephen Leslie, Naomi Allen, Peter Donnelly, and Jonathan Marchini. The UK Biobank resource with deep phenotyping and genomic data. Nature, 562(7726):203–209, October 2018. ISSN 1476-4687. doi: 10.1038/s41586-018-0579-z.

[78] Gibran Hemani, Athanasios Theocharidis, Wenhua Wei, and Chris Haley. EpiGPU: Exhaustive pairwise epistasis scans parallelized on consumer level graphics cards. Bioinformatics, 27(11): 1462–1465, June 2011. ISSN 1367-4803. doi: 10.1093/bioinformatics/btr172.

[79] Arash Bayat, Brendan Hosking, Yatish Jain, Cameron Hosking, Milindi Kodikara, Daniel Reti, Natalie A. Twine, and Denis C. Bauer. Fast and accurate exhaustive higher-order epistasis search with BitEpi. Scientific Reports, 11(1):15923, August 2021. ISSN 2045-2322. doi: 10.1038/s41598-021-94959-y.

[80] Shijia Zhu and Gang Fang. MatrixEpistasis: Ultrafast, exhaustive epistasis scan for quantitative traits with covariate adjustment. Bioinformatics, 34(14):2341–2348, July 2018. ISSN 1367-4803. doi: 10.1093/bioinformatics/bty094.

[81] Lorin Crawford, Ping Zeng, Sayan Mukherjee, and Xiang Zhou. Detecting epistasis with the marginal epistasis test in genetic mapping studies of quantitative traits. PLOS Genetics, 13(7): e1006869, 2017. ISSN 1553-7404. doi: 10.1371/journal.pgen.1006869. URL https://dx.plos.org/10.1371/journal.pgen.1006869.

[82] Lorin Crawford and Xiang Zhou. Genome-wide marginal epistatic association mapping in case-control studies, 2018. URL https://www.biorxiv.org/content/10.1101/374983v1. Section: New Results Type: article.

[83] Michael C. Turchin, Gregory Darnell, Lorin Crawford, and Sohini Ramachandran. Pathway analysis within multiple human ancestries reveals novel signals for epistasis in complex traits, 2020. URL https://www.biorxiv.org/content/10.1101/2020.09.24.312421v1. Pages: 2020.09.24.312421 Section: New Results.

[84] Rachel Moore, Francesco Paolo Casale, Marc Jan Bonder, Danilo Horta, Lude Franke, Inês Barroso, and Oliver Stegle. A linear mixed-model approach to study multivariate gene–environment interactions. Nature Genetics, 51(1):180–186, 2019. ISSN 1546-1718. doi: 10.1038/s41588-018-0271-0. URL https://www.nature.com/articles/s41588-018-0271-0. Number: 1 Publisher: Nature Publishing Group.

[85] Matthew Kerin and Jonathan Marchini. Inferring gene-by-environment interactions with a bayesian whole-genome regression model. The American Journal of Human Genetics, 107(4):698–713, 2020. ISSN 00029297. doi: 10.1016/j.ajhg.2020.08.009. URL https://linkinghub.elsevier.com/retrieve/pii/S0002929720302779.

[86] Xiang Zhou and Matthew Stephens. Efficient multivariate linear mixed model algorithms for genome-wide association studies. Nature Methods, 11(4):407–409, 2014. ISSN 1548-7105. doi: 10.1038/nmeth.2848. URL https://www.nature.com/articles/nmeth.2848. Number: 4 Publisher: Nature Publishing Group.

[87] Xiang Zhou. A unified framework for variance component estimation with summary statistics in genome-wide association studies. Annals of Applied Statistics, 11(4):2027–2051, 2017. ISSN 1932-6157, 1941-7330. doi: 10.1214/17-AOAS1052. URL https://projecteuclid.org/euclid.aoas/1514430276. Publisher: Institute of Mathematical Statistics.

[88] Angela M Phillips, Katherine R Lawrence, Alief Moulana, Thomas Dupic, Jeffrey Chang, Milo S Johnson, Ivana Cvijovic, Thierry Mora, Aleksandra M Walczak, and Michael M Desai. Binding affinity landscapes constrain the evolution of broadly neutralizing anti-influenza antibodies. eLife, 10:e71393, 2021. ISSN 2050-084X. doi: 10.7554/eLife.71393. URL https://doi.org/10.7554/eLife.71393. Publisher: eLife Sciences Publications, Ltd.

[89] William Valdar, Jonathan Flint, and Richard Mott. Simulating the collaborative cross: power of quantitative trait loci detection and mapping resolution in large sets of recombinant inbred strains of mice. Genetics, 172(3):1783–1797, 2006. ISSN 0016-6731. doi: 10.1534/genetics.104.039313.

[90] William Valdar, Leah C. Solberg, Dominique Gauguier, Stephanie Burnett, Paul Klenerman, William O. Cookson, Martin S. Taylor, J. Nicholas P. Rawlins, Richard Mott, and Jonathan Flint. Genome-wide genetic association of complex traits in heterogeneous stock mice. Nature Genetics, 38(8):879–887, 2006. ISSN 1546-1718. doi: 10.1038/ng1840. URL https://www.nature.com/articles/ng1840. Number: 8 Publisher: Nature Publishing Group.

[91] Wellcome trust centre for human genetics - mouse resources. URL http://mtweb.cs.ucl.ac.uk/mus/www/mouse/index.shtml.

[92] Brendan K. Bulik-Sullivan, Po-Ru Loh, Hilary K. Finucane, Stephan Ripke, Jian Yang, Nick Patterson, Mark J. Daly, Alkes L. Price, and Benjamin M. Neale. LD Score regression distinguishes confounding from polygenicity in genome-wide association studies. Nature Genetics, 47(3):291–295, March 2015. ISSN 1546-1718. doi: 10.1038/ng.3211.

[93] Jian Yang, Beben Benyamin, Brian P. McEvoy, Scott Gordon, Anjali K. Henders, Dale R. Nyholt, Pamela A. Madden, Andrew C. Heath, Nicholas G. Martin, Grant W. Montgomery, Michael E. Goddard, and Peter M. Visscher. Common SNPs explain a large proportion of the heritability for human height. Nature Genetics, 42(7):565–569, July 2010. ISSN 1546-1718. doi: 10.1038/ng.608.

[94] Michael C. Wu, Seunggeun Lee, Tianxi Cai, Yun Li, Michael Boehnke, and Xihong Lin. Rare-variant association testing for sequencing data with the sequence kernel association test. American Journal of Human Genetics, 89(1):82–93, 2011. ISSN 0002-9297. doi: 10.1016/j.ajhg.2011.05.029. URL https://www.ncbi.nlm.nih.gov/pmc/articles/PMC3135811/.

[95] Xiang Zhou, Peter Carbonetto, and Matthew Stephens. Polygenic modeling with bayesian sparse linear mixed models. PLOS Genetics, 9(2):e1003264, 2013. ISSN 1553-7404. doi: 10.1371/journal.pgen.1003264. URL https://journals.plos.org/plosgenetics/article?id=10.1371/journal.pgen.1003264. Publisher: Public Library of Science.

[96] S. R. Searle. Estimating multivariate variance and covariance components using quadratic and bilinear forms. Biometrical Journal, 21(4):389–398, 1979. ISSN 03233847, 15214036. doi: 10.1002/bimj.4710210407. URL https://onlinelibrary.wiley.com/doi/10.1002/bimj.4710210407.

[97] Robert B. Davies. Algorithm AS 155: The distribution of a linear combination of 2 random variables. Journal of the Royal Statistical Society. Series C (Applied Statistics), 29(3):323–333, 1980. ISSN 0035-9254. doi: 10.2307/2346911. URL https://www.jstor.org/stable/2346911. Publisher: [Wiley, Royal Statistical Society].

[98] R. A. Fisher. Statistical Methods for Research Workers. Biological Monographs and Manuals. Oliver and Boyd, 5 edition, 1925. ISBN 0-05-002170-2. URL http://www.haghish.com/resources/materials/Statistical_Methods_for_Research_Workers.pdf.

[99] Daniel J. Wilson. The harmonic mean p-value for combining dependent tests. Proceedings of the National Academy of Sciences, 116(4):1195–1200, 2019. doi: 10.1073/pnas.1814092116. URL https://www.pnas.org/doi/10.1073/pnas.1814092116. Publisher: Proceedings of the National Academy of Sciences.

[100] Paul R. Burton, David G. Clayton, Lon R. Cardon, Nick Craddock, Panos Deloukas, Audrey Duncanson, Dominic P. Kwiatkowski, Mark I. McCarthy, Willem H. Ouwehand, Nilesh J. Samani, John A. Todd, Peter Donnelly, Jeffrey C. Barrett, Paul R. Burton, Dan Davison, Peter Donnelly, Doug Easton, David Evans, Hin-Tak Leung, Jonathan L. Marchini, Andrew P. Morris, Chris C. A. Spencer, Martin D. Tobin, Lon R. Cardon, David G. Clayton, Antony P. Attwood, James P. Boorman, Barbara Cant, Ursula Everson, Judith M. Hussey, Jennifer D. Jolley, Alexandra S. Knight, Kerstin Koch, Elizabeth Meech, Sarah Nutland, Christopher V. Prowse, Helen E. Stevens, Niall C. Taylor, Graham R. Walters, Neil M. Walker, Nicholas A. Watkins, Thilo Winzer, John A. Todd, Willem H. Ouwehand, Richard W. Jones, Wendy L. McArdle, Susan M. Ring, David P. Strachan, Marcus Pembrey, Gerome Breen, David St Clair, Sian Caesar, Katherine Gordon-Smith, Lisa Jones, Christine Fraser, Elaine K. Green, Detelina Grozeva, Marian L. Hamshere, Peter A. Holmans, Ian R. Jones, George Kirov, Valentina Moskvina, Ivan Nikolov, Michael C. O’Donovan, Michael J. Owen, Nick Craddock, David A. Collier, Amanda Elkin, Anne Farmer, Richard Williamson, Peter McGuffin, Allan H. Young, I. Nicol Ferrier, Stephen G. Ball, Anthony J. Balmforth, Jennifer H. Barrett, D. Timothy Bishop, Mark M. Iles, Azhar Maqbool, Nadira Yuldasheva, Alistair S. Hall, Peter S. Braund, Paul R. Burton, Richard J. Dixon, Massimo Mangino, Suzanne Stevens, Martin D. Tobin, John R. Thompson, Nilesh J. Samani, Francesca Bredin, Mark Tremelling, Miles Parkes, Hazel Drummond, Charles W. Lees, Elaine R. Nimmo, Jack Satsangi, Sheila A. Fisher, Alastair Forbes, Cathryn M. Lewis, Clive M. Onnie, Natalie J. Prescott, Jeremy Sanderson, Christopher G. Mathew, Jamie Barbour, M. Khalid Mohiuddin, Catherine E. Todhunter, John C. Mansfield, Tariq Ahmad, Fraser R. Cummings, Derek P. Jewell, John Webster, Morris J. Brown, David G. Clayton, G. Mark Lathrop, John Connell, Anna Dominiczak, Nilesh J. Samani, Carolina A. Braga Marcano, Beverley Burke, Richard Dobson, Johannie Gungadoo, Kate L. Lee, Patricia B. Munroe, Stephen J. Newhouse, Abiodun Onipinla, Chris Wallace, Mingzhan Xue, Mark Caulfield, Martin Farrall, Anne Barton, The Biologics in RA Genetics and Genomics (BRAGGS), Ian N. Bruce, Hannah Donovan, Steve Eyre, Paul D. Gilbert, Samantha L. Hider, Anne M. Hinks, Sally L. John, Catherine Potter, Alan J. Silman, Deborah P. M. Symmons, Wendy Thomson, Jane Worthington, David G. Clayton, David B. Dunger, Sarah Nutland, Helen E. Stevens, Neil M. Walker, Barry Widmer, John A. Todd, Timothy M. Frayling, Rachel M. Freathy, Hana Lango, John R. B. Perry, Beverley M. Shields, Michael N. Weedon, Andrew T. Hattersley, Graham A. Hitman, Mark Walker, Kate S. Elliott, Christopher J. Groves, Cecilia M. Lindgren, Nigel W. Rayner, Nicholas J. Timpson, Eleftheria Zeggini, Mark I. McCarthy, Melanie Newport, Giorgio Sirugo, Emily Lyons, Fredrik Vannberg, Adrian V. S. Hill, Linda A. Bradbury, Claire Farrar, Jennifer J. Pointon, Paul Wordsworth, Matthew A. Brown, Jayne A. Franklyn, Joanne M. Heward, Matthew J. Simmonds, Stephen C. L. Gough, Sheila Seal, Breast Cancer Susceptibility Collaboration (UK), Michael R. Stratton, Nazneen Rahman, Maria Ban, An Goris, Stephen J. Sawcer, Alastair Compston, David Conway, Muminatou Jallow, Melanie Newport, Giorgio Sirugo, Kirk A. Rockett, Dominic P. Kwiatkowski, Suzannah J. Bumpstead, Amy Chaney, Kate Downes, Mohammed J. R. Ghori, Rhian Gwilliam, Sarah E. Hunt, Michael Inouye, Andrew Keniry, Emma King, Ralph McGinnis, Simon Potter, Rathi Ravindrarajah, Pamela Whittaker, Claire Widden, David Withers, Panos Deloukas, Hin-Tak Leung, Sarah Nutland, Helen E. Stevens, Neil M. Walker, John A. Todd, Doug Easton, David G. Clayton, Paul R. Burton, Martin D. Tobin, Jeffrey C. Barrett, David Evans, Andrew P. Morris, Lon R. Cardon, Niall J. Cardin, Dan Davison, Teresa Ferreira, Joanne Pereira-Gale, Ingileif B. Hallgrimsdóttir, Bryan N. Howie, Jonathan L. Marchini, Chris C. A. Spencer, Zhan Su, Yik Ying Teo, Damjan Vukcevic, Peter Donnelly, David Bentley, Matthew A. Brown, Lon R. Cardon, Mark Caulfield, David G. Clayton, Alistair Compston, Nick Craddock, Panos Deloukas, Peter Donnelly, Martin Farrall, Stephen C. L. Gough, Alistair S. Hall, Andrew T. Hattersley, Adrian V. S. Hill, Dominic P. Kwiatkowski, Christopher G. Mathew, Mark I. McCarthy, Willem H. Ouwehand, Miles Parkes, Marcus Pembrey, Nazneen Rahman, Nilesh J. Samani, Michael R. Stratton, John A. Todd, Jane Worthington, The Wellcome Trust Case Control Consortium, Management Committee, Data and Analysis Committee, UK Blood Services and University of Cambridge Controls, 1958 Birth Cohort Controls, Bipolar Disorder, Coronary Artery Disease, Crohn’s Disease, Hypertension, Rheumatoid Arthritis, Type 1 Diabetes, Type 2 Diabetes, Tuberculosis, Ankylosing Spondylitis, Autoimmune Thyroid Disease, Breast Cancer, Multiple Sclerosis, Gambian Controls, Genotyping DNA, Data QC and Informatics, Statistics, and Primary Investigators. Genome-wide association study of 14,000 cases of seven common diseases and 3,000 shared controls. Nature, 447(7145):661–678, June 2007. ISSN 1476-4687. doi: 10.1038/nature05911.

[101] Karla F. Castro-Ochoa, Idaira M. Guerrero-Fonseca, and Michael Schnoor. Hematopoietic cell-specific lyn substrate (HCLS1 or HS1): A versatile actin-binding protein in leukocytes. Journal of Leukocyte Biology, 105(5):881–890, 2019. ISSN 1938-3673. doi: 10.1002/JLB.MR0618-212R.

[102] Obtaining and loading phenotype annotations from the international mouse phenotyping consortium (IMPC) database. In Obtaining and Loading Phenotype Annotations from the International Mouse Phenotyping Consortium (IMPC) Database. Mouse Genome Informatics and the International Mouse Phenotyping Consortium, 2014. URL http://www.informatics.jax.org/reference/J:211773.

[103] Marc H. G. P. Raaijmakers, Siddhartha Mukherjee, Shangqin Guo, Siyi Zhang, Tatsuya Kobayashi, Jesse A. Schoonmaker, Benjamin L. Ebert, Fatima Al-Shahrour, Robert P. Hasserjian, Edward O. Scadden, Zinmar Aung, Marc Matza, Matthias Merkenschlager, Charles Lin, Johanna M. Rommens, and David T. Scadden. Bone progenitor dysfunction induces myelodysplasia and secondary leukaemia. Nature, 464(7290):852–857, 2010. ISSN 1476-4687. doi: 10.1038/nature08851. URL https://www.nature.com/articles/nature08851. Number: 7290 Publisher: Nature Publishing Group.

[104] Jill M. Norris and Stephen S. Rich. Genetics of glucose homeostasis. Arteriosclerosis, Thrombosis, and Vascular Biology, 32(9):2091–2096, 2012. doi: 10.1161/ATVBAHA.112.255463. URL https://www.ahajournals.org/doi/10.1161/ATVBAHA.112.255463. Publisher: American Heart Association.

[105] Juan Carlos Souto, Laura Almasy, Montserrat Borrell, Francisco Blanco-Vaca, José Mateo, José Manuel Soria, Inma Coll, Rosa Felices, William Stone, Jordi Fontcuberta, and John Blangero. Genetic susceptibility to thrombosis and its relationship to physiological risk factors: The GAIT study. The American Journal of Human Genetics, 67(6):1452–1459, 2000. ISSN 0002-9297. doi: 10.1086/316903. URL https://www.sciencedirect.com/science/article/pii/S0002929707632145.

[106] Judith A Blake, Richard Baldarelli, James A Kadin, Joel E Richardson, Cynthia L Smith, and Carol J Bult. Mouse genome database (MGD): Knowledgebase for mouse–human comparative biology. Nucleic Acids Research, 49:D981–D987, 2020. ISSN 0305-1048. doi: 10.1093/nar/gkaa1083. URL https://www.ncbi.nlm.nih.gov/pmc/articles/PMC7779030/.

[107] Cynthia L. Smith and Janan T. Eppig. The mammalian phenotype ontology: enabling robust annotation and comparative analysis. Wiley Interdisciplinary Reviews. Systems Biology and Medicine, 1(3):390–399, 2009. ISSN 1939-005X. doi: 10.1002/wsbm.44.

[108] Xiang Zhang, Shunping Huang, Fei Zou, and Wei Wang. TEAM: Efficient two-locus epistasis tests in human genome-wide association study. Bioinformatics, 26(12):i217–i227, June 2010. ISSN 1367-4803. doi: 10.1093/bioinformatics/btq186.

[109] Jakub Pecanka and Marianne A. Jonker. Two-Stage Testing for Epistasis: Screening and Verification. In Ka-Chun Wong, editor, Epistasis: Methods and Protocols, Methods in Molecular Biology, pages 69–92. Springer US, New York, NY, 2021. ISBN 978-1-07-160947-7. doi: 10.1007/978-1-0716-0947-7_6.

[110] Huanhuan Zhu and Xiang Zhou. Statistical methods for SNP heritability estimation and partition: A review. Computational and Structural Biotechnology Journal, 18:1557–1568, January 2020. ISSN 2001-0370. doi: 10.1016/j.csbj.2020.06.011.

[111] Sang Hong Lee, Naomi R. Wray, Michael E. Goddard, and Peter M. Visscher. Estimating Missing Heritability for Disease from Genome-wide Association Studies. The American Journal of Human Genetics, 88(3):294–305, March 2011. ISSN 0002-9297, 1537-6605. doi: 10.1016/j.ajhg.2011.02.002.

[112] David Golan, Eric S. Lander, and Saharon Rosset. Measuring missing heritability: Inferring the contribution of common variants. Proceedings of the National Academy of Sciences, 111(49): E5272–E5281, December 2014. doi: 10.1073/pnas.1419064111.

[113] G A Churchill and R W Doerge. Naive Application of Permutation Testing Leads to Inflated Type I Error Rates. Genetics, 178(1):609–610, January 2008. ISSN 1943-2631. doi: 10.1534/genetics.107.074609.

[114] Ali Pazokitoroudi, Yue Wu, Kathryn S. Burch, Kangcheng Hou, Aaron Zhou, Bogdan Pasaniuc, and Sriram Sankararaman. Efficient variance components analysis across millions of genomes. Nature Communications, 11(1):4020, 2020. ISSN 2041-1723. doi: 10.1038/s41467-020-17576-9. URL https://www.nature.com/articles/s41467-020-17576-9. Number: 1 Publisher: Nature Publishing Group.

[115] Yue Wu and Sriram Sankararaman. A scalable estimator of SNP heritability for biobank-scale data. Bioinformatics, 34(13):i187–i194, 2018. ISSN 1367-4803. doi: 10.1093/bioinformatics/bty253. URL https://doi.org/10.1093/bioinformatics/bty253.

[116] Matthew Kerin and Jonathan Marchini. A non-linear regression method for estimation of gene–environment heritability. Bioinformatics, 36(24):5632–5639, December 2020. ISSN 1367-4803. doi: 10.1093/bioinformatics/btaa1079.

[117] Carrie Zhu, Matthew J. Ming, Jared M. Cole, Mark Kirkpatrick, and Arbel Harpak. Amplification is the Primary Mode of Gene-by-Sex Interaction in Complex Human Traits, May 2022.

[118] Malachy Campbell, Harkamal Walia, and Gota Morota. Utilizing random regression models for genomic prediction of a longitudinal trait derived from high-throughput phenotyping. Plant Direct, 2(9):e00080, 2018. ISSN 2475-4455. doi: 10.1002/pld3.80.

[119] Eva K. F. Chan, Heather C. Rowe, Jason A. Corwin, Bindu Joseph, and Daniel J. Kliebenstein. Combining Genome-Wide Association Mapping and Transcriptional Networks to Identify Novel Genes Controlling Glucosinolates in Arabidopsis thaliana. PLOS Biology, 9(8):e1001125, August 2011. ISSN 1545-7885. doi: 10.1371/journal.pbio.1001125.

[120] Elizabeth M. Demmings, Brigette R. Williams, Cheng-Ruei Lee, Paola Barba, Shanshan Yang, Chin-Feng Hwang, Bruce I. Reisch, Daniel H. Chitwood, and Jason P. Londo. Quantitative Trait Locus Analysis of Leaf Morphology Indicates Conserved Shape Loci in Grapevine. Frontiers in Plant Science, 10, 2019. ISSN 1664-462X.

[121] Seyoon Ko, Christopher A. German, Aubrey Jensen, Judong Shen, Anran Wang, Devan V. Mehrotra, Yan V. Sun, Janet S. Sinsheimer, Hua Zhou, and Jin J. Zhou. GWAS of longitudinal trajectories at biobank scale. The American Journal of Human Genetics, 109(3):433–445, March 2022. ISSN 0002-9297. doi: 10.1016/j.ajhg.2022.01.018.

[122] Weimiao Wu, Zhong Wang, Ke Xu, Xinyu Zhang, Amei Amei, Joel Gelernter, Hongyu Zhao, Amy C Justice, and Zuoheng Wang. Retrospective Association Analysis of Longitudinal Binary Traits Identifies Important Loci and Pathways in Cocaine Use. Genetics, 213(4):1225–1236, December 2019. ISSN 1943-2631. doi: 10.1534/genetics.119.302598.

[123] Alexessander Couto Alves, N. Maneka G. De Silva, Ville Karhunen, Ulla Sovio, Shikta Das, H. Rob Taal, Nicole M. Warrington, Alexandra M. Lewin, Marika Kaakinen, Diana L. Cousminer, Elisabeth Thiering, Nicholas J. Timpson, Tom A. Bond, Estelle Lowry, Christopher D. Brown, Xavier Estivill, Virpi Lindi, Jonathan P. Bradfield, Frank Geller, Doug Speed, Lachlan J. M. Coin, Marie Loh, Sheila J. Barton, Lawrence J. Beilin, Hans Bisgaard, Klaus Bønnelykke, Rohia Alili, Ida J. Hatoum, Katharina Schramm, Rufus Cartwright, Marie-Aline Charles, Vincenzo Salerno, Karine Clément, Annique A. J. Claringbould, BIOS Consortium, Cornelia M. van Duijn, Elena Moltchanova, Johan G. Eriksson, Cathy Elks, Bjarke Feenstra, Claudia Flexeder, Stephen Franks, Timothy M. Frayling, Rachel M. Freathy, Paul Elliott, Elisabeth Widén, Hakon Hakonarson, Andrew T. Hattersley, Alina Rodriguez, Marco Banterle, Joachim Heinrich, Barbara Heude, John W. Holloway, Albert Hofman, Elina Hyppönen, Hazel Inskip, Lee M. Kaplan, Asa K. Hedman, Esa Läärä, Holger Prokisch, Harald Grallert, Timo A. Lakka, Debbie A. Lawlor, Mads Melbye, Tarunveer S. Ahluwalia, Marcella Marinelli, Iona Y. Millwood, Lyle J. Palmer, Craig E. Pennell, John R. Perry, Susan M. Ring, Markku J. Savolainen, Fernando Rivadeneira, Marie Standl, Jordi Sunyer, Carla M. T. Tiesler, Andre G. Uitterlinden, William Schierding, Justin M. O’Sullivan, Inga Prokopenko, Karl-Heinz Herzig, George Davey Smith, Paul O’Reilly, Janine F. Felix, Jessica L. Buxton, Alexandra I. F. Blakemore, Ken K. Ong, Vincent W. V. Jaddoe, Struan F. A. Grant, Sylvain Sebert, Mark I. McCarthy, Marjo-Riitta Järvelin, and EARLY GROWTH GENETICS (EGG) CONSORTIUM. GWAS on longitudinal growth traits reveals different genetic factors influencing infant, child, and adult BMI. Science Advances, 5(9):eaaw3095, September 2019. doi: 10.1126/sciadv.aaw3095.

[124] Diana L. Cousminer, Yadav Wagley, James A. Pippin, Ahmed Elhakeem, Gregory P. Way, Matthew C. Pahl, Shana E. McCormack, Alessandra Chesi, Jonathan A. Mitchell, Joseph M. Kindler, Denis Baird, April Hartley, Laura Howe, Heidi J. Kalkwarf, Joan M. Lappe, Sumei Lu, Michelle E. Leonard, Matthew E. Johnson, Hakon Hakonarson, Vicente Gilsanz, John A. Shepherd, Sharon E. Oberfield, Casey S. Greene, Andrea Kelly, Deborah A. Lawlor, Benjamin F. Voight, Andrew D. Wells, Babette S. Zemel, Kurt D. Hankenson, and Struan F. A. Grant. Genomewide association study implicates novel loci and reveals candidate effector genes for longitudinal pediatric bone accrual. Genome Biology, 22(1):1, January 2021. ISSN 1474-760X. doi: 10.1186/s13059-020-02207-9.

[125] Joachim Weischenfeldt, Inge Damgaard, David Bryder, Kim Theilgaard-Mönch, Lina A. Thoren, Finn Cilius Nielsen, Sten Eirik W. Jacobsen, Claus Nerlov, and Bo Torben Porse. NMD is essential for hematopoietic stem and progenitor cells and for eliminating by-products of programmed DNA rearrangements. Genes & Development, 22(10):1381–1396, 2008. ISSN 0890-9369, 1549-5477. doi: 10.1101/gad.468808. URL http://genesdev.cshlp.org/content/22/10/1381. Company: Cold Spring Harbor Laboratory Press Distributor: Cold Spring Harbor Laboratory Press Institution: Cold Spring Harbor Laboratory Press Label: Cold Spring Harbor Laboratory Press Publisher: Cold Spring Harbor Lab.

[126] Kevin Berendse, Maxim Boek, Marion Gijbels, Nicole N. Van der Wel, Femke C. Klouwer, Marius A. van den Bergh-Weerman, Abhijit Babaji Shinde, Rob Ofman, Bwee Tien Poll-The, Sander M. Houten, Myriam Baes, Ronald J. A. Wanders, and Hans R. Waterham. Liver disease predominates in a mouse model for mild human zellweger spectrum disorder. Biochimica et Biophysica Acta (BBA) - Molecular Basis of Disease, 1865(10):2774–2787, 2019. ISSN 0925-4439. doi: 10.1016/j.bbadis.2019.06.013. URL https://www.sciencedirect.com/science/article/pii/S0925443919302121.

[127] Patricia Isnard, Nathalie Coré, Philippe Naquet, and Malek Djabali. Altered lymphoid development in mice deficient for the mAF4 proto-oncogene. Blood, 96(2):705–710, 2000. ISSN 0006-4971. doi: 10.1182/blood.V96.2.705. URL https://doi.org/10.1182/blood.V96.2.705.

[128] Xiaoming Feng, Gregory C. Ippolito, Lifeng Tian, Karla Wiehagen, Soyoung Oh, Arivazhagan Sambandam, Jessica Willen, Ralph M. Bunte, Shanna D. Maika, June V. Harriss, Andrew J. Caton, Avinash Bhandoola, Philip W. Tucker, and Hui Hu. Foxp1 is an essential transcriptional regulator for the generation of quiescent naive t cells during thymocyte development. Blood, 115(3):510–518, 2010. ISSN 0006-4971. doi: 10.1182/blood-2009-07-232694. URL https://doi.org/10.1182/blood-2009-07-232694.

[129] Hui Hu, Bin Wang, Madhuri Borde, Julie Nardone, Shan Maika, Laura Allred, Philip W. Tucker, and Anjana Rao. Foxp1 is an essential transcriptional regulator of b cell development. Nature Immunology, 7(8):819–826, 2006. ISSN 1529-2916. doi: 10.1038/ni1358. URL https://www.nature.com/articles/ni1358. Number: 8 Publisher: Nature Publishing Group.

[130] Monika Mortensen, Elizabeth J. Soilleux, Gordana Djordjevic, Rebecca Tripp, Michael Lutteropp, Elham Sadighi-Akha, Amanda J. Stranks, Julie Glanville, Samantha Knight, Sten-Eirik W. Jacobsen, Kamil R. Kranc, and Anna Katharina Simon. The autophagy protein atg7 is essential for hematopoietic stem cell maintenance. Journal of Experimental Medicine, 208(3):455–467, 2011. ISSN 0022-1007. doi: 10.1084/jem.20101145. URL https://doi.org/10.1084/jem.20101145.

[131] Alexander J. Clarke, Ursula Ellinghaus, Andrea Cortini, Amanda Stranks, Anna Katharina Simon, Marina Botto, and Timothy J. Vyse. Autophagy is activated in systemic lupus erythematosus and required for plasmablast development. Annals of the Rheumatic Diseases, 74(5):912–920, 2015. ISSN 0003-4967, 1468-2060. doi: 10.1136/annrheumdis-2013-204343. URL https://ard.bmj.com/content/74/5/912. Publisher: BMJ Publishing Group Ltd Section: Basic and translational research.

[132] Yanzhong Yang, Kevin M. McBride, Sean Hensley, Yue Lu, Frederic Chedin, and Mark T. Bedford. Arginine methylation facilitates the recruitment of TOP3b to chromatin to prevent r loop accumulation. Molecular Cell, 53(3):484–497, 2014. ISSN 1097-2765. doi: 10.1016/j.molcel.2014.01.011. URL https://www.sciencedirect.com/science/article/pii/S1097276514000434.

[133] Feng Xu, Judianne Davis, Michael Hoos, and William E. Van Nostrand. Mutation of the kunitz-type proteinase inhibitor domain in the amyloid *β*-protein precursor abolishes its anti-thrombotic properties in vivo. Thrombosis Research, 155:58–64, 2017. ISSN 0049-3848. doi: 10.1016/j.thromres.2017.05.003. URL https://www.sciencedirect.com/science/article/pii/S0049384817303110.

[134] Keiko Nakayama, Shigetsugu Hatakeyama, Shun-ichiro Maruyama, Akira Kikuchi, Kazunori Onoé, Robert A. Good, and Keiichi I. Nakayama. Impaired degradation of inhibitory subunit of NF-*κ*b (I*kappa*B) and *β*-catenin as a result of targeted disruption of the *β*-TrCP1 gene. Proceedings of the National Academy of Sciences, 100(15):8752–8757, 2003. doi: 10.1073/pnas.1133216100. URL https://www.pnas.org/doi/full/10.1073/pnas.1133216100. Publisher: Proceedings of the National Academy of Sciences.

[135] Sagiv Shifman, Jordana Tzenova Bell, Richard R. Copley, Martin S. Taylor, Robert W. Williams, Richard Mott, and Jonathan Flint. A high-resolution single nucleotide polymorphism genetic map of the mouse genome. PLOS Biology, 4(12):e395, 2006. ISSN 1545-7885. doi: 10.1371/journal.pbio.0040395. URL https://journals.plos.org/plosbiology/article?id=10.1371/journal.pbio.0040395. Publisher: Public Library of Science.

[136] Paulino Perez and Gustavo de los Campos. Genome-wide regression and prediction with the bglr statistical package. Genetics, 198(2):483–495, 2014.

[137] P. Duchesne and P. Lafaye de Micheaux. Computing the distribution of quadratic forms: Further comparisons between the liu-tang-zhang approximation and exact methods. Computational Statistics and Data Analysis, 54:858–862, 2010.

[138] D Kuonen. Miscellanea. Saddlepoint approximations for distributions of quadratic forms in normal variables. Biometrika, 86(4):929–935, 1999. ISSN 0006-3444. doi: 10.1093/biomet/86.4.929. URL https://doi.org/10.1093/biomet/86.4.929.

[139] F. E. Satterthwaite. An Approximate Distribution of Estimates of Variance Components. Biometrics Bulletin, 2(6):110–114, 1946. ISSN 0099-4987. doi: 10.2307/3002019.

[140] Michael Dewey. metap: meta-analysis of significance values, 2022. R package version 1.8.

[141] Daniel J. Wilson. The harmonic mean p-value for combining dependent tests. Proceedings of the National Academy of Sciences U.S.A., 116(4):1195–1200, 2019. URL https://www.pnas.org/content/116/4/1195.

[142] Daniel J. Wilson. harmonicmeanp: Harmonic Mean p-Values and Model Averaging by Mean Maximum Likelihood, 2019. URL https://CRAN.R-project.org/package=harmonicmeanp. R package version 3.0.

[143] Julian Stamp. Leveraging the Genetic Correlation between Traits Improves the Detection of Epis tasis in Genome-wide Association Studies, 2022. URL https://doi.org/10.7910/DVN/WPFIGU.

